# Junctional Adhesion Molecule (JAM)-C recruitment of Pard3 and drebrin to cell contacts initiates neuron-glia recognition and layer-specific cell sorting in developing cerebella

**DOI:** 10.1101/2024.03.26.586832

**Authors:** Liam P. Hallada, Abbas Shirinifard, David J Solecki

## Abstract

Sorting maturing neurons into distinct layers is critical for brain development, with disruptions leading to neurological disorders and pediatric cancers. Lamination coordinates where, when, and how cells interact, facilitating events that direct migrating neurons to their destined positions within emerging neural networks and control the wiring of connections in functional circuits. While the role of adhesion molecule expression and presentation in driving adhesive recognition during neuronal migration along glial fibers is recognized, the mechanisms by which the spatial arrangement of these molecules on the cell surface dictates adhesive specificity and translates contact-based external cues into intracellular responses like polarization and cytoskeletal organization remain largely unexplored. We used the cerebellar granule neuron (CGN) system to demonstrate that JAM-C receptor cis-binding on the same cell and trans-binding to neighboring cells controls the recruitment of the Pard3 polarity protein and drebrin microtubule-actin crosslinker at CGN to glial adhesion sites, complementing previous studies that showed Pard3 controls JAM-C exocytic surface presentation. Leveraging advanced imaging techniques, specific probes for cell recognition, and analytical methods to dissect adhesion dynamics, our findings reveal: 1) JAM-C cis or trans mutants result in reduced adhesion formation between CGNs and cerebellar glia, 2) these mutants exhibit delayed recruitment of Pard3 at the adhesion sites, and 3) CGNs with JAM-C mutations experience postponed sorting and entry into the cerebellar molecular layer (ML). By developing a conditional system to image adhesion components from two different cells simultaneously, we made it possible to investigate the dynamics of cell recognition on both sides of neuron-glial contacts and the subsequent recruitment of proteins required for CGN migration. This system and an approach that calculates local correlation based on convolution kernels at the cell adhesions site revealed that CGN to CGN JAM recognition preferentially recruits higher levels of Pard3 and drebrin than CGN to glia JAM recognition. The long latency time of CGNs in the inner external germinal layer (EGL) can be attributed to the combined strength of CGN-CGN contacts and the less efficient Pard3 recruitment by CGN-BG contacts, acting as gatekeepers to ML entry. As CGNs eventually transition to glia binding for radial migration, our research demonstrates that establishing permissive JAM-recognition sites on glia via cis and trans interactions of CGN JAM-C serves as a critical temporal checkpoint for sorting at the EGL to ML boundary. This mechanism integrates intrinsic and extrinsic cellular signals, facilitating heterotypic cell sorting into the ML and dictating the precise spatial organization within the cerebellar architecture.

**Graphical Abstract:** 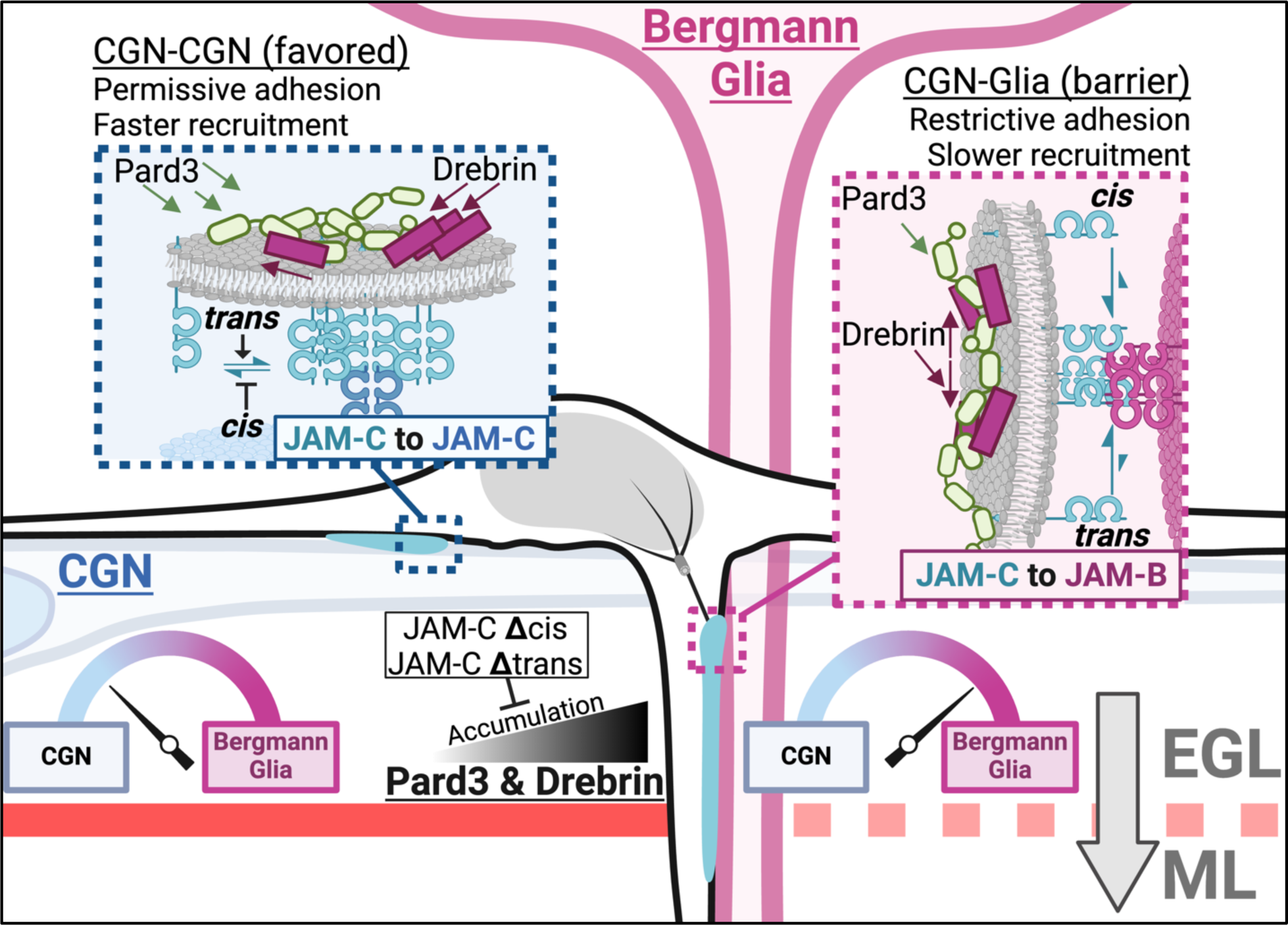

## Introduction

The laminar organization of cells in the brain underlies developmental trajectories and cognitive function. Lamination coordinates where, when, and how cells interact, facilitating the sorting events that guide migrating neurons to their destined positions within emerging neural networks and direct wiring of connections in functional circuits [1]. Identifying molecular mechanisms controlling neuronal migration in the brain, a primary drive underlying lamination, is crucial to understanding how brains develop, and how neuronal positioning goes awry in neurological disorders or pediatric cancer [2]. How neurons navigate the complex environment of axons and cell bodies of neighboring neurons and recognize glial fibers is an area of active investigation, with neuroscientists delving into the intricate and as-yet-unraveled aspects of cellular interactions and how they lead to laminar sorting during development. Significant progress has been made in identifying essential substrates [3–8], guidance systems [9–12], cytoskeletal components [13–21], post-translational modifications [22–26], and how adhesion levels are tuned for neuronal migration [27–33]. However, a comprehensive understanding of how the spatial arrangement and quantity of adhesion molecules on the cell surface regulate adhesive specificity in cell recognition and the translation of contact-based external cues into internal cellular organization remain key knowledge gaps that are challenging to solve. Traditional methods to study adhesion, like electron microscopy, generally lack molecular specificity and have difficulties capturing dynamic processes. Moreover, the field lacks biosensors to directly examine adhesion molecule configuration at cell-cell contacts, and the analytical techniques to define dynamic spatial relationships between adhesive components are still underdeveloped.

Studying the mouse cerebellum’s lamination offers valuable insights into developmental cell recognition mechanisms, particularly through the analysis of cerebellar granule neurons (CGNs), the brain’s most experimentally accessible and numerous neuronal type populating the cerebellar internal granule layer (IGL) [34]. The IGL forms via outside-in development, where cerebellar granule neuron progenitors (GNPs) in the external granule layer (EGL) become postmitotic and then initiate glia-guided radial migration [35]. This well-documented lamination transition involves postmitotic CGNs in the EGL extending parallel fibers in the inner EGL (iEGL) before extending a leading process along Bergmann glia (BG) into the molecular layer (ML) and ultimately migrating along BG fibers towards the IGL [36–38]. Although BG fibers are accessible to GNPs and pre-migratory CGNs in the EGL, mouse CGNs typically spend almost a day post-cell cycle exit interacting with maturing CGNs in the iEGL before effectively engaging with BG fibers for radial migration [36, 39]. The mechanisms that shift CGN contacts to BG fiber recognition and control iEGL residency duration are elusive, largely because integrating assembly of subcellular adhesions with molecular programs that initiate this sorting event of ML entry remains ill-defined.

While many adhesion molecules have been shown to function in the developing cerebellum [32, 40–45], including Astrotactin and N-Cadherin (CDH2) as a CGN-BG receptor pair supporting radial migration [5, 33, 46–48], few provide the defined signaling entry point to attack the challenge of translating spatially arranged contact cues into an internal cellular organization. In contrast, the Junctional Adhesion Molecule (JAM) -B and -C adhesion receptor pair is a tractable starting point to dissect this key feature of cerebellar development. Previous research from our lab showed the Partitioning defective (Pard) polarity protein complex, composed of Pard3, Pard6, and aPKC, is necessary and sufficient to initiate ML entry [32, 49–51]. Several developmental cues interface with the Pard complex in the cerebellum to control CGN migration timing and dynamics [12, 16, 18, 51–53]. Notably, Pard3 induction of EGL exit in development and disease states requires direct binding to the cell adhesion protein JAM-C and is negatively controlled by Siah2 degradation of Pard3 [32, 54]. Our initial work showed that Pard3 provides a predominant control on EGL exit by directing JAM-C exocytosis, so we expect that its interaction with JAM-C initiates glial recognition and migration. However, it’s unclear if 1) the Pard3-JAM-C pathway allows CGNs to distinguish between CGN-CGN or CGN-BG contacts, 2) Pard3 modulates JAM-C spatial arrangement, or 3) JAM-C surface occupancy impacts CGN internal organization. High spatiotemporal resolution achieved by current technical and analytical approaches enabled us in this study to exploit JAM-C as a molecularly defined recognition site to study the recruitment and nucleation of Pard3 to JAM-C adhesions in CGN ML entry.

JAM-A, -B, and -C are Pard3 binding adhesion molecules that are typically studied in tight junction formation, polarity, and migration in non-neuronal cells [55–57]. JAMs belong to the CTX subfamily of the immunoglobulin superfamily, exclusive to vertebrates that homo- or heterotypically adhere to JAMs on the same cell in cis and on opposing cell surfaces in trans [58, 59]. In the developing mouse cerebellum, CGNs express JAM-C, and BG primarily express JAM-B [32, 60]. Structural studies show that JAM proteins possess distinct interaction interfaces that can be mutated to eliminate either cis- or trans-binding, enabling precise manipulation of their spatial arrangement on the cell surface [61–64]. Importantly, altering JAM-B/-C cis- or trans-binding activity allows for experiments testing how JAM-C spatial arrangement contributes to Pard-based mechanisms of how CGNs distinguish between neuronal and BG contacts to address how contact-based external cues are translated into internal cellular organization.

In this work, we tested how JAM recognition connects PARD polarity to glia-guided ML entry by replacing JAM-C with cis- or trans-binding mutants in live, migrating CGNs, in a novel conditional CGN-glial binding assay that allows JAM imaging of both sides of developing adhesions and imaging tissue level CGN sorting in ex vivo cerebellar slices. We showed JAM-C cis- or trans-binding is required for adhesion formation, the cell-intrinsic recruitment of Pard3 or the drebrin cytoskeletal adaptor protein at neuron-glial contacts, and ML entry timing but not oEGL exit for individual CGNs. These results illustrate how JAM-C cell extrinsic recognition through trans binding and regulation by cell-intrinsic cis interactions on seconds to minutes timescales impacts longer-term migration timing and how CGNs discriminate between cellular binding partners during sorting at laminar boundaries.

## Results

### The mutation of JAM-C *cis* and *trans* binding sites reduces CGN adhesion

At the outset of this study, we were interested in dissecting how the spatial arrangement of adhesion molecules on the surface of migrating neurons regulates adhesive specificity in cell recognition. Previous work on JAM-A defines potential cis- and trans-binding interfaces, making the JAM family particularly well-suited to this task. We used point mutations on the extracellular domain of the JAM-C protein to disrupt its cis and trans interactions. Sequence alignment of the cis and trans interaction domains of the JAM proteins shows the conservation of key glutamate residues in mice and humans [Figure 1A]. AlphaFold of the JAM-C monomer was used to visualize the structural conservation of these key residues aligned to the crystal structure of JAM-A (1F97) [65–67]. Zoomed in on the cis [Figure 1B] and trans [Figure 1C] binding sites of JAM-C, each JAM contains structurally conserved arginine to glutamate salt bridges. AlphaFold Multimer structural prediction of four JAM-C subunits reliably recapitulates two independent units of cis binding as previously described crystal structure of JAM-A [Figure 1 – Supplement 1A,C]. Interestingly, upon including 2 subunits of JAM-C and two subunits of JAM-B in the prediction, dimers are preferentially predicted as JAM-C to JAM-B cis heterodimers [Figure 1 – Supplement 1B,D]. This observation is in line with the previous research that showed JAM-B heterodimerization can displace JAM-C homodimers [68].

**Figure 1.**
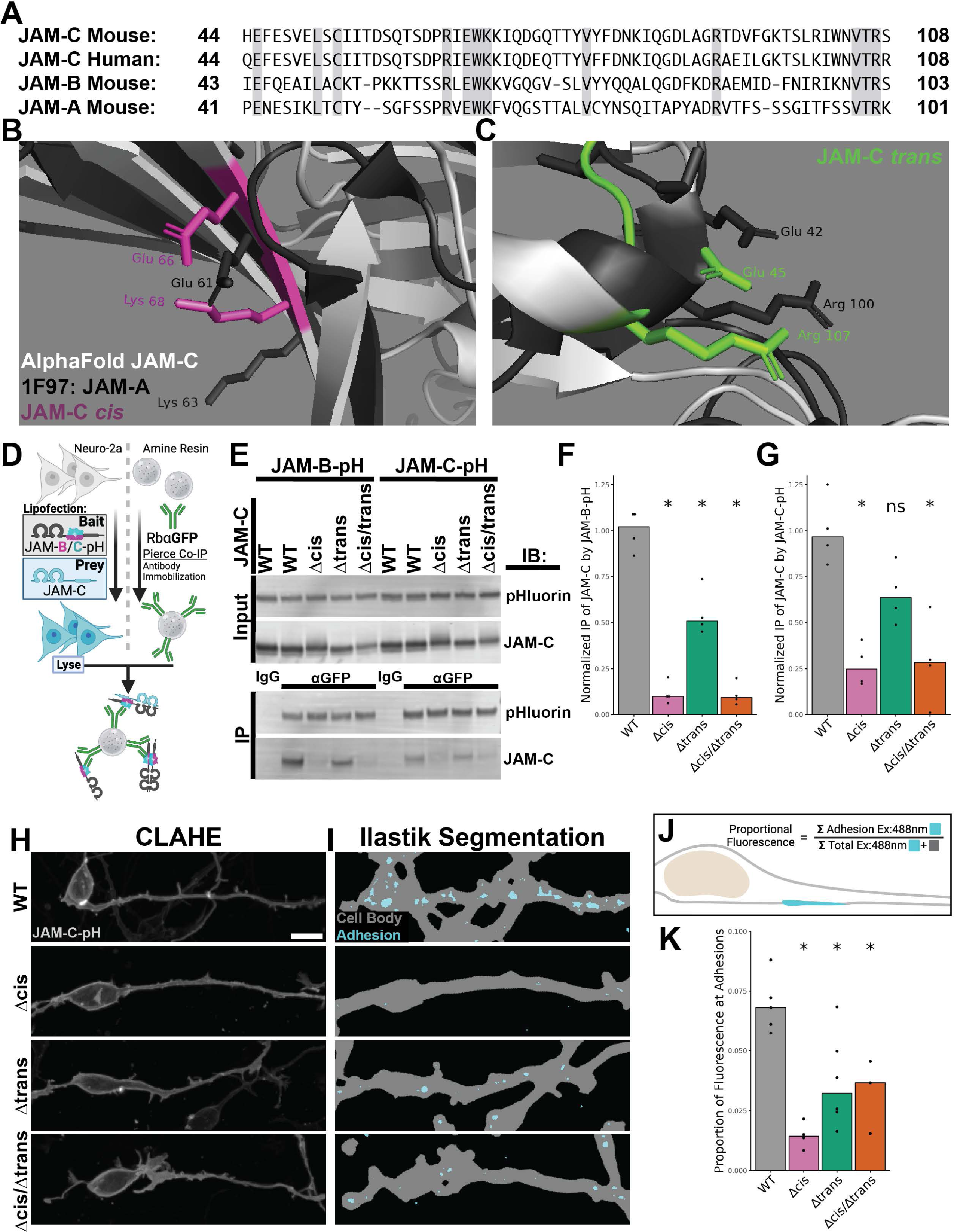
JAM-C *cis* and *trans* binding sites are required for JAM adhesions in CGNs. (**A**) Protein BLAST encompassing the V-Domain of Junctional Adhesion Molecule (JAM)-A, -B, and -C from *Mus musculus* and human JAM-C. Amino acids conserved across all four proteins are highlighted in gray. (**B,C**) AlphaFold prediction of JAM-C protein structure compared to the crystal structure of JAM-A (PDB:1F97) with side chains shown and labeled with one letter amino acid abbreviation for key charged residues that have been previously associated with salt bridges at JAM cis and trans binding sites. JAM-A is represented in black. JAM-C is represented in light gray with the (**B**) cis binding interface highlighted in pink and the (**C**) trans binding interface highlighted in green. (**D-G**) Co-immunoprecipitation (Co-IP) of JAM-C point mutants with JAM-pHluorin. (**D**) Diagram of Rabbit αGFP (Molecular Probes) Co-IP experimental design using JAM-pH as the bait protein with JAM-C prey molecules bearing point mutations expressed in Neuro-2a cell line. (**E**) Representative western blot images of inputs and elutions of JAM-pH immunoblotted with Chicken αGFP (Aves) or Goat αJAM-C (R&D Biosciences). Lanes were loaded in two sets of 5 for JAM-B-pH and JAM-C-pH respectively with negative IP of JAM-pH by Rabbit IgG isotype control (Biolegend) followed by four samples of Rb αGFP CoIP. Quantification of the relative amount of JAM-C bait detected in the IP eluate divided by JAM-C in the input for (**F**) JAM-B-pH and (**G**) JAM-C-pH. The relative amount was normalized to the Co-IP of JAM-C WT for the JAM-pH conditions independently for all biological replicates (N=4) and statistically compared to JAM-C WT using a Wilcoxon Rank Sum test. (**H-K**) Quantification of JAM-C-pH adhesion morphology in JAM-C-replaced CGNs. (**H**) Representative max projection JAM-C-pH images of excitation 488nm fluorescence in the CGNs imaged at 63x 1.46NA and subsequently CLAHE contrast-enhanced to display low signal features. Scale bar = 10 µm (**I**) Representative examples of Ilastik segmentation of cell body in gray and adhesion in cyan for the corresponding fluorescence images. (**J**) Diagram of the quantification for the proportion of total fluorescent signal measured by the adhesion mask divided by the total fluorescent signal measured within the cell body mask. (**K**) Plotted comparison of all biological replicates (N≥3) as daily aggregates of technical replicates that are statistically compared to JAM-C-pH WT using a Wilcoxon Rank Sum test. Statistical significance is represented by standard convention: ns p>0.05, * p<0.05, ** p<0.01, ***p <0.001, and **** p<0.0001.

After identifying potential interaction sites, we aimed to verify that changing key receptor residues affects binding in cis and trans. Focusing on neuronal JAM-C, mutations were introduced at key charged interfaces, swapping glutamates at positions 45 and 66 in JAM-C to arginine to generate the constructs Δtrans (E45R), Δcis (E66R), and Δcis/Δtrans (E45/66R). JAM-C Δcis has been used in other experimental systems to reduce homophilic and heterophilic (JAM-B) cis binding interactions [64, 68, 69]. We chose JAM-C Δtrans based on similarity to the JAM-A surface mediating trans interactions [61]. To further incorporate these constructs into our work, pHluorin fusion proteins (JAM-C-pH) were made to measure changes in cell-cell adhesion, and they were made shRNA resistant to replace endogenous JAM-C with mutants for functional experiments [32, 70]. Using lipofection of Neuro-2a cells as a model cell line with neural characteristics, all fusion proteins were deemed stable and appropriately trafficked to the cell membrane as measured by cycloheximide pulse-chase and surface biotinylation respectively [Figure 1 – Figure Supplement 2]. Interestingly, Δcis/Δtrans received significantly more surface biotinylation, suggesting enhanced trafficking to the cell surface.

With our mutants in hand, we tested how the cis and trans-binding interfaces impact JAM interactions via co-immunoprecipitation (Co-IP) to assess the efficacy of our selected point mutations [Figure 1D]. Co-IP of the JAM constructs with JAM-C and JAM-B pHluorin in whole cell lysates from Neuro-2a cells showed that cis and trans mutations reduce both JAM-C to JAM-C and JAM-C to JAM-B binding [Figure 1E-G]. In both contexts, JAM-B-pH and JAM-C-pH exhibited negligible co-IP of Δcis or Δcis/Δtrans JAM-C. Δtrans yielded milder reduction in co-IP, but this is to be expected since it is still capable of significant cis interaction. This relative difference in adhesion aligns with previously proposed models that cis JAM interactions are required for trans interactions [62, 71]. When comparing JAM-C-pH to JAM-B-pH, JAM-B-pH precipitated more JAM-C. This result further supports a model whereby JAM-B heterodimerization is stronger than JAM-C homodimers [68].

After developing tools to alter JAM-C affinity in cis and trans, we assessed the effects on CGN adhesion by replacing endogenous JAM-C with pHluorin-labeled mutants and analyzing adhesion formation with live cell imaging. Imaging CGNs expressing extracellular mutants revealed a marked decrease in JAM-C-pH adhesion when the cis and trans-binding interfaces are altered [Figure 1H]. To quantify this phenotype, we developed a method to measure the equilibrium of total surface protein and the formation and reduction of JAM-C-pH adhesions, adept at handling adhesion diversity by being stable for hours and dynamically changing during CGN migration [Figure 1 – Figure Supplement 3A-C]. Ilastik, a machine learning based image segmentation software package, was used to identify regions of adhesion and cell bodies in CGNs across expression levels [Figure 1I] [72, 73]. By quantifying proportional fluorescence over time at adhesion, proportion of cell area with adhesion, adhesion size, and total cell area adhesion quantity remained stable between 24-34 hours, suggesting a steady state of adhesion [Figure 1 – Figure Supplement 4A-D]. At 24 hours after nucleofection, the relative JAM-C fluorescent signal in adhesions was analyzed to assess adhesion amount [Figure 1J]. All JAM-C mutants displayed a reduction in the pHluorin signal at CGN-CGN contacts. While Δcis, Δtrans, and Δcis/Δtrans receptors had fewer contacts assessed by our segmentation, the reduction of the Δtrans, and Δcis/Δtrans mutants were less prominent, and assay variability diminished statistical significance [Figure 1K]. The reduction in receptor at adhesion coincided with reduced proportion of cell area with adhesion for Δcis and reduced adhesion size for Δcis and Δcis/Δtrans without any changes in total cell area analyzed [Figure 1 – Figure Supplement 5A-C]. Although the Δcis/Δtrans mutant shows a visible reduction in adhesion, its higher surface biotinylation levels [SuppFig 1b] suggest an increased baseline of JAM-C-pH on the surface. This might spuriously concentrate into segmentable adhesions, potentially leading to underestimating the CGNs phenotype. Altogether, these results showed that not only do our cis and trans mutants alter JAM binding but also affect adhesion formation in CGNs.

### Radial migration initiation into the ML requires *cis* and *trans* interactions

Our lab previously established that CGNs depend on JAM-C adhesion for migration, using shRNA to knock down JAM-C and expressing a dominant negative variant that inhibits JAM-C binding to Pard3 [32]. We aimed to determine the effects of JAM-C’s cis and trans interactions on migration within cerebellar slice environments by substituting native JAM-C with its cis and trans mutants. Validation of our ex vivo slice assay system included a new semi-automated platform for CGN tracking in cerebellar slices and confirmation that JAM-C knockdown could be rescued by a shRNA-resistant JAM-C construct. This was essential as we needed to monitor individual CGN migration events due to their asynchronous nature in the EGL and document cell recognition changes beyond the previously manual analyses applied to understand Pard3 or the drebrin cytoskeletal adaptor in migration [16, 32]. To expand this analysis to hundreds of fluorescently labeled CGNs per slice as unbiasedly as possible, we devised a semi-automated protocol using StarDist for nucleus segmentation and Trackmate for migration tracking [74, 75]. This analytical method matched the accuracy of manual ground truth analyses by three independent lab members on the same raw data subset [Figure 2 – Figure Supplement 1].

**Figure 2.**
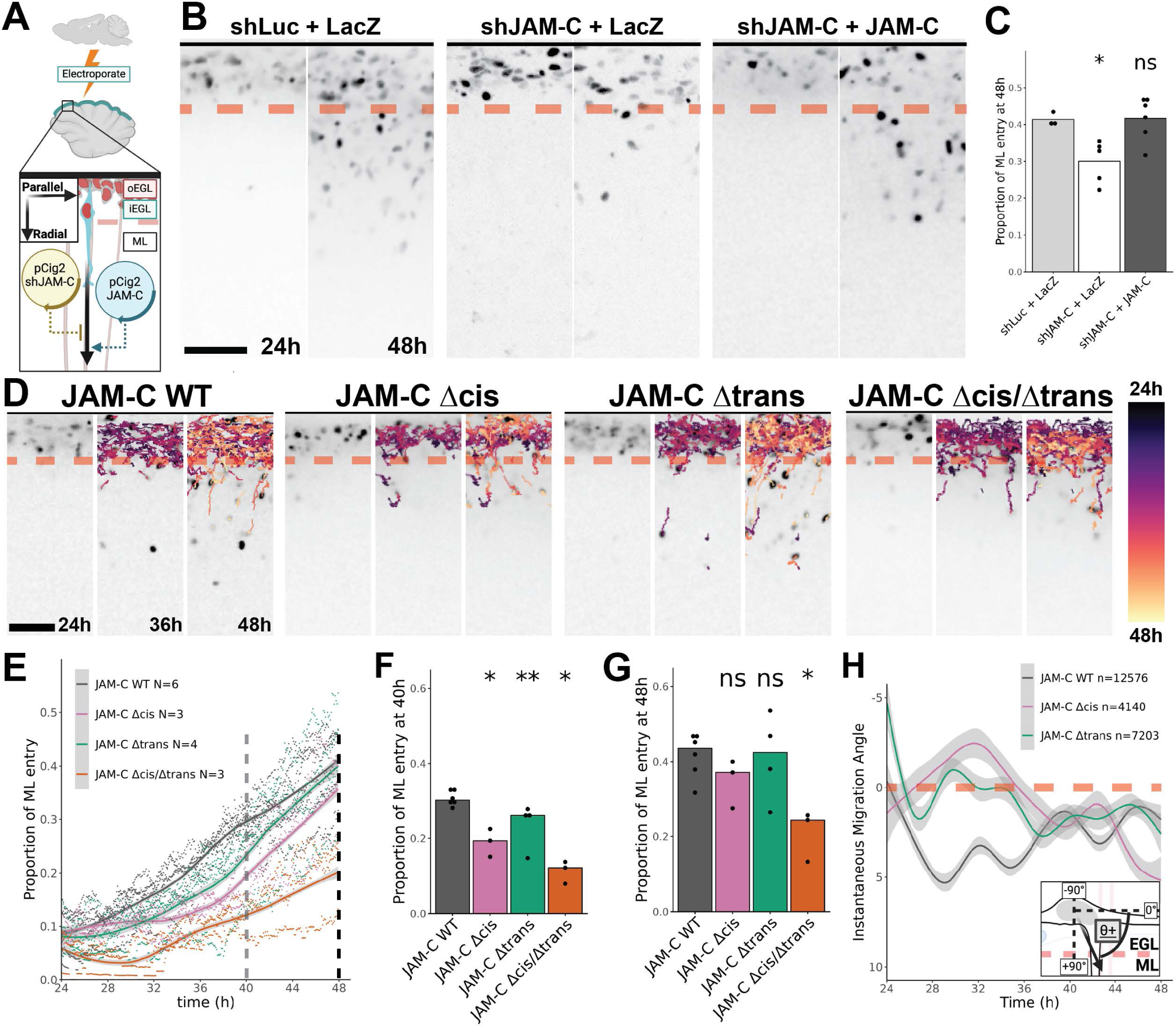
The combination of JAM-C *cis* and *trans* interactions is required to initiate ML entry. (**A-C**) JAM-C replacement rescues JAM-C knockdown in *ex vivo* cerebellar slice cultures. (**A**) Experimental design of JAM-C replacement by electroporating cDNA into the superficial layer of dissected cerebella to knockdown endogenous expression with MiR30 shRNA and express shRNA insensitive JAM-C. (**B**) Representative max projection fluorescent images of Ex 560 H2B-mCherry labeled CGNs at 24 and 48 hours captured with a 20x 0.8 NA objective. The dashed line represents the barrier between the EGL and ML at 50 µm from the cerebellar external edge. (**C**) Quantification of the proportion of all StarDist segmented nuclei further than 50 µm from the slice edge at 48h. Data shown represents single biological replicates (N≥3) that are an aggregate of corresponding technical replicates. Statistical comparison to the shLuc+LacZ positive control for ML entry was performed via Wilcoxon Rank Sum Test. (**D**) Representative tracking data of H2B-mCherry labeled CGNs at 24, 36, and 48 hour. The track color is encoded to represent time point in track based on the provided look up table. The dashed line marks the boundary between EGL and ML at 50 µm from the cerebellum edge. (**E**) Plot of the proportional of H2B-mCherry expressing CGNs >50 µm from the cerebellar edge in the ML. Data points represent independent biological replicates (N) collected at each 5-minute time interval from 24-48h. Data trends over time are plotted as a smoothed Generalized Additive Model (gam). Statistical comparison of proportion of ML entry to JAM-C WT rescue was performed at (**F**) 40 and (**G**) 48 hours of culture using Wilcoxon Rank Sum tests. (**H**) Plot of instantaneous migration angle in relation to the cerebellar edge as a function of time for a subset of the data including JAM-C WT, Δcis, and Δtrans. The trend in time was generated using gam smoothing for the aggregate of all technical and biological replicates (n) of cell motility. Gray shading on the plots represents the 95% confidence intervals of the estimated means. Scale bars = 50 µm. Statistical significance is represented by standard convention: ns p>0.05, * p<0.05, ** p<0.01, *** p<0.001, and **** p<0.0001.

Using our new analysis method, we silenced JAM-C in CGNs with shRNA and introduced shRNA-resistant JAM-C constructs [Figure 2A], then tracked glial-guided migration from 24 to 48 hours by the normalized proportion of CGN nuclei in the ML. JAM-C rescue restored ML entry and IGL-directed migration by 48 hours, showing no notable difference at 24 hours post electroporation [Figure 2B-C, Figure 2 – Video 1], which is before the typical CGN migration onset seen in ex vivo cerebellar slices or in the in vivo mouse cerebellum [16, 32, 36, 52]. By analyzing more than ten thousand individual migration tracks as functions of time and depth in slice, we identified previously unappreciated migration dynamics and their relation to JAM-C function [Figure 2 – Figure Supplement 2E-H]. Silencing JAM-C results in a ML entry deficit that did not affect iEGL occupancy [Figure 2 – Figure Supplement 2A-D], suggesting that its germinal zone exit function occurs at a subsequent stage of CGN maturation. The ML entry delay seen with JAM-C silencing is linked not to altered migration speed but to a change in migration angle [Figure 2 – Figure Supplement 2E,F], suggesting lamina-specific sorting depends on JAM-C adhesion. Supporting a role in laminar sorting, migration speed was unchanged between conditions as a function of depth and time [Figure 2 – Figure Supplement 2E,G], strikingly following trends from previously validated speeds of CGN migration in the EGL (∼0.005 µm/s) towards CGN-BG migration (0.002-0.003 µm/s) [37, 76]. These results not only validate the JAM-C replacement approach and our analysis pipeline but also show that the new careful migration analysis shows that JAM-C loss impacts ML entry, a detail not directly assessed in our previous JAM-C germinal zone (GZ) exit model [32].

Having validated our ex vivo tools, we substituted native JAM-C with its cis and trans variants in the live slice cultures [Figure 2D, Figure 2 – Video 2]. CGNs expressing JAM-C Δcis or JAM-C Δtrans enter the Molecular Layer (ML) significantly slower than those with shRNA-resistant, wild-type JAM-C (Δcis 35% reduction p-value = 0.0238 and Δtrans 13% reduction p-value = 0.0095 at 40 hours post-transfection) [Figure 2E] Interestingly, cells expressing JAM-C Δcis or JAM-C Δtrans ultimately reached similar ML entry levels, indicating that losing just one binding interface delays ML entry. In contrast, CGNs expressing JAM-C Δcis or Δtrans, lacking both binding interfaces, experienced slowed ML entry and never fully occupied the ML. [Figure 2F]. Analysis of migration track dynamics revealed additional insights. Replacing JAM-C with C Δcis or Δtrans did not affect iEGL occupancy. Aggregating migration track data over time revealed that, while migration speed showed minimal difference among JAM constructs, regardless of the layer analyzed [Figure 2 – Figure Supplement 3A,C], CGNs expressing JAM-C Δcis and Δtrans were delayed in instantaneous migration angles into the ML [Figure 2G, Figure 2 – Figure Supplement 3B,D]. The link between migration angle and ML entry, resulting from the loss of JAM-C cis or trans interactions, highlights that CGN recognition and sorting by JAM-C at the EGL/ML boundary are crucial for the observed migration patterns, rather than JAM-C being required for basal motility like those observed when the cytoskeletal components required for migration are inhibited.

### Establishing coculture models to molecularly dissect JAM mediated adhesion

Having found that JAM-C cis and trans interactions modulate ML entry—the lamina where CGNs interact with Bergmann glia—we developed a refined coculture system of CGNs and primary glia. This model allows precise control over adhesion event timing and molecular specificity, facilitating a mechanistic analysis of how these adhesions influence CGN recognition. Distinct from traditional coculture models based on extended coculture and cell interactions, which generate stochastic and unsynchronized adhesion events, our approach of acutely combining specific cell populations to precisely spur adhesion initiation for a detailed examination of adhesion dynamics. The core of this model centers on directly visualizing adhesion molecules on interacting cell surfaces, overcoming the lack of molecular specificity with simple cell aggregation assays or electron microscopy previously employed in older cell recognition studies. Building on our success with pHluorin to visualize JAM-C at cell adhesions, we aimed to define the behavior of cognate JAM receptors by employing a spectrally distinct, pH-sensitive fluorescent protein. To achieve this goal, we labeled JAM-B SNAP using LAMPshade Violet (LSV), a pH-sensitive fluorescent ligand covalently linked to the benzyl guanine SNAPtag ligand, recently developed by Dr. Luke Lavis at Janelia Research Campus [77]. By expressing JAM-C-pH in CGNs and JAM-B-SNAP in glia [Figure 3A], we conducted the first live imaging of molecularly defined CGN-glia adhesion formation using Airy Scan microscopy, chosen for its superior spatial resolution and minimal photobleaching or phototoxicity [Figure 3B-C]. The timelapse images within a representative seven-minute period revealed a swift increase in JAM-C-pH at the CGN growth cone upon glial contact, paralleled by an enrichment of JAM-B-SNAP in the glia. JAM-C-pH fluorescence around the adhesion site decreased from the CGN at this contact point, suggesting lateral recruitment of JAM-C from the neighboring membrane to the developing adhesion.

**Figure 3.**
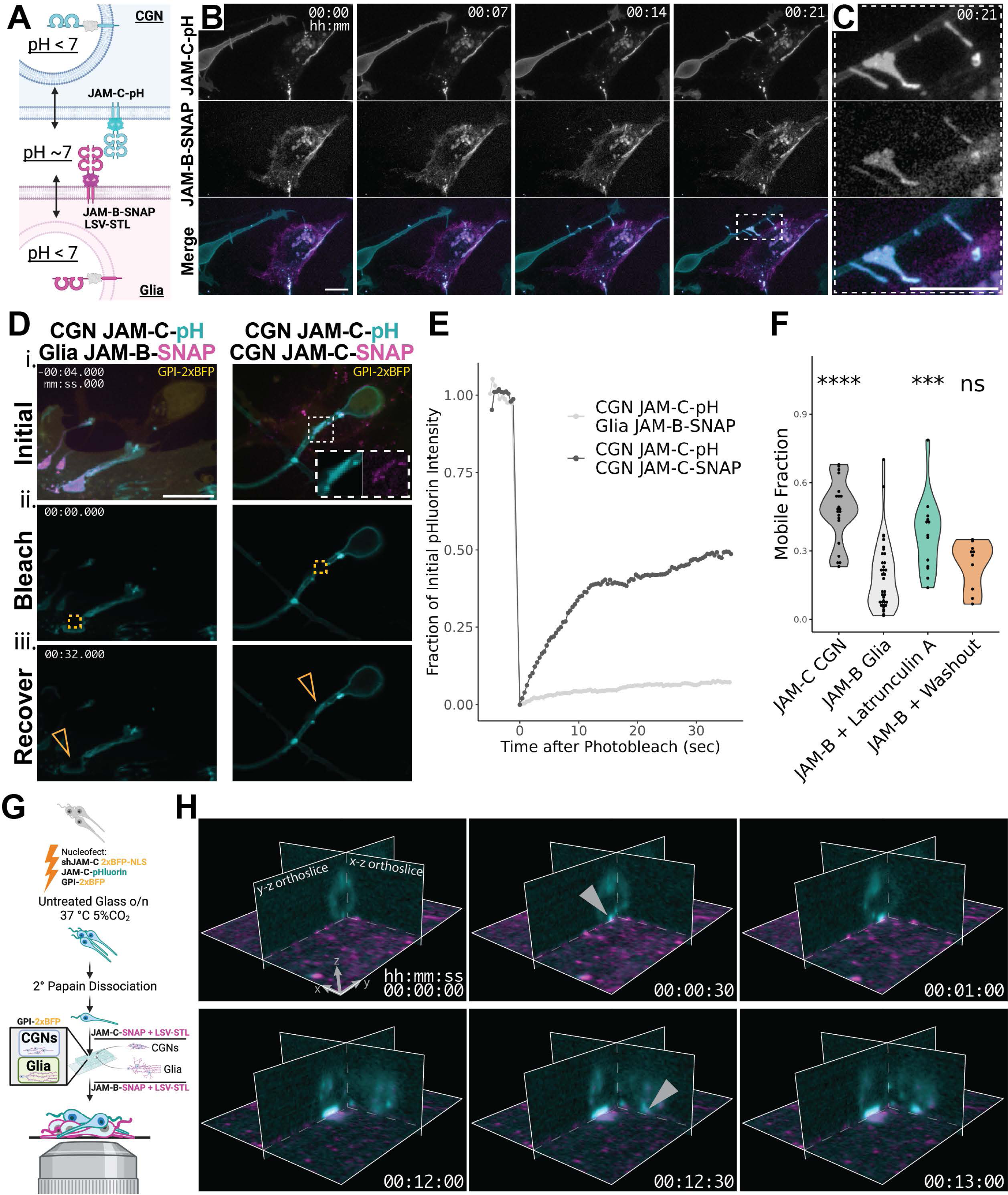
Establishing Coculture Models to Molecularly Dissect JAM Mediated Adhesion. (**A**) Diagram of the conditional detection of JAM specific contacts using two pH sensitive fusion proteins expressed in apposed CGNs and glia: JAM-C-pH and LAMPshade Violet (LSV)-SNAPtag Ligand (-STL). (**B**) Representative time series of contact formation between JAM-C-pH in a CGN and JAM-B-SNAP on a glial cell acquired using AiryScan Microscopy with a 63x 1.4NA objective. Images shown are 0.5 gamma adjusted sum projections of pixel reassigned z-stacks captured every 7 minutes. Channels are shown split into single channel grayscale images and merged in color. Scale bar = 10 µm (**C**) Enlarged view of JAM-C-pH to JAM-B-SNAP contact site. Scale bar = 10 µm (**D**) Representative FRAP images corresponding to: i.) gamma adjusted initial max projections of GPI-2xBFP, JAM-C-pH, and LSV-STL labeled JAM-SNAP multichannel images ii.) JAM-C-pH immediately after photobleach, and iii.) JAM-C-pH at max recovery at 32 seconds. An inset is provided for JAM-C-pH to JAM-C-SNAP to show signal overlap. The orange box in ii.) is the photobleached area, and the orange triangles in iii.) points to local recovery. (**E**) Trace of normalized JAM-C-pH fluorescence for the data shown in (**D**). Statistical significance is represented by standard convention: ns p>0.05, * p<0.05, ** p<0.01, *** p<0.001, and **** p<0.0001. (**F**) Comparison of Mobile Fraction calculated for adhesion to JAM-C on CGNs, JAM-B on glia, JAM-B on glia treated with Latrunculin A, JAM-B on glia after washing away Latrunculin A. Technical replicates from at least 2 biological replicates are plotted as individual points within violin plots. Statistical comparison is performed using Wilcoxon Rank Sum Tests compared to JAM-B on glia. (**G**) Diagram of experimental setup to detect adhesion events for secondarily dissociated JAM-C-pH replaced CGNs plated on monolayers of JAM-C-SNAP expressing CGNs or JAM-B-SNAP replaced glia. (**H**) Representative orthoslice views of CGN JAM-C-pH adhesion to LSV-STL labeled glial JAM-B-SNAP fluorescence acquired by lattice lightsheet microscopy detection through a 50x 1.0NA objective. Orthoslice views of deskewed and deconvolved light sheet timelapses were selected to visualize the cross-sections of two CGNs as they contact JAM-B-SNAP expressing glia at a 30 second intervals. The axial position of the images was manually registered over time, and JAM-B-SNAP fluorescence was photobleach corrected using histogram equalization between time points. Images are gamma adjusted by 0.75 to highlight low fluorescence signal for both JAM-C-pH and JAM-B SNAP. Gray arrowheads are used to point out the observed first instance of JAM-C-pH to JAM-B-SNAP contact formation.

These JAM adhesions, once established, persisted for over 12 hours, extending well beyond the live imaging duration (data not shown). Intrigued by the long-term adhesion lifetimes, we explored JAM-C-pH dynamics at contact sites using fluorescent recovery after photobleach (FRAP) [Figure 3D, Figure 3 – Video 1]. Initial snapshots of SNAP-labeled receptors confirmed CGN JAM-C-pH interactions with JAM-B-SNAP on glia. FRAP at these verified contacts revealed little JAM-C-pH recovery at glial JAM-B-SNAP contacts, while JAM-C pH recovery was rapid when CGNs adhered to other neurons expressing JAM-C-SNAP [Figure 3E]. This slow FRAP recovery of JAM-C-pH diffusion, against the backdrop of adhesion lifetimes spanning hours, indicates a high stability of JAM-glia connections. We hypothesized this contact stability could be due to cortical actin near adhesions because of our prior findings that JAM-C-pH adhesions frequently are coincident with f-actin and adhesion receptor FRAP recovery is limited when actomyosin is stabilized in CGNs, suggesting a critical role of the cytoskeleton in sustaining these connections [17]. Using Latrunculin A, an actin-depolymerizing drug, neuronal JAM-C at CGN-glial contacts became significantly more mobile [Figure 3F], showing the same recovery time as JAM-C at CGN contacts.

We also used lattice light sheet microscopy image growing JAM-C/JAM-B adhesions at the highest speeds, with reduced photobleaching and near isotropic resolution in 3D [Figure 3G]. At this unparalleled level of spatiotemporal resolution, adding dissociated CGNs expressing pHluorin labeled JAM-C onto a glial monolayer expressing JAM-B-SNAP showed JAM-C appeared to preconcentrate on the CGN immediately prior to detectable glial JAM-B recruitment [Figure 3H, Figure 3 – Video 2]. Impressively, detectable JAM-C to JAM-B contact formation occurred on the timescale of seconds and continued growing on the minute scale. These findings highlight the utility of a precisely timed, molecularly defined adhesion system in revealing new insights into CGN-glia adhesion, such as synchronized receptor recruitment upon CGN-glial engagement, lateral JAM-C transfer from adjacent membranes to contact sites, JAM-C pre-clustering prior to glial JAM-B recruitment, and significant restriction of JAM-C diffusion by the actin cytoskeleton at CGN-glial interactions.

### Initiating JAM-C recognition of glial JAM-B requires *cis* and *trans* interactions

Having developed a method to examine cognate JAM receptor interactions between CGNs and glial cells, we then used this approach to explore how JAM-C’s cis- or trans-binding affects CGN-glial recognition. This involved replacing endogenous JAM-C with shRNA-resistant JAM-C pH as wild-type JAM-C or JAM-C mutants cDNAs, like our ex vivo slice experiments strategy. We controlled adhesion timing by adding secondarily dissociated CGNs to monolayers with JAM-SNAP receptors and determined JAM contact prevalence by letting WT or mutant JAM-C-pH CGNs settle for 2 hours on CGN or glial monolayers expressing JAM-C-SNAP or JAM-B-SNAP, respectively [Figure 4A – Figure Supplement 1A]. We then detected overlaps between pH and SNAP receptor species on the surface of admixed cell populations using Ilastik machine learning segmentation [Figure 4B, Figure 4 – Figure Supplement 1B]. The glia with JAM-B-SNAP had more contact formation and larger area per contact overall with JAM-C pH expressing CGNs than monolayers of JAM-C-SNAP CGNs [Figure 4 – Figure Supplement 1C-F]. A significant reduction in overlap between CGN JAM-C-pH and glial JAM-B-SNAP occurred when CGNs expressed JAM-C-pH Δcis, Δtrans, or Δcis/Δtrans mutants, underscoring the essential roles of these interaction interfaces for JAM-C-mediated CGN-glia recognition [Figure 4C]. Notably, there were no significant differences in JAM-C-pH or JAM-SNAP adhesion area or size across conditions. [Figure 4 – Figure Supplement 2]. Conversely, in addition to substantially less SNAP-pH contacts for CGNs, there was no change in overlap between CGN JAM-C-pH Δcis, Δtrans, or Δcis/Δtrans mutants and JAM-C-SNAP on CGNs [Figure 4D]. This unchanged adhesion prevalence on CGNs despite loss of cis- or trans-interfaces suggests *de novo* CGN-CGN interactions are less dependent on JAM-C oligomerization, consistent with our slice experiments where iEGL occupancy, a CGN-rich compartment, was not affected by cis- or trans-mutations. While there was a dramatic reduction in the incidence of JAM-based CGN-Glia adhesion when JAM-C cis or trans interactions were inhibited, in rare cases, adhesions formed when CGNs expressing JAM-C mutants contacted glia. Of these rare instances of JAM-C Δcis, Δtrans, or Δcis/Δtrans CGNs adhesion to JAM-B, the resulting adhesion plaques formed adhesions larger than a CGN soma, similar to the size of adhesions with the wild-type receptor [Figure 4 – Figure Supplement 2D]. Together the results showed that cis and trans interactions of JAM-C influence the probability of JAM-B contact formation.

**Figure 4.**
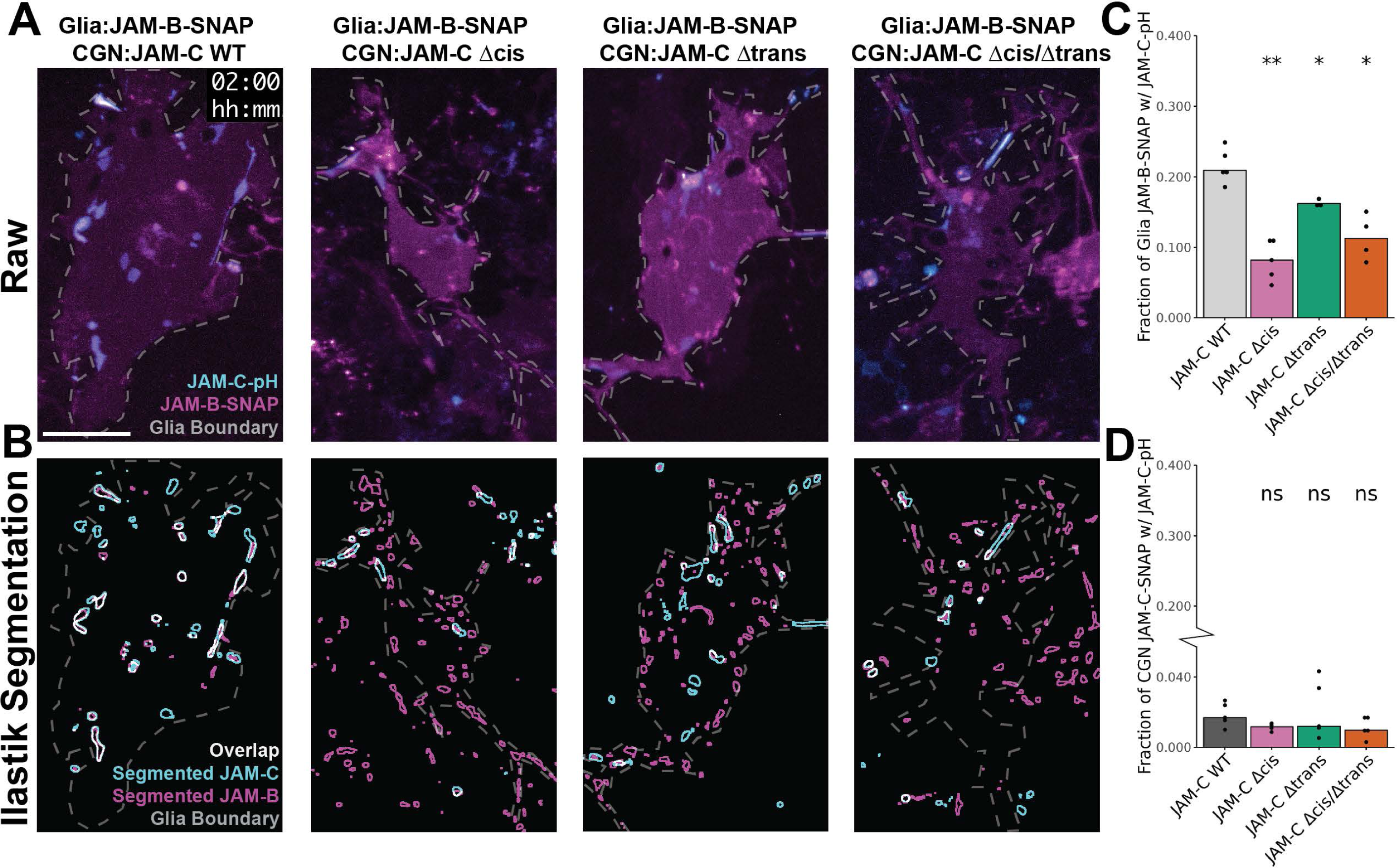
The *cis* and *trans* binding of JAM-C in CGNs initiates recognition of JAM-B on glia. (**C-F**) Evaluation of the prevalence of JAM-C-pH contact formation with LSV-STL labeled JAM-SNAP at 2 hours. (**C**) Representative images of 0.6 gamma adjusted JAM-C-pH fluorescence with 1.2 gamma adjusted JAM-B-SNAP fluorescence for JAM-C-pH WT, Δcis, Δtrans, and Δcis/Δtrans CGNs. Instances of single glia were selected from large fields of view captured by a 20x 0.8NA objective. The gray dashed line is used to manually indicate the edges of a single glia. Scale bar = 50 µm. (**D**) Examples of Ilastik segmentation of JAM-C-pH adhesion and JAM-B-SNAP for the associated fluorescence images. The gray dashed line is used to manually indicate the edges of a single glia as a landmark to compare the fluorescent image. The segmentation is shown as traces of the boundaries for segmented objects. Quantification of adhesion prevalence for (**E**) JAM-B-SNAP on glia and (**F**) JAM-C-SNAP on CGNs. Adhesion prevalence is defined as the area of segmented JAM-C-pH overlapping the segmented JAM-B-SNAP divided by total JAM-B-SNAP segmented area. Data points correspond to the aggregate values of independent biological replicates (N≥3) after excluding outliers. Broken y-axis is used on the CGN graph to show spread of data at low proportion values. Pairwise statistical comparisons are performed using Wilcoxon Rank Sum Tests against JAM-C-WT replaced CGNs. Statistical significance is represented by standard convention: ns p>0.05, * p<0.05, ** p<0.01, *** p<0.001, and **** p<0.0001.

### JAM-C *trans* recognition of JAM-B recruits Pard3 at glial contacts

Although JAM-C mutant CGNs show significantly reduced adhesive recognition of glia, we leveraged the rare events where productive adhesions formed to investigate how cis oligomerization and trans recognition directly influence the recruitment of proteins supporting CGN migration. Previous results from the lab highlighted that Pard3 drove JAM-C exocytosis. However, JAM-C’s role in Pard3 localization in epithelial cells, combined with our findings with the stable CGN-glia contacts and JAM-C depletion on CGNs, suggests a hypothesis that JAM contacts foster a subcellular membrane environment that supports the recruitment of proteins, like Pard3, that is necessary for migration. Indeed, periodic proximity of actin to JAM-C-pH adhesions in the leading process of migrating CGNs, and accumulation of the Siah-regulated actin-microtubule adaptor drebrin suggests this JAM-C environment could also impact cytoskeletal proteins [16–18, 32]. We built upon our JAM recognition assay for studying cognate adhesion receptor recruitment at CGN-glial contacts to measure the impacts of adhesion formation on Pard3 and drebrin organization to sites of cell contact, again taking advantage of the controlled addition of neurons to glia to synchronize the timing of adhesion formation [Figure 5A-B, Figure 5 – Video 1]. We imaged JAM-pH as a reporter for productive adhesion formation and the recruitment of CGN Pard3 and drebrin to the contact site. Similar to our ex vivo slice and neuron-glial contact experiments, JAM-C was silenced in CGNs and JAM-C mutants introduced to assess their impact on Pard3 and drebrin recruitment. We captured unprecedented detail of JAM contact formation for many glia at high spatiotemporal resolution because JAM-B-pH provided a generous photon budget and SoRa spinning disk confocal microscopy enhanced signal brightness due to microlensing [78]. In CGNs expressing WT JAM-C, neuron-glial contacts begin with JAM-pH puncta appearing, aligning with recruited CGN Pard3. Laterally, these expand and merge into multi-micron, clearly defined adhesion compartments containing both species, typically confined to a diffraction-limited axial imaging plane. This observation is the first evidence to our knowledge of a JAM contact influencing Pard3 recruitment at an intercellular contact in neurons. Drebrin showed a complex and heterogeneous relationship with JAM-B-pH, marked by areas of both overlap and exclusion, leading to a ring-like structure around the adhesion at later time points. With a 30-second acquisition rate, the arrival sequence of JAM-B-pH and Pard3 was indiscernible. By increasing temporal resolution to 3 seconds, instances where JAM-B-pH contact preceded Pard3 recruitment were apparent; however, co-recruitment occurred in other cases suggesting temporal resolution will have to be increased to resolve recruitment order fully [Figure 5 – Figure Supplement 1, Figure 5 – Video 2].

**Figure 5.**
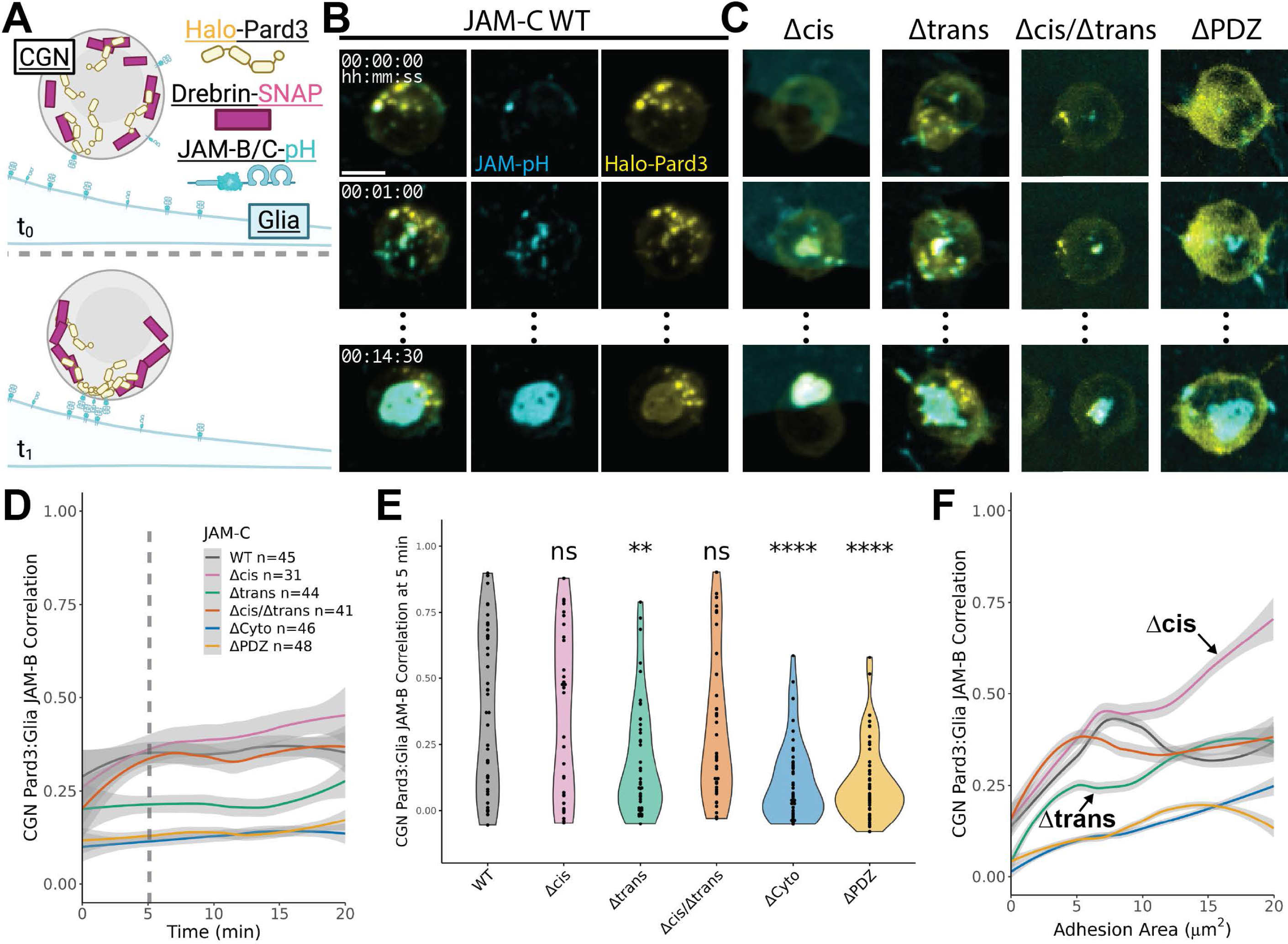
JAM-C recognition of glial JAM-B contacts in *trans* recruits Pard3 in the CGNs. (**A**) Diagram of experimental set up. drebrin-SNAP and Halo-Pard3 are expressed in JAM-C-pH replaced CGNs and Glia express JAM-B-pH at t_0_. We then track the recruitment of these cytoplasmic proteins to the point of contact at t_1_. (**B**) Representative max projection images of JAM-B-pH glial forming a contact with JAM-C WT replaced CGNs that have JFX650-Halo tag ligand (-HTL) labeled Halo-Pard3. Individual channel images of JAM-pH and Halo-Pard3 are presented alongside the multicolor merged image to show the degree of overlapping signal. (**C**) representative multichannel, max projection images of Halo-Pard3 recruitment to JAM contacts for mutant JAM-C replacements that include: Δcis, Δtrans, Δcis/Δtrans, and ΔPDZ which is missing the 6 amino acid C-terminal motif that binds Pard3 (**D**) Loess smoothed plot of Halo-Pard3 recruitment to glial JAM contacts measured by Pearson correlation for individual cells as a function of the time after first adhesion is identified including the Extracellular Domain (ECD) that lacks the entire 48 amino acid cytoplasmic tail in addition to the constructs with associated representative images. The trends are generated from all technical replicates (n) from at least 3 independent biological replicates (N≥3). The dashed line indicates the point in time used for statistical comparison. (**E**) Statistical comparison of Pearson correlation between JAM-pH and Halo-Pard3 at 5 minutes after first sign of adhesion. The Pearson correlation of each technical replicate is plotted within the violin plot. Pairwise statistical significance is calculated in relation to JAM-C WT replaced CGNs using Wilcoxon Rank Sum Tests. (**F**) Loess smoothed plot of Halo-Pard3 recruitment to glial JAM contacts measured by Pearson correlation for individual cells as a function of the segmented adhesion size. The correlation plots and statistical comparisons were generated using all technical replicates (n) from at least 3 independent biological replicates (N≥3) uniformly subsampled at 30 second intervals. Gray shading on the plots represents the 95% confidence intervals of the estimated mean values. Scale bar = 5 µm. Statistical significance is represented by standard convention: ns p>0.05, * p<0.05, ** p<0.01, *** p<0.001, and **** p<0.0001.

We then analyzed Pard3 recruitment in CGNs where mutant receptors replaced JAM-C to identify how cis and trans interactions contribute to intercellular Pard3 recruitment [Figure 5B-C, Figure 5 – Video 5]. Two additional JAM-C mutants were added to our panel: one lacking the PDZ ligand motif (ΔPDZ) at the receptor’s c-terminus that is known to mediate JAM-C interaction with Pard3 [56, 79], and another lacking the 48 amino acid cytoplasmic domain (ΔCyto). We initially assessed the kinetics of glial JAM-B-pH recruitment of CGN Pard3 in JAM-C mutant expressing cells using Pearson’s correlation on a cell-by-cell basis and presented the correlation time courses as Loess smoothed data [Figure 5D]. Interestingly, all three Δcis and Δtrans-JAM-C constructs exhibit a trend toward delayed recruitment over time. Compared to WT JAM-C, JAM-C Δtrans showed decreased Pard3 recruitment at five minutes after initial adhesion [Figure 5E] but caught up by 10 minutes [Figure 5 – Figure supplement 2A], similar to ex vivo slices where this mutant delayed ML entry. When analyzed by adhesion area, JAM-C WT adhesions grew faster than the mutant adhesions, but consistent with the delayed Pard3 recruitment, mutant adhesion growth caught to the WT receptor, and JAM-C Δtrans was the only receptor with significantly smaller adhesions at the end of the time-lapse sequences [Figure 5 – Figure supplement 2B-C]. To identify if the observed variation in Pard3 recruitment was only driven by adhesion size, we plotted proportional recruitment of Pard3 as a function of the JAM-B adhesion area [Figure 5F]. Interestingly, JAM-C Δtrans recruits proportionally less Pard3 as the adhesion grows (4–8µm^2^), indicating the receptor has compromised recruitment and contact formation. In contrast, JAM-C Δcis recruits excess Pard3 when the adhesions get large (16–20µm^2^).

We similarly analyzed drebrin’s recruitment to the JAM contacts [Figure 5 – Figure supplement 3A-C] and detected a low to negative correlation with JAM-B in all scenarios. Pearson’s correlation, treating JAM adhesion sites uniformly, blurred drebrin’s distinct regional correlations and anticorrelations [Figure 5 – Figure supplement 3D]. Acknowledging this limitation of bulk Pearson’s correlation, this approach revealed that JAM-B consistently recruits Pard3 via JAM-C *trans* interactions in the CGNs, with *cis* interactions preventing excess Pard3 recruitment. However, although we anecdotally discerned intriguing spatial relationships between drebrin and growing neuron-glial adhesions, a bulk Pearson’s correlation was insufficient to reveal the complexities or quantitate these relationships.

### Local organization of Pard3 and drebrin at glia contacts requires JAM-C *cis* and *trans* interactions

We created a method to measure local fluorescent signal correlation, bypassing the image inhomogeneity limitations of whole-image spatial analysis to assess the impact of JAM cis or trans interactions on Pard3 and drebrin recruitment. Rather than using Pearson’s correlation on images of individual CGNs, we analyzed local correlation within a sliding window to generate a pixel map for each time point [Figure 6A]. Initial tests ranged from 3×3 pixel (0.618×0.618µm) to 15×15 pixel (3.09×3.09µm) kernel windows [Figure 6 – Figure Supplement 1, Figure 6 – Video 1]. The 3×3 pixel window amplified background noise, while the 15×15 pixel window reduced sensitivity. A 9×9 pixel window (1.85×1.85µm) offered an optimal balance, minimizing noise and maintaining sensitivity. As with the previous bulk correlation analysis, we analyzed the rare adhesions formed by JAM-C mutants to determine adhesion kinetics and binding partner recruitment using our local analysis. Building on the bulk correlation analysis, our local analysis was performed on WT JAM-C adhesions to determine adhesion kinetics and binding partner recruitment for our panel of JAM-C constructs [Figure 6B-C]. Analyzing JAM-C WT CGNs interacting with JAM-B-pH on glia showed that JAM to Pard3 correlation in adhesion formation is a shift from low to positive correlation together accounting for about 90% of the adhesion area [Figure 6 – Figure Supplement 2A-B], whereas JAM:drebrin correlation to JAM adhesion occurs in a concomitant rise of both positive and negative correlation that accounts for about 25% of adhesion area each [Figure 6 – Figure Supplement 2C-D]. The local correlations maps implied the correlation between JAM-B-pH and drebrin tended to have a positive correlation at growing adhesion edges [JAM:drebrin pixel maps in Figure 6B and Figure 6 – Video 2], suggesting drebrin engages adhesions at their margin, similar to the spatial distribution of JAM-C and drebrin we characterized in super-resolution cryo-CLEM [80]. These analyses provide us remarkable access to these subdomains of adhesion formation in a dynamic fashion, enabling us to interrogate the contribution of positive and negative correlation to subcellular recruitment and organization.

**Figure 6.**
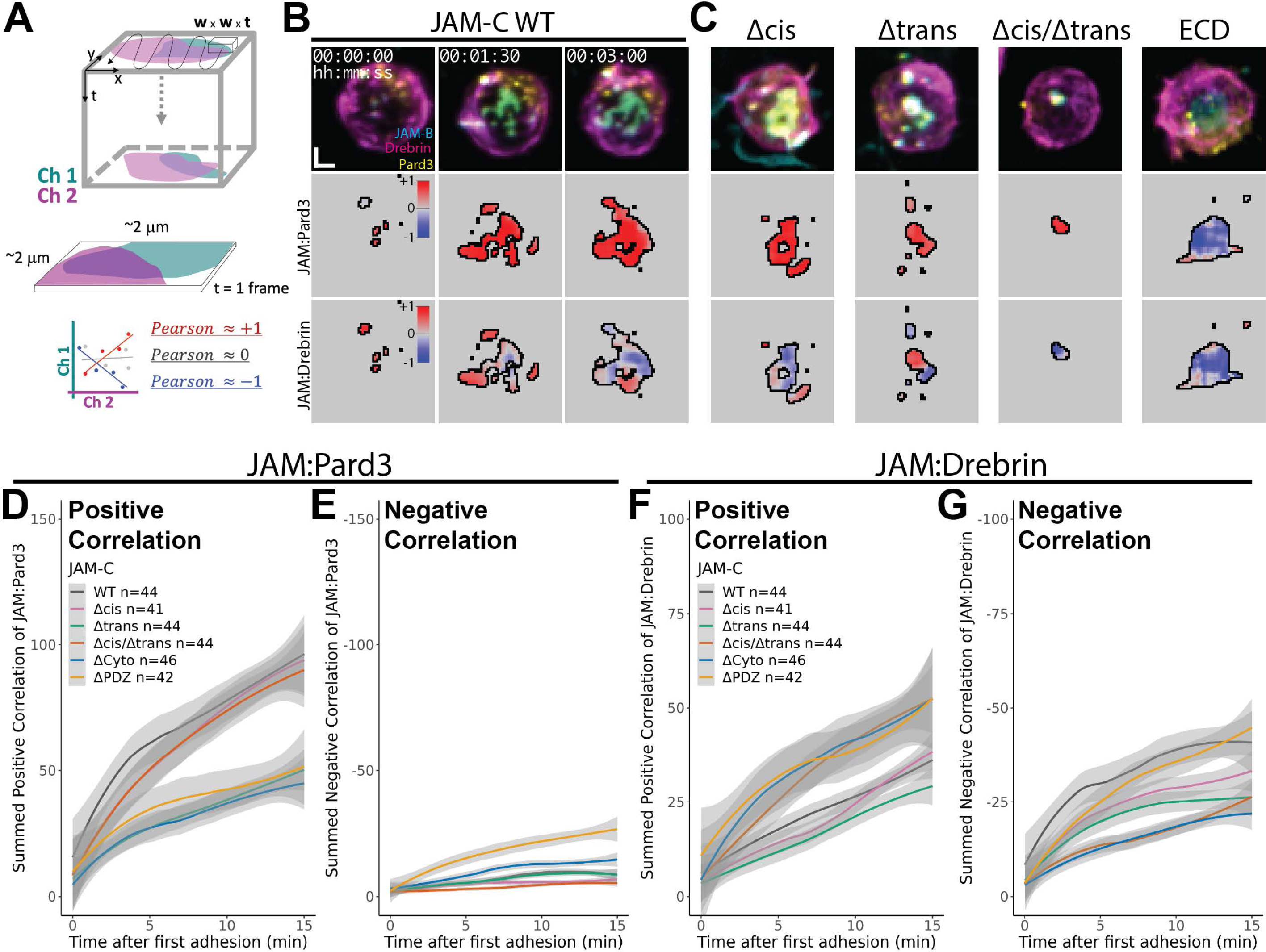
JAM-C *trans* binding is required to locally recruit Pard3 and drebrin to glial contacts. (**A**) Diagram of scheme for analyzing and representing the local correlation of JAM contacts by calculating Pearson correlation in a kernel of defined dimension in x, y, and t that is subsequently scanned across the time lapse image. (**B**) Representative fluorescent data (first row), local recruitment of Halo-Pard3 at JAM contacts (second row), and local recruitment of drebrin-SNAP at JAM contacts (third row) for JAM-C WT replaced CGNs. Pixelwise correlation within segmented JAM-contacts is encoded from blue (correlation = -1) to red (correlation = +1) for paired fluorescence correlation. (**C**) Representative fluorescence images and associated local correlation for the JAM-C mutant constructs: Δcis, Δtrans, Δcis/Δtrans, and ECD that lacks the entire 48 amino acid cytoplasmic tail. Loess smoothed plots of the sum of the (**D**) positive (Pearson > 0) and (**E**) negative (Pearson<0) JAM:Pard3 local correlation as a function of the time after first adhesion. Loess smoothed plots of the sum of the (**F**) positive (Pearson > 0) and (**G**) negative (Pearson<0) JAM:drebrin local correlation as a function of the time after first adhesion. The plots of sum correlation were generated from the combination of technical replicates (n) from at least 3 independent biological replicates (N≥3) uniformly subsampled at 30 second intervals. Gray shading on the plots represents the 95% confidence intervals of the estimated mean values. Scale bar = 2×2 µm.

We next analyzed the sum positive [Pearson>0, Figure 6D, Figure 6 – Figure Supplement 3A] and sum negative [Pearson<0, Figure 6E, Figure 6 – Figure Supplement 3B] local correlation of JAM:Pard3 or JAM:drebrin. Examining extracellular domain mutants, JAM-C Δtrans diminished Pard3’s positive correlation and, consequently, its recruitment as a factor of time post-adhesion initiation and adhesion size. Sum positive [Pearson>0, Figure 6F, Figure 6 – Figure Supplement 3C] and sum negative [Pearson<0, Figure 6G, Figure 6 – Figure Supplement 3D] local correlation of JAM:drebrin enabled us to quantify a relationship of JAM adhesion to drebrin organization that was impossible in our whole image Pearson analysis. Over time and as a function adhesion area, JAM-C WT adhesion significantly enhanced the positive and negative correlation of drebrin at the adhesion site. JAM-C Δtrans trended to reduced drebrin positive correlation with growing CGN-glial adhesions, while in contrast, Δcis/Δtrans led to a consistent increase in drebrin recruitment seen by positive correlation. These findings reveal JAM extracellular domain interactions in CGN-glial adhesions—essential for receptor oligomerization, probability of the glia recognition, and ML laminar sorting—are crucial in recruiting pro-migratory proteins, Pard3 and drebrin, to cell-cell contact sites.

As a complement to the extracellular recognition, investigating cytoplasmic domain mutants highlights cytoplasmic determinants regulating Pard3 and drebrin recruitment at adhesion sites. JAM-C ΔCyto and ΔPDZ exhibited decreased Pard3 recruitment with respect to adhesion initiation time and size [Figure 6D, Figure 6 – Figure Supplement 3A]. Although most JAM-C constructs showed minimal negative correlation, ΔPDZ increased negative JAM:Pard3 correlation, suggesting the cytoplasmic domain not only recruits Pard3 but also controls polarity protein retrieval rates from the membrane [Figure 6E, Figure 6 – Figure Supplement 3B]. Examining JAM:drebrin correlations, JAM-C ΔCyto and ΔPDZ mutants displayed enhanced drebrin recruitment [positive correlation, Figures 6F, Supplement 3C] with ΔCyto additionally reducing negative correlation [Figures 6G, Supplement 3D]. Notably, JAM-C Δcis/Δtrans was the only other receptor to show a similar decrease in negative correlation with drebrin. [Figure 6 – Figure Supplement 4A-I]. Taken together, these results show: 1) Pard3 recruitment at JAM contacts hinges on an intact PDZ-binding motif, 2) ΔPDZ’s enhanced JAM:drebrin correlation reveals Pard3 as crucial to define drebrin recruitment at JAM boundaries, 3) ΔCyto’s positive and negative correlation changes indicate additional boundary definition determinants for drebrin by the rest of the cytoplasmic domain, and 4) the behavior of JAM-C Δcis/Δtrans underscores the extracellular domain’s contribution to drebrin boundary definition [see each behavior highlighted and a summary diagram: Figure 6 – Figure Supplement 4].

Motivated by the anecdotal evidence from correlation maps showing drebrin recruitment at JAM adhesion edges (Figure 5 – Video 1, Figure 6B), and mutant receptor adhesion overgrowing into the drebrin domain, we deepened our analysis to verify if this boundary manifests as measurable patterns of spatial organization. We calculated the spatial distribution of the correlation of fluorescence at adhesion periphery by first analyzing the 0.5 µm edge around JAM adhesions [Figure 6 – Figure Supplement 5A-C, Figure 6 – Video 3] then plotting the ratio of peripheral to whole adhesion values of sum correlation and area [Figures 6, Supplement 5D-G]. The displacement of the correlation sum ratio from the ratio of adhesion area indicates a preferential distribution of correlation. A correlation ratio exceeding the adhesion ratio signifies more correlation in the peripheral 0.5 µm area, and the reverse indicates a central concentration. Interestingly, a higher proportion of positive JAM:drebrin correlation localized to adhesion periphery [Figure 6 – Figure Supplement 5D], which marks the first quantifiable metric of drebrin peripheral distribution during adhesion formation. In contrast, an internal distribution of JAM:drebrin negative correlation and a peripheral distribution of JAM:Pard3 negative correlation were apparent [Figure 6 – Figure Supplement 5E], highlighting regions where these proteins are removed from the adhesion. To evaluate the role of JAM oligomerization and cytoplasmic domain interactions in this boundary condition, this analysis was applied to JAM-C Δcis/Δtrans, ΔCyto, and ΔPDZ mutants [Figure 6 – Figure Supplement 5F-I]. To normalize measurements of distribution against adhesion size between mutant receptors, we subtracted the ratio of peripheral area from respective correlation ratios of JAM:Pard3 and JAM:drebrin [Figure 6 – Figure Supplement 5F]. JAM-C WT and Δcis/Δtrans exhibited even distribution of JAM:Pard3 recruitment, but ΔCyto and ΔPDZ JAM:Pard3 correlation was distributed to adhesion periphery suggesting aberrant adhesion growth of these mutants may indirectly disrupt optimal spatial organization in regards to Pard3 [Figure 6 – Figure Supplement 5G]. The measurements of JAM:drebrin sum correlation distribution revealed the peripheral positive correlation and internal negative correlation of wild-type JAM-C adhesion were both diminished in ΔCyto and Δcis/Δtrans mutants [Figure 6 – Figure Supplement 5H-I]. These findings reveal the intricate interplay between JAM oligomerization, Pard3 function, and cytoskeletal recruitment, establishing a boundary condition crucial for nanoscale adhesion organization that was not discernable without our novel local correlation analyses. Through this complex interaction, CGNs can precisely regulate the segregation and integration of cellular components, ensuring optimal spatial organization and functional efficacy within the adhesion microenvironment.

### Glial JAM contacts are slower at recruiting Pard3 and drebrin, a developmental barrier for CGNs

In our final experiments, we used local correlation analysis to compare CGN-to-CGN and CGN-to-glia interactions in our coculture system, exploring how JAM contacts distinctively recruit adaptor proteins for adhesive recognition in the developing cerebellum in a cell type-specific manner. These studies used local correlation analysis in cocultures for unbiased comparison of Pard3 and drebrin recruitment at CGN-CGN and CGN-glial contacts [Figure 7A-B, Figure 7 – Video 1]. The median of the local correlation values measured within the segmented adhesion area demonstrated that CGNs on a CGN monolayer recruited more Pard3 and drebrin [Figure 7 – Figure Supplement 1A-B]. Despite SNAP-pHluorin glial adhesions enlarging quickly at 2.5 minutes after initial contact and being about twice the size of CGN-CGN adhesions 2 hours later [Figure 7 – Figure Supplement 1D and Figure 4 – Figure Supplement 2F], the adhesion growth between JAM receptors and monolayers exhibited minimal differences [Figure 7 – Figure Supplement 1C] showing that adhesion growth disparities do not explain increased Pard3/drebrin recruitment to CGN-CGN contacts.

**Figure 7.**
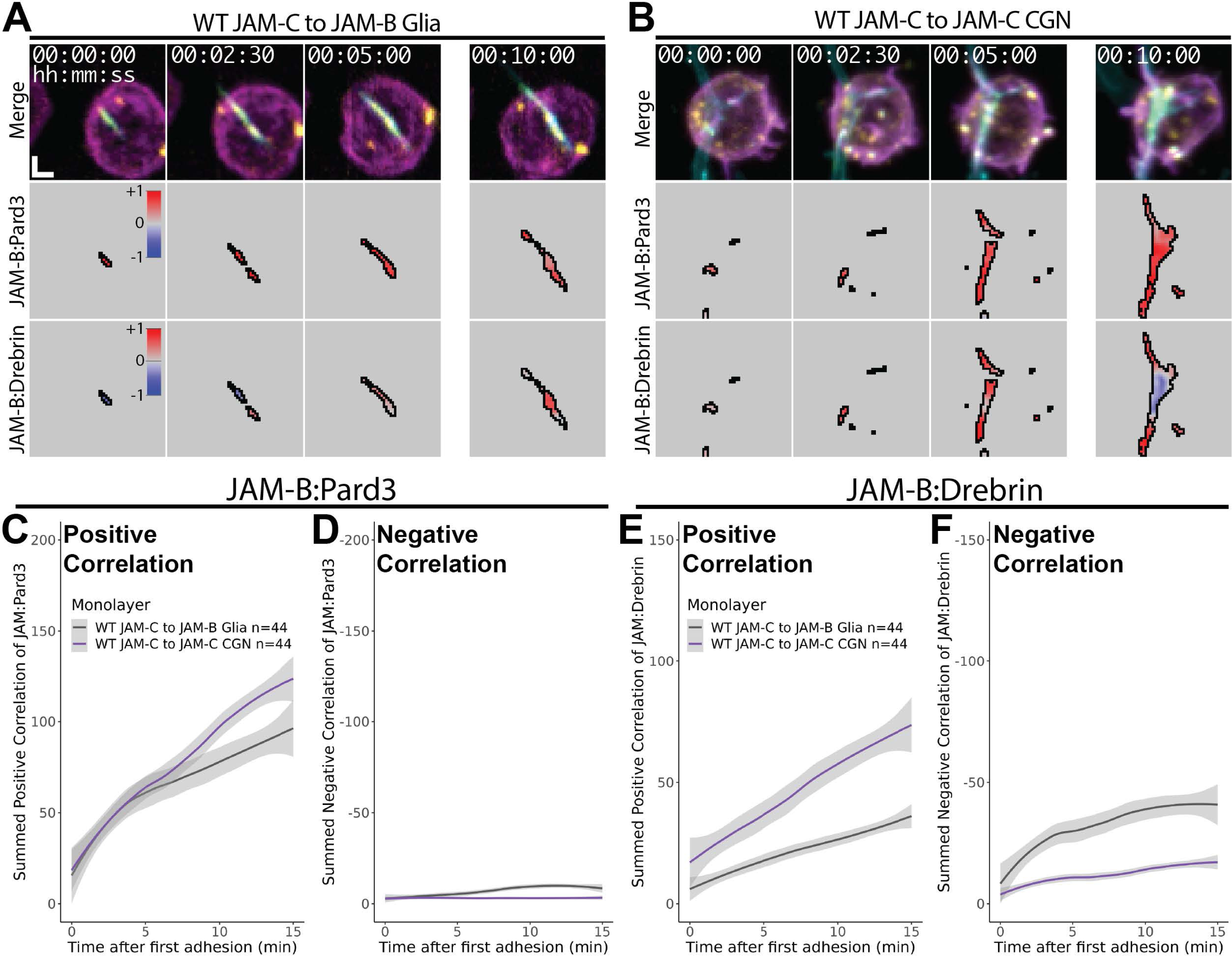
CGN JAM-C recognition of JAM-B on glia is a spatiotemporally defined developmental barrier to the recruitment of Pard3 and drebrin. Representative imaging data of fluorescence and local correlation for drebrin-SNAP, Halo-Pard3, and JAM contacts are shown for (**A**) JAM-B on glia or (**B**) JAM-C on CGNs. Similar adhesion morphology examples were selected for qualitative comparison. (**C-F**) The sum correlation of JAM:Pard3 and JAM:drebrin for JAM-C expressing CGNs is compared for adhesion to either JAM-B on glia or JAM-C on CGNs. Loess smoothed plots of JAM:Pard3 are shown for (**C**) sum positive correlation and (**D**) sum negative correlation as a function of time after first adhesion. Loess smoothed plots of JAM:drebrin are shown for (**E**) sum positive correlation and (**F**) sum negative correlation as a function of time after first adhesion. Loess plots were generated from the combination of all technical replicates (n) indicated in the plot legends. Gray shading on the plots represents the 95% confidence intervals of the estimated mean values. Scale bar = 2×2 µm.

Intrigued by the distinct recruitment profile of CGN-CGN JAM adhesions, we analyzed the contribution of positive and negative correlation domains using sum correlations of JAM:Pard3 and JAM:drebrin [Figure 7C-F, Figure 7 – Figure Supplement 2]. Analysis of JAM:Pard3 positive correlation showed CGN-CGN JAM adhesions exhibited a higher positive correlation of Pard3 over time and by adhesion area [Figure 7C, Figure 7 – Figure Supplement 2A,E], with no significant difference in negative correlation [Figure 7D, Figure 7 – Figure Supplement 2B,F]. Interestingly, this analysis revealed increased JAM:drebrin positive correlation at CGN-CGN adhesions [Figure 7E, Figure 7 – Figure Supplement 2C,G], alongside a significant decrease in negative correlation [Figure 7F, Figure 7 – Figure Supplement 2D,H]. The increase in JAM:drebrin positive correlation and decrease in negative correlation at CGN-CGN JAM adhesions imply that neuronal contacts uniquely boost the recruitment and redistribution of this cytoskeletal partner, a detail overlooked in prior neuronal migration studies [16, 81]. Further extending this concept, analysis of the distribution of positive and negative correlation [Figure 7 – Figure Supplement 3] revealed that CGN-glial JAM adhesion more effectively recruits drebrin at the periphery than CGN-CGN contacts. We first determined the spatial distribution of area to the 0.5 µm peripheral region of the JAM adhesions [Figure 7 – Figure Supplement 3A], and then, with difference from adhesion ratio close to 0 we identified even distribution of Pard3 positive correlation [Figure 7 – Figure Supplement 3B]. Conversely, glial adhesions demonstrated a pronounced peripheral distribution of drebrin positive correlation [Figure 7 – Figure Supplement 3C] and corresponding internal distribution for negative JAM:drebrin correlation [Figure 7 – Figure Supplement 3D]. Unexpectedly, the negative JAM:drebrin recruitment at CGN-CGN contacts even exhibited peripheral distribution. This result suggests that, in contrast to the stability of peripheral drebrin recruitment at the CGN-glia contacts associated with ML entry, the edge of CGN-CGN contacts is engaged in both drebrin recruitment and redistribution. Given that process extension and EGL residency are crucial for CGN maturation, this distinction from glial recognition may allow CGNs to reach a complex morphological state of maturity before migrating into the ML.

Even though JAM receptors appear interchangeable regarding Pard3 and drebrin recruitment, we noted that JAM-C is more likely to spontaneously form adhesions than JAM-B [Figure 7 – Video 2]. Assuming the JAM receptor dichotomy between CGNs and glia acts as an essential laminar sorting signal—specifically, JAM-C in CGNs and JAM-B in glia—we explored whether JAM-B can substitute for JAM-C in silenced CGNs, by tracking migration in live ex vivo cerebellar slice cultures. [Figure 7 – Figure Supplement 4]. Strikingly, replacing CGN JAM-C with JAM-B delayed iEGL entry at 24 hours [Figure 7 – Figure Supplement 4B] and almost completely halted ML entry by 48 hours [Figure 7 – Figure Supplement 4C]. Furthermore, evaluation of the migration speed and angle of CGNs [Figure 7 – Figure Supplement 4D-G] revealed a steady decrease in migration angle over time, favoring parallel migration [Figure 7 – Figure Supplement 4E]. This result indicates that the cell-specific JAM receptor pairs directly regulate laminar sorting into the ML through glial recognition.

Our findings show that CGNs initially favor JAM-C adhesions among themselves, as evidenced by greater recruitment of Pard3 and drebrin at CGN-CGN contacts. However, as CGNs eventually shift towards glial binding in the ML, the less efficient CGN-glial recognition becomes a developmental hurdle CGNs must surpass to enter the ML. Together with our ex vivo slice findings, the diminished cis and trans binding of JAM-C delayed Pard3 recruitment at CGN-glial contacts, causing significant delays in ML entry for CGNs expressing the mutant receptors. This enhances our understanding of the dichotomy between parallel and radial CGN migration in the developing cerebellum. Our findings reveal, for the first time, that JAM recognition of glia serves as a developmental checkpoint coordinating PAR-dependent ML entry.

## Discussion

Understanding how neurons navigate the complex environment of neighboring neurons’ axons, cell bodies, and glial fibers is crucial for deciphering neuronal recognition and connectivity principles. This process, involving interactions between cell adhesion molecules, signaling pathways, and cytoskeletal organization, determines neuronal connection specificity, facilitates accurate positioning by sorting neurons through distinct laminae, and establishes the architecture for functional neural circuits. Our previous work highlighted the essential role of Pard3-dependent JAM-C exocytosis in enabling differentiating CGNs to exit the EGL and initiate radial migration to the inner granular layer (IGL). While increased polarity signaling in CGNs enhanced JAM-C-mediated recognition during radial migration initiation, it remained uncertain whether sorting involves CGN-CGN or CGN-BG contacts or if dissecting JAM-C function ultimately could address the long-standing challenge of how adhesion molecule spatial arrangement influences laminar sorting in neural development. With powerful new imaging modalities, probes for cell-recognition events, and analytical methods dissecting adhesion dynamics, this study dramatically expands our understanding of these problems. It’s now clear JAM-C mediates both CGN-CGN and CGN-BG contacts, with ex vivo slice experiments and coculture studies revealing CGN-BG contacts less efficiently recruit proteins essential for migration, such as Pard3 and drebrin, making them more sensitive to loss of JAM-C cis- or trans-interactions. Indeed, reversing the typical pairing by having CGNs express JAM-B instead of JAM-C, which is less efficient at recruiting Pard3, blocks migration. Seminal research by Fujita fifty years ago found that during CGNs’ 48-hour journey from EGL to IGL, 36 hours were spent in high CGN-CGN contact layers before transitioning to radial migration. This high latency in the iEGL can be attributed to the combined strength of CGN-CGN contacts and the less efficient Pard3 recruitment by CGN-BG contacts, acting as a gatekeeper to ML entry. What distinguishes the glial membrane, making it less conducive to Pard3 and drebrin recruitment in CGN contacts? JAM-B on glial membranes appears more dispersed than JAM-C in CGNs or when glia express JAM-C (See Figure 7, video 2), indicating that the glial membrane’s low JAM oligomerization status contributes to the barrier against forming productive contacts. Our novel capability to dynamically assess adhesion receptor configurations on the glial side of neuron-glial contacts enables the future dissection of unique mechanisms by which glia organize cell surface receptors to non-cell autonomously regulate recognition.

ML entry is a critical developmental transition for CGNs and cerebellar tissue with potential pathological consequences. This study reinforces that CGN maturation status is pivotal in determining ML entry timing, leading to the formation of the IGL, the brain’s largest neuronal compartment. CGNs display a binary migration pattern during normal development, initially migrating parallel to the cell surface within the EGL before transitioning to radial migration, partly because of the favorable CGN-CGN contacts in the iEGL. Previous studies demonstrated that CGNs coordinate ML entry by responding to released guidance molecules like Netrin, Semaphorins, chemokines, or chemotactic neurotrophins, alongside adjustments in their sensitivity to the microenvironmental cues [12, 51–53, 82–88]. These examples highlight mechanisms where CGNs follow putative gradients of external cues to their final positions instead of a binary shift from parallel to radial migration patterns. In contrast, our research shows that CGNs can initiate ML entry by modulating JAM recognition through cis oligomerization and trans recognition of other JAMs. This offers the first evidence that JAM contact formation at the EGL/ML boundary is crucial to instruct ML entry, driven by recognition events rather than the more commonly proposed response to guidance gradients. Indeed, intrinsic decisions such as Pard3 or drebrin recruitment are directly driven by the spatial arrangement of JAM-C at the CGN cell surface in cis and patterned through contact with neighboring cells. Previous models, such as the Astrotactin (ASTN1) to N-cadherin (CDH2) adhesion pair, suggest an expression mechanism, like increased ASTN1 mRNA during CGN differentiation, to facilitate neuron-glial binding changes [46, 48]. However, it remains unclear how these adhesion events are tuned on rapid cell biological timescales—contrasting with the slower mRNA accumulation in differentiation—and their connection to signaling events to essential migration machinery. Our work reveals that JAM contacts between CGNs and their binding partner nucleate Pard3 and drebrin recruitment on rapid time scales, effectively creating a platform that supports migration. As CGNs mature, JAM-C cis- and trans interactions enable CGNs to exchange EGL adhesions to form productive contacts with glia and take a position in the IGL through glial-guided migration. The delays in migration resulting from the loss of JAM-C cis- or trans-binding, significantly lowering adhesion frequency, indicate that once JAM and associated Pard3 or drebrin accumulation at glial contacts surpass a critical threshold—as demonstrated in the rare adhesion events quantified in Figures 6 and 7— sufficient JAM adhesion exists to facilitate movement to the IGL. Alternatively, other glial-guided migration mechanisms, like ASTN1/CDH2, may make up for lower JAM adhesion levels.

This study illustrates that JAM recognition is a post-translational migration checkpoint at glial contacts, regulating Pard3 recruitment and polarization timing via cis and trans interactions. During neuronal differentiation, Zeb1-driven fluctuations in Pard complex mRNA serve as a rheostatic gate globally linking polarization to CGN maturation [50, 89] Conversely, JAM-C mRNA escapes this regulation, making Pard3 recruitment through JAM recognition a mechanism for spatially fine-tuning polarization at the protein level [50]. This recognition event not only focally recruits Pard3 but also peripherally recruits drebrin, a microtubule-actin crosslinker responsible for steering CGN migration direction. Extrinsically, our findings demonstrate that cis regulation of JAM-C recognition and Pard polarity depend on trans interactions to dictate the timing and location of positioning events. Notably, a glycosylation site near the trans interface, previously linked to JAM adhesion, emerges as a potential regulatory mechanism for trans-interactions [90]. Furthermore, neither Δcis nor Δtrans mutants could replicate the local recruitment of Pard3 seen with intact JAM interactions, suggesting that the loss of dominant JAM interactions allows cell-intrinsic Pard polarity and extrinsic secondary heterotypic interactions to prevail. A close parallel to how JAM-C functions in this respect of cell recognition is seen in research demonstrating CDH2’s crucial role in glial recognition and polarity within the murine cerebral cortex [29, 30]. Varied cell-intrinsic mechanisms harmonize CDH2’s surface stabilization and internalization for cell recognition, reminiscent of Pard3’s control over JAM-C exocytosis [27, 31, 91]. CDH2 interprets cell extrinsic information, functioning downstream of surface stabilization in response to the extracellular ligand Reelin, whose homotypic interactions are enhanced by glycosylation, while studies involving the related protein E-Cadherin demonstrate that integrin activation can allosterically regulate adhesion [27, 92–94]. Intriguingly, recent reports indicate that JAMs collaborate with adhesion molecules in non-neuronal cells, with Apex2 proximity ligation experiments revealing JAM-C association with CDH2. This suggests the potential for co-regulation of the CGN migration by JAMs and cadherins, a topic ripe for future exploration [95, 96]. While cadherins engage in low-affinity cis interactions that enhance trans cooperativity, leading to large contact areas averaging around 5 µm^2^ in giant unilamellar vesicles [97], our findings reveal that JAM-C in live CGNs governs glial contact prevalence, with areas often exceeding 20 µm^2^. This underscores the unique contribution of JAM-C to neuronal layer sorting in the brain, highlighting both the similarities and distinct differences from cadherin-mediated mechanisms.

Cryo-CLEM in CGNs revealed JAM-C contacts are flat, EM-dense structures similar to tight junctions as compared to the adherens junctions that are less dense, in part, to accommodate the large extracellular domain of cadherins [80, 98] Local correlation analysis and coculture methods to synchronize cell contacts unveiled novel aspects of these specialized JAM contacts. Pard3 is evenly distributed within JAM contacts, escalating in abundance as adhesions expand. Conversely, drebrin, localized peripherally, maintains a positive correlation with JAM contacts without a significant increase in abundance alongside adhesion growth. The distinct localization of JAM partners underscores their specialized functions: Pard3, crucial for polarity, permeates adhesions to set up a polarity axis, drebrin, by settling at the adhesion margins, underscores the idea that these edges are pivotal for cytoskeletal interaction within JAM-mediated adhesions. Cytoplasmic domain mutants reveal the potential for interplay between Pard3 and drebrin. For instance, the ΔPDZ mutant, unable to bind Pard3, exhibits more JAM-drebrin positive correlation than wild-type JAM-C. This suggests Pard3 may delineate JAM-C contact boundaries, influencing where drebrin is preferentially recruited. Additionally, JAM-C Δcis and Δtrans mutants display enhanced drebrin positive local correlation, indicating a complex relationship where both extracellular and cytoplasmic domains of JAM-C contribute to defining proper adhesion boundaries. Despite the new insights into adhesion formation and partner recruitment, how do perturbing adhesion dynamics on the seconds and minutes time scale seen by Δcis and Δtrans mutants lead to multiple-hour delays to ML entry and migration initiation? CGN migration is cyclical, coordinating adhesion dynamics with cytoskeletal movements across nucleokinesis cycles. Delays in adhesion formation or diminished recruitment of adhesion partners across these cycles can cumulatively delay CGN’s entry into the IGL. How do cerebellar granule neurons (CGNs) navigate the developmental transition from initially favoring CGN-CGN contacts to embracing CGN-glial contacts? The Siah2 ubiquitin ligase could be a critical regulator in this process, negatively influencing germinal zone exit and eventual entry into the molecular layer (ML) by weakening JAM adhesions and targeting Pard3 and drebrin for proteasomal degradation. Our local correlation analysis reveals that JAM adhesions act as scaffolds for Pard3 and drebrin recruitment, establishing them as units regulated by Siah. Consequently, reducing Siah2 expression during differentiation may lift restrictions on these recruitment events, thereby fostering interactions with glial cells. This intricate interplay of molecular interactions and regulations at its heart represents the molecular asymmetries that underly the polarity surrounding adhesions and underscores the complexity of CGN maturation and cell recognition during migration, spotlighting the pivotal roles of Siah2, JAM adhesions, and the orchestrated recruitment and function of Pard3 and drebrin.

The role of JAMs in CGN sorting at the EGL-ML boundary suggests broader implications beyond neural development. Developmental cell sorting models, like the Differential Affinity Hypothesis (DAH) proposed by Steinberg in 1963, highlight cells’ drive to maximize mutual adhesion while minimizing non-adhesive areas [99, 100]. The introduction of the Differential Interfacial Tension Hypothesis (DITH) by Brodland in 2002 offers a framework for understanding ML entry as a sorting event, addressing the complexity of cerebellar development beyond simple adhesion receptor comparisons [101]. In DITH, membrane and cortical cytoskeleton interfacial tension influences sorting by adhesive strength [102, 103], which in the case of the CGN that experiences higher actomyosin contractility and JAM mediated adhesion during differentiation provides a timer by which CGNs can respond at EGL/ML boundary [18, 32]. SHH-driven medulloblastoma cells continuing to reside in their germinal niche may be linked to CGN mis-sorting. This pediatric cancer can be modeled in mice by constitutively de-repressing Gli through the conditional loss of the Patched SHH receptor, keeping CGNs in a progenitor-like state [104, 105]. Interestingly, our previous research demonstrated that knocking down Ras-MAPK or Siah2 ubiquitin ligase, a negative regulator of JAM adhesion [32], reinstates proper CGN sorting into the ML in cKO Ptch1 mice [51]. Reduction of ECM adhesion through integrin β1 in the CGNs subsequently decreased Ras-MAPK signaling [32, 51], suggesting a decrease in ECM adhesion or increase in JAM cell-cell adhesion can restore CGN sorting. The shift from extracellular matrix (ECM) to cell-cell adhesions aligns with the Differential Adhesion Hypothesis (DAH); CGNs transition inwardly in the outer EGL, simultaneously reducing ECM adhesion while enhancing intercellular adhesion. JAM recognition acts as a gating mechanism for CGN sorting into the ML, with CGN-CGN contacts enhancing process extension and Pard3 recruitment to balance cortical tension and adhesion. CGN-glia contacts, facilitated by JAM, enable precise single-cell sorting, shifting the balance towards specific, stable cell adhesions. Once sorted into the ML by JAM recognition, the process transitions to CDH2-driven sorting, consistent with the Differential Adhesion Hypothesis (DAH), allowing for swift movement through the ML to the IGL. Similarly, in Drosophila, the IgSF protein Echinoid, binding to the Pard3 homolog Bazooka, polarizes cortical actin, delineating cell sorting boundaries during embryonic development [106, 107]. We expanded on the IgSF barrier concept, demonstrating that the cis and trans dynamics of JAM contacts serve as checkpoints for individual CGN sorting during cerebellar development. The evolution of IgSF boundaries as a mechanism for tissue sorting presents a promising avenue for future research in disease and development.

### Limitations of this study

At this point in the work, we have not shown *in vivo* data, but the data herein has provided the basis for developing *in vivo* models to validate our observations. While our choices for JAM-C mutations are based on structurally validated interfaces for JAM adhesion, we have not performed crystallographic validation of the change in receptor orientation. While we examined deviation from negative and positive controls throughout this work, we did use shRNA knockdown, which may yield incomplete loss of endogenous protein, and overexpression does not represent physiological protein amount. The cerebellar glia used for cocultures are a heterogeneous population of astroglia that are not restricted to BG. From the analysis perspective, the data reported in this manuscript is mostly 2D projections. While this simplification was sufficient for our conclusions, the full realization of the complexity of our model in 3D will require significant analytical and technological modifications.

## Author Contributions

L.P.H. designed, performed, and analyzed all the experiments and generated the initial draft of the manuscript. A.S. developed the computational tool to calculate local Pearson’s correlation in a Jupyter notebook interface. D.J.S. conceived the idea for the study, performed pilot experiments showing changes in adhesion with early versions of JAM-C-pH mutants, guided project design, and provided a thorough revision of the manuscript. All authors contributed to the editing and revision of the final manuscript.

## Acknowledgements

We would like to thank and acknowledge Christophe Laumonnerie for generating and sharing cDNA constructs for pCig2 MiR-30 shJAM-C 2XBFP-NLS, pCig2 GPI-2xBFP, and pCig2 Halo-Pard3. We thank DJS’s lab manager, Niraj Trivedi for animal colony maintenance and method troubleshooting for standard laboratory procedures. We thank William Bodeen for years of maintaining DJS’s Marianis Spinning Disk Confocal. Thank you to William Bodeen and Christophe for providing the benchmark data to evaluate our approach’s quality to track CGN migration in live cerebellar slices. We thank all members of the Solecki lab, current and past, for advice throughout this work’s development, implementation, and reporting. Finally, a very special thanks to St. Jude Children’s Research Hospital Cell and Tissue Imaging Center Light Microscopy core facility for providing access to and maintaining some of the microscopes used in this work: 2 Marianis Spinning Disk Confocal Microscopes, the Zeiss 980 LSM for AiryScan, and the Zeiss Lattice Lightsheet 7. Of note, current facility director Aaron Taylor, staff Aaron Pitre, and past facility director Victoria Frohlich were instrumental in the training and properly implementing the imaging techniques used in this work. Aaron Pitre provided invaluable resources and hands-on training for FRAP, Airy Scan, and Lattice Lightsheet. All graphical models in this manuscript were generated using a paid subscription of BioRender. LPH has received funding and support from the St. Jude Graduate School of Biomedical Sciences throughout this work. DJS is funded by the American Lebanese Syrian Associated Charities (ALSAC) and by grants 1R01NS066936 and R01NS104029 from the National Institute of Neurological Disorders (NINDS). The content of this manuscript is solely the responsibility of the authors and does not necessarily represent the official views of the NIH.

## Materials Availability Statement

All materials generated in this work are available from DJS upon reasonable request.

## Materials & Methods

### Data reporting

We did not perform any power analyses to determine sample sizes. Experiments were all performed in at least biological triplicate, defined as independent experiments performed on different days labeled as ‘N’. Total replicate values for experiments that are labeled ‘n’ include both technical and biological replicates aggregated as specified. Experiments performed were not randomized, and investigators were not blinded to conditions or outcomes of experiments. Inclusion and exclusion criteria are described in the corresponding method descriptions.

### Animals

All mouse lines were housed and bred in standard conditions (e.g., pathogen free and continuous access to food/water) in accordance with guidelines established and approved by the Institutional Animal Care and Use Committee at St. Jude Children’s Research Hospital (Protocol no. 483). The experiments described here were performed in wild type C57BL6 mice collected between postnatal days 6 and 8 as this is the time when the CGNs in the EGL have reached a maximum population. Equal amounts of male and female mice were used in this experiment, since, to this date, there is no evidence of a significant difference in EGL exit based on animal sex.

### Chemicals and Antibodies

See the Key Resources Table for resource identification, supplier, and concentrations/dilutions used in this work.

### Plasmids Vectors

The sources for cDNA expression vectors used in this work are reported in the Key Resources table. New constructs were commercially synthesized and subcloned into either the pCA LNL or pCig2 plasmid backbones by GenScript (Piscataway, NJ). Subsequent exchange of expression vectors between pCig2 and pCA LNL and other subcloning were performed in house using standard procedures. Donor cDNAs were digested with NEB high-fidelity restriction enzymes for 2 hours at 37 °C. Linear fragments were purified by 1% agarose (Lonza) gel electrophoresis and subsequent QIAquick™ gel extraction (Qiagen). Linear fragments were then re-ligated using the Rapid DNA Dephosphorylation and Ligation Kit (Roche, Millipore Sigma): dephosphorylating the backbone fragment and ligating at room temperature for at least fifteen minutes.

All cDNAs were amplified and stored by transforming Top10 (Thermo Fisher) competent *E. coli*, single colonies selected on Luria Broth (LB) agar (MP Biomedicals) plates containing 100 µg/mL Carbenicillin (Invitrogen), expanded in 100-150 mL of 100 µg/mL Carbenicillin containing LB (RPI) liquid media supplemented with 10% Terrific Broth (TB, RPI), and purified after overnight shaking culture at 37 °C via MaxiPrep (Invitrogen). The list of all cDNAs used in this work, and associated amount used in experiments are included in the Key Resources Table.

### Predicting protein structure and alignment

JAM structures as monomers and multimers were predicted using the CollabFold Jupyter Notebook (https://github.com/sokrypton/ColabFold) and visualized using PyMol (https://pymol.org/2/) [65–67, 108]. Peptide sequences for prediction were submitted as copies of the extracellular domain omitting the associated signal peptides at the N-terminus of JAM-B (Uniprot Accession: Q9JI59, A.A. 29-236) or JAM-C (Uniprot Accession: Q9D8B7, A.A. 35-240) from *Mus musculus*. Displayed structures are the hierarchically assigned best prediction out of 5 simulations. Alignment to the JAM-A crystal structure (1F97) was performed in PyMol through sequence alignment and structural superposition without any further refinement.

### Cell Line

The mouse Neuro-2a cell line was obtained from ATCC and cultured following standard practices. The cells were grown in High Glucose DMEM (Gibco) supplemented with 10% fetal bovine serum (Gibco), 2 mM glutamine (Gibco), 100 U/mL Penicillin, and 100 µg/mL Streptomycin (1:100 Pen Strep Gibco) at 37°C and 5% CO_2_. Cells were passaged two to three times per week at a ratio of one to six or one to ten. Cells were washed once with 1X DPBS and detached from cell culture-treated vented flasks (TPP) with TrypLE dissociation reagent (Gibco).

### Co-Immunoprecipitation

Co-immunoprecipitation of JAM-pH was performed on overexpressed protein produced in Neuro-2a cells. The Neuro-2a cells were transfected with cDNA using Lipofectamine™ 2000 (Invitrogen) based on manufacturer’s instructions. One million Neuro-2a cells were plated on 60 mm tissue-culture treated dishes at a cell density of 250,000/mL and incubated overnight at 37°C and 5% CO_2_ mixed into ambient O_2_ to achieve approximately 50% confluency in high glucose DMEM (Gibco) supplemented with 10% FBS (Gibco), 100 U/mL Penicillin, and 100 µg/mL Streptomycin (1:100 Pen Strep Gibco). The following day, cells were lipofected with a 600 µL formulation containing 12 µL Lipofectamine™ 2000, 2 µg of pCig2 JAM-C-pH or JAM-B-pH bait, and 2 µg of pCig2 JAM-C prey constructs bearing point mutations. Lipofections proceeded overnight to achieve stable expression.

The following day, 90-100% confluent cells were processed based on the Pierce™ Co-Immunoprecipitation Kit. Cells were lysed in 300 µL of Pierce™ IP lysis buffer containing 1X Halt™ Protease and Phosphatase inhibitor (Thermo Scientific) and 25 µL was set aside for SDS-PAGE analysis of protein input. Co-IP resin was prepared by crosslinking 100 µg of Rabbit αGFP (Molecular Probes) or Rabbit IgG isotype (BioLegend) to 60 µL of AminoLink™ Plus Coupling Resin (ThermoFisher) with sodium cyanoborohydride per manufacturers instruction. 275 µL of lysate was applied to 15 µL bed volume of the functionalized Co-IP resin and mixed end-over-end overnight at 4 °C. The flow-through, washes, and elution were all performed in a tabletop microcentrifuge at 1000xg for 1 minute. Each sample was washed 10x with 250 µL of Dulbecco’s PBS at 4 °C as 3 extra washes after NanoDrop A280 nm measured protein concentration reached <10 ng/µL. Elution was performed by incubating the washed resin at room temperature in 40 µL of the acidic Elution Buffer for 10 minutes and subsequent centrifugation, collecting 3 elution samples for each column. 10 µL of 1:5 diluted input and 40 µL of undiluted elution were separated via SDS-PAGE and analyzed by Western Blot as described later.

Protein content was measured in the input and primary elution using Chicken IgY αGFP (Aves) to detect the pHluorin bait and Goat IgG αJAM-C (R&D Bioscience) to detect JAM-C prey. Immunprecipitation was validated based on the ratio of JAM-pH in the Input and elution fractions. Co-IP was calculated as the proportion of the total signal of JAM-C in the elution divided by JAM-C in the input. The ratios of elution:input was normalized to WT JAM-C Co-IP from >4 biological replicates for JAM-pH bait.

### Preparation and Enrichment of primary Cerebellar Granule Neurons (CGNs) and glia

We collected and enriched primary CGNs from postnatal day 6-8 (P6-P8) C57BL6 mice cerebella as previously described [109]. Cerebella were dissected from the rest of the brain and the meningeal layer was carefully removed. The dissected cerebella were homogenized by mincing and papain dissociation the Postnatal Neural Dissociation Kit (Miltenyi Biotec). Homogenate was run through fire-polished glass pipettes to generate a single-cell suspension. The single cell suspension was centrifuged at 2700xg with low deceleration through 35% Percoll layered on top of 60% Percoll to isolate the dense granule cells.

Isolated granule cells were collected in media containing 10% horse serum (Gibco), 2mM Glutamine (Gibco), 100 U/mL Penicillin, and 100 µg/mL Streptomycin (1:100 Pen Strep Gibco) in DMEM-Flourbrite (Gibco), referred to as GCM-FB. The granule cells were then pre-plated on 0.0025% Poly-ornithine (Sigma-Aldrich) treated; cell-culture coated polystyrene dishes (Corning) to remove residual astroglial cells. The CGNs were then used for *in vitro* experiments as described later. Primary astroglia were propagated from the pre-plate using cell-culture media containing 10% FBS, 2mM Glutamine (Gibco), 100 U/mL Penicillin, and 100 µg/mL Streptomycin (1:100 Pen Strep Gibco) in DMEM-Flourbrite after removing the granule cells and incubated at 37 °C and 5% CO_2_ mixed into ambient O_2_. Transfections of primary cells were all performed with the Amaxa Mouse Neuron Nucleofector Kit (Lonza), using the O-005 nucleofection program. Nucleofections in CGNs were performed with 55 µg cDNA formulations on 6,000,000-10,000,000 CGNs. Glia nucleofections were performed with 30 µg cDNA formulations on 400,000-1,000,000 glia grown to confluency that were dissociated with TrypLE (Gibco).

### Adhesion Morphology Analysis in JAM-C-pH replaced CGNs

Analysis of adhesion morphology was performed on live CGNs building on our previously described methods [32]. 8-well #1.5 glass bottom µ-slides (Ibidi) were plasma-cleaned to activate the glass surface, treated with 0.0025% Poly-ornithine in water for 1-2 hours, and subsequently coated with 1 µg/cm^2^ purified mouse laminin (Millipore Sigma) diluted in FB-DMEM at 37 °C for about 1 hour. During this coating, a freshly dissociated single cell suspension of CGNs after pre-plating was used for JAM-C replacement. Between six and ten million CGNs were nucleofected as follows: 15µg pCig2 miR-30 shRNA 2XBFP-NLS directed at *Mus musculus* JAM3 mRNA (UCAAAUUCAUGUACCACUGGG) [53], 10µg pCig2 Cre T2A H2B-mCherry, 10µg GPI-linked 2XBFP for constitutive membrane labeling, and 15µg of miR-30 insensitive JAM-C-pHluorin restricted to Cre expressing cells by the pCA LNL plasmid backbone [110]. Cultures were allowed 24 hours at 37 °C and 5% CO_2_ to achieve stable expression before imaging adhesion morphology in live cells using an environmentally controlled chamber on a Marianis Spinning Disk Confocal Microscopy (Intelligent Imaging Innovation). Images were taken at multiple fields of view for each biological replicate of conditions with a 1.46 NA 63X objective. We acquired Z-stacks at 0.23 µm axial steps across 8-10 µm for both 405nm and 488nm excitation at low power set empirically for each biological replicate to minimize photo damage.

Morphology analysis was performed as deviation from JAM-C-pH WT adhesion. Maximum intensity projections of Z-stacks were generated and exported as tiffs with SlideBook (Intelligent Imaging Innovation). We used Ilastik machine learning image segmentation to independently 1) segment the cell body of healthy cells trained on all conditions for membrane-linked BFP (Ex 405 nm) using pixel classification with object classification and 2) segment disproportionately bright JAM-C-pH adhesions (Ex 488 nm) using pixel classification trained to identify WT adhesion [72, 73]. Analysis of segmentation masks was performed on adhesions segmented within the cell body masks using Analyze Particles in FIJI applied to the raw pHluorin images [111]. Quantification of the csv output from FIJI was performed in RStudio to acquire three key metrics associated with each field of view. 1) Proportional fluorescence in adhesions was calculated by applying segmentation masks to the raw pHluorin data, dividing pHluorin fluorescence in segmented adhesions by the pHluorin fluorescence for the entire cell body. 2) Proportional area was calculated as the sum area of segmented adhesion divided by the total area of segmented cell body. 3) Mean adhesion size was calculated as the average size of the pHluorin-segmented adhesions. 4) Total cell area analyzed was calculated as the sum area of the cell mask. To evaluate reproducibility for day-to-day biological replication in nucleofected cells across expression levels, fields of view were combined as technical replicates to constitute individual biological replicates used in our statistical comparisons. Adhesion morphology over time was analyzed using the same methods described for Z-stacks collected every 15 minutes for 10 hours.

### Cerebellar Slice Cultures

The cerebellar slice culture experiments were performed based on our previously outlined procedure [32]. Cerebella were dissected from postnatal day seven (P7) C57BL6 mice in modified Hank’s balanced salt solution (mHBSS: 2.5mM HEPES, 30mM D-Glucose, 1mM CaCl_2_, 1mM MgSO_4_, and 4 mM NaHCO_3_ in 1X HBSS). The dorsal meningeal layer was carefully removed from the vermis of the cerebella to fully expose lobe VI/VII for electroporation, and the choroid plexus was pulled away from the ventral surface. For the experiments reported in this work we electroporated individual cerebellum with 135µL of 5mg/mL cDNA formulations composed of 1µg/mL pCig2 H2B-mCherry, 3µg/mL of pCig2 MiR-30 2XBFP-NLS shRNA constructs, and 1µg/mL of either unlabeled pCig2 JAM-C constructs or pCig2 LacZ as a control. Isolated cerebella were incubated in the mHBSS formulation containing cDNA at 4 °C for 15 minutes. The cerebella were then electroporated with four 500 ms pulses of 80V interspaced by 50 ms pauses along the dorsoventral axis. Next, electroporated cerebella were embedded in 4% LMP agarose dissolved in mHBSS and then cut into 300 mm sagittal slices using a VT1200 vibratome (Leica). The slices were finally placed on 0.4 mm PTFE cell culture inserts (Millicell) equilibrated in a 6-well plate with serum-free media containing 1x B27 supplement (Gibco), 1x N2 supplement (Gibco), 2mM Glutamine (Gibco), 100 U/mL Penicillin, and 100 µg/mL Streptomycin (1:100 Pen Strep Gibco) in DMEM-Fluorbrite (Gibco), referred to as SFM-FB. The slices were then incubated at 37 °C and 5% CO_2_ mixed into ambient O_2_. Slices were then used for live-imaging.

### Analysis of Neuron migration in Cerebellar Slice Cultures

CGN nuclear migration in live cerebellar slices was evaluated by elaborating on methods we have previously used to quantify migration speed and direction [32]. At about 23 hours of culture, props were cut off the bottom of PTFE inserts and transferred to a large #1.5 glass-bottom dish (Electron Microscopy Sciences) containing 1 mL of warmed SFM-FB. A metal washer was placed on top of the insert rim to stabilize the samples. The chamber was sealed with parafilm and then transferred to an environmentally controlled chamber on a Marianis Spinning Disk microscope (Intelligent Imaging Innovation) to equilibrate before imaging. Single fields of view were selected at the locally optimal region of H2B-mCherry (Excitation 560 nm) fluorescence at the flat region of the 6^th^ lobe for each cerebellar slice using a 20x 0.8NA objective. Nuclear migration was imaged as 50µm stack of Z-slices at 1 µm intervals acquired every 5 minutes from 24-48 hours using minimal laser power empirically set for individual biological replicates to reduce photo damage. Replicates were excluded from analysis if we observed insufficient fluorescence, detachment from the PTFE support, or identified tissue damage.

2D CGN nuclear migration analysis was performed on maximum projections generated in SlideBook (Intelligent Imaging Innovation) from a sub-stack for cerebellar slices that exclude CGNs that have exited from the tissue slice at the interface with the PTFE insert. Subsequent image processing was performed in FIJI. First, lateral displacement during imaging was corrected using the plugin in FIJI for Linear Stack Alignment with SIFT [112]. Next, the distance from surface (DFS) was calculated and added to a new channel using a Euclidean distance transform of a manually specified border that was interpolated across the imaging sequence [113]. Finally, we used StarDist to automatically segment fluorescent nuclei combined with the Linear Assignment Problem (LAP) tracker in Trackmate to follow CGN nuclei migration [74, 75]. CGN nuclei with radii between 2-8µm were tracked if the associated centroid was within the cerebellum (DFS 0-250 µm) and the distance moved over individual 5-minute intervals did not exceed 12 µm (<0.04µm/s). Tracks with >6 consecutive points (>30 min) were included in the analysis, excluding tracks at the edge of the field of view. The spots data from Trackmate was then processed using RStudio. Migration events were calculated for linked spots with centroid displacement greater than 2 pixels (∼0.002µm/s). Speed was calculated for each tracked event as difference in spot location per 5-minute interval. Migration angle was calculated using simple trigonometry so that parallel to the surface is 0°, towards the pia is -90°, and away from the pia is 90° as:

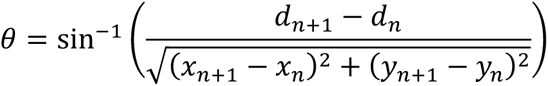

where *d* is DFS and (*x,y*) is location at timepoint *n*. The proportion of cells in the ML, as a tissue-level characteristic, was determined, reported, and statistically analyzed as biological replicates (N). Migration speed and direction were analyzed and reported as an aggregation of individual tracks from all biological and technical replicates as migration-specific parameters (n). The trends in migration speed and direction were graphed using ggplot in R for each metric based on the generalized additive method (Gam) in the geom_smooth package due to dataset size. Technical and biological replicates excluded for the previously indicated reasons are indicated in the associated R scripts used for this analysis.

### SNAP-pH adhesion in CGN and cerebellar glia coculture

For our first experiments of JAM-C-pH to JAM-B-SNAP adhesion, glia and CGNs were prepared from simultaneously prepared cell components. 8-well #1.5 glass bottom µ-slides (Ibidi) were prepared as previously described with Poly-ornithine and laminin. Glia were nucleofected with 30µg pCig2 JAM-B-SNAP cDNA. 50,000 of the nucleofected glia were then combined with a fresh preparation of 2,000,000 JAM-C-pH replaced CGNs. Live images were acquired the folin an environmentally controlled chamber on the Zeiss LSM 980 AiryScan microscope using a 63X 1.4NA objective, since it causes low photodamage and enhances resolution for contact formation after pixel reassignment using Zen Blue (Zeiss). Images were acquired as 6-8 µm z-stacks at 0.17 µm axial steps for multiple fields of view every 7 minutes overnight or every 3 minutes for 25 timepoints.

### Fluorescent Recovery After Photobleach (FRAP)

JAM-C-pH replaced CGNs were nucleofected as described for the morphology assays. 2,000,000 of the JAM-C-pH replaced CGNs were added to 1,000,000 non-nucleofected CGNs, 100,000 JAM-B-SNAP expressing glia, or 1,000,000 JAM-C-SNAP replaced CGNs into single wells of #1.5 coverslip bottom µ-slides (Ibidi). The conditions were live-imaged for FRAP experiments in an environmentally controlled on a Marianis Spinning Disk Confocal Microscope (Intelligent Imaging Innovation). Photobleach of pHluorin was performed using Vector high-speed x,y scanner 405 nm laser line in a 2×2µm square using a 25-raster block at 2ms. Because of microscope and variability in laser alignment, laser power was empirically set during biological replicates to achieve 80-90% reduction in fluorescence. Initial fluorescence was collected as Z-stacks to identify JAM-SNAP LSV-STL positive cells and GPI-2XBFP cell boundaries with Ex 560nm and Ex 405nm respectively, whereas photobleach and recovery were only imaged for pHluorin with Ex 488nm at representative Z-slices every 0.400 seconds. Fluorescent signal was measured within the 2×2µm photobleached region of interest in Slidebook and exported as an Excel document. Latrunculin A was used to reversibly depolymerize cortical actin. For drug treatment, 6X Latrunculin A prepared in GCM-FB was added directly to cells mounted in the environmental chamber on the microscope to a final concentration of 0.2 µg/mL. FRAP data was collected between 20-90 minutes of drug exposure, and then washed out for subsequent data collection. Parameters associated with local recovery of fluorescence were calculated as previously described [114]. Half time to recovery (t_1/2_) was calculated with nonlinear curve fitting in R to the equation:

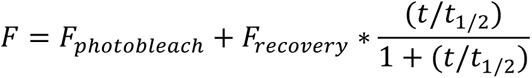

where F equals fluorescence intensity at time (t); F_photobleach_ is fluorescent intensity right after photobleach; and the parameters t_1/2_ and F_recovery_ are acquired from the non-linear fit for the half time to recovery and asymptote of fluorescent recovery respectively. The associated Mobile Fraction was calculated from the fit model as:

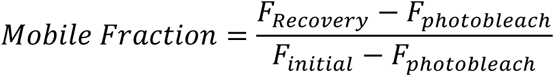

Instances when photobleached regions moved outside the regions of interest were manually excluded during final model fit as needed. Results were excluded from statistical comparisons if the dataset did not follow the described model as defined by: the mean residual from the model was >0.00001, mobile fraction calculated outside 0-1, or t_1/2_ greater than 40 seconds.

### Addition of secondarily dissociated JAM-pH replaced CGNs

Monolayers of CGNs and glia were prepared as described for the FRAP experiments. Monolayers were plated in #1.5 8-well dishes coated with 0.0025% Poly-ornithine (Sigma-Aldrich) that were dried, subdivided using 2-well silicone inserts (Ibidi), and coated with 1 µg/cm^2^ laminin. Without nucleofection, 500,000 CGNs or 25,000 glia were plated in subdivided wells to achieve 90-100% confluence. With nucleofection, 1,000,000 CGNs or 35,000-50,000 glia were plated in subdivided wells. Regardless of preparation considerations, monolayers were incubated overnight at 37°C and 5% CO_2_ mixed with humidified ambient air. The following day, for subdivided wells, silicone inserts were either left attached to glass or removed with sterile forceps as described for associated experiments.

JAM-pH replaced CGNs were prepared simultaneously to the monolayers. In all cases, 10,000,000 cells were nucleofected to replace endogenous JAM-C and immediately plated on multiwell uncoated glass slides overnight at 37°C and 5% CO_2_ mixed with humidified ambient air. JAM-pH replaced CGNs were aspirated from the uncoated slides with a cut pipette tip and secondarily dissociated based on the manufacturers guidelines in the Papain Dissociation System (Worthington Biochemical Corporation). Briefly, cell aggregates were dissociated at 37°C for 5 minutes with 150 µL of 20 units/mL of papain, 1mM L-cysteine, 0.5mM EDTA, and 100 units/mL of Deoxyribonuclease I (DNase I) in Earle’s Balanced Salt Solution (EBSS). Dissociated cells were then isolated by centrifugation through 10 mg/mL albumin and 10 mg/mL ovomucoid inhibitor in EBSS at 100xg in a fixed angle, tabletop centrifuge. The secondarily dissociated CGNs were then counted and added to respective monolayer conditions as specified in respective methods and figures.

### Imaging JAM-C-pH:JAM-B-SNAP adhesion formation with lattice lightsheet microscopy

We imaged the formation of contact formation between JAM-C-pH replaced CGNs and JAM-B-SNAP expressing glial monolayers using the Zeiss Lattice Lightsheet 7 for its speed and low photo damage. Glial cells were labeled with 1µM LSV-SNAPtag ligand for about 1 hour before washing and imaging. The glial monolayers in subdivided µ-slide were mounted and equilibrated in the environmentally controlled imaging chamber of the open top lattice lightsheet. The microscope stage was aligned to the invariant lightsheets using the monolayer sample. Coarse scanning of 560 excitation on JAM-B-SNAP glia was used to identify single glia exhibiting substantial fluorescence. Freshly dissociated CGNs were added to the open top of the dish and immediately imaged. Sample fluorescence was excited with 488nm and 560nm rectangular sheets generated with a 0.44NA objective and detected using a water immersion 50X 1.0NA objective capturing 50 ms exposure simultaneously on 2 Hamamatsu Fusion cameras. We collected an oblique stack at 0.3 µm intervals to cover volumes of about 200×200×20µm at 30 second temporal resolution for up to 1 hour. Raw images were deskewed and deconvolved using Zen Blue then exported as Tiffs for further processing and visualization in FIJI. Representative images of individual adhesion events were cropped from the deskewed images. Axial displacement caused by equilibration after cell addition was corrected manually. Photobleach correction was applied to the 560nm excitation channel by histogram equalization.

### SNAP-pH adhesion prevalence

JAM-C-pH replaced CGNs plated on untreated glass were secondarily dissociated as described. Monolayers were prepared with glia expressing JAM-B-SNAP and neurons expressing JAM-C-SNAP cultured in subdivided wells on a µ-slide coated with laminin that were treated with 1µM of the pH sensitive LSV-SNAPtag ligand for 1 hour. After labeling, 200,000 freshly dissociated CGNs were added to the monolayers. Cells were allowed to settle for 1 hour, washed extensively, and then imaged at 2 hours on one of two different Marianis Spinning Disk confocal microscopes (Intelligent Imaging Innovation) using a 20x, 0.8NA objective. We collected 12 µm Z-stacks at 1 µm intervals for 405 nm, 488 nm, and 560 nm excitation at excitation powers chosen empirically based on the ability to detect LSV labeled glia and CGNs in the 560 channel. Max-projection fluorescent images were generated in Slidebook (Intelligent Imaging Innovation) and used for all subsequent analyses.

Ilastik semi-automated image segmentation was used to acquire three separate masks for each biological replicate [72, 73]. 1) Monolayer cell bodies as viable area for cell adhesion were segmented using pixel classification and object classification trained on the constitutive GPI-2XBFP in the 405nm excitation channel for all conditions. 2) JAM-SNAP positive adhesions were segmented with pixel classification trained to distinguish locally bright regions in the 560nm excitation associated with LSV labeling. 3) JAM-C-pH adhesions were segmented with pixel classification trained to distinguish locally bright regions of 488nm excitation. Data was analyzed by combining all three segmentations in FIJI. Analyze particles was applied to monolayer segmentation and JAM-SNAP segmentation to measure co-occurrence of pHluorin adhesion. Analysis of output data files was performed in RStudio. Prevalence of SNAP-pH adhesions was measured as the proportional area of the segmented JAM-SNAP adhesions within the monolayer that coincides with pHluorin segmentation. Data from multiple fields of view were aggregated into single biological replicates for reporting and statistical comparison. Biological replicates were excluded from statistical comparison if proportional overlap of SNAP-pH was 1.5 times outside of the condition’s inter quartile range.

### Imaging recruitment of Drebrin and Pard3 to JAM contacts after secondary dissociation

Recruitment of cytoplasmic binding partners to extracellular adhesion sites was performed by secondary dissociation. JAM-C-pH replaced CGNs coexpressing drebrin-SNAP and Halo-Pard3 cultured on untreated glass were initially labeled with 1 µM JF552cp-SNAPtag ligand and 1 µM JFX 650x Halotag ligand for 1 hour. Monolayers were prepared from CGNs or glia that were nucleofected to express either JAM-B-pH or JAM-C-pH. Monolayers were equilibrated for imaging in an environmentally controlled chamber on a Marianis Spinning Disk Confocal Microscope (Intelligent Imaging Innovation) using a 63X 1.46NA oil objective and SoRa microlens disk. Freshly dissociated CGNs were warmed to 37 °C and directly added to the monolayers on the microscope. Fluorescence was collected using a quad-pass filter for 488 nm (50ms), 560 nm (35 ms), and 640 nm (50ms) for 5-7 µm Z-stacks sampled at 0.5µm axial steps for as many fields of view to achieve 3 (1 FOV), 15 (3 FOVs), or 30 (5-6 FOVs) second intervals. Max projections were generated in Slidebook (Intelligent Imaging Innovation). Individual adhesion events were identified and manually cropped from the max projection images. To preprocess adhesion events, fluorescent intensities were normalized and change in position over time was reduced by rigid type registration of the images based on monolayer JAM-pH signal over time via ANTs software [115, 116]. The registered images were then used for downstream analysis of signal correlation.

### Analysis of bulk recruitment with Pearson’s Correlation

Pearson’s correlation for pairwise comparisons of JAM-pH, drebrin-SNAP, and Halo-Pard3 over time were calculated using Just Another Colocalization Program (JaCoP) plugin in FIJI [117]. We developed a FIJI macro to facilitate a series of user inputs for each event. We first cropped the registered image to the area of adhesion to reduce null area included in correlation calculation. Next, condition-specific thresholds of signal minima and maxima for each channel were manually specified. From there, the time point at which adhesion is first visible is manually identified. Finally, the Pearson’s correlation is calculated iteratively for each timepoint as pairwise comparisons of the three fluorescent signals. Technical replicates were excluded from if any of the three signals exhibited insufficient normalized fluorescence or the JAM-pH adhesion signal was unable to be properly segmented by intensity thresholding. Output data from FIJI was assembled into a data table and analyzed in RStudio. Technical replicates across ≥3 biological replicates per condition were aggregated, plotted, and statistically compared. Trends as functions of time after first sign of cell adhesion or size of adhesion were plotted without assumption of model for the data uniformly subsampled at 30 seconds using the LOESS method in geom_smooth.

### Analysis of local recruitment with Pearson’s Correlation

Local correlation was evaluated as the Pearson’s correlation in a kernel of defined pixel dimension for ANTs registered data using a python environment in a Jupyter notebook. The kernel was scanned over the image for each time point, plotting local correlation superimposed at the kernel center. Kernel size was selected based on pilot analysis of local correlation between glia JAM-B-pH and drebrin-SNAP in JAM-C WT replaced CGNs. Local correlation maps were generated for square kernels ranging from 3×3 up to 15×15 pixels. We applied a manually determined segmentation mask of JAM-B-pH adhesion to the pixel map to identify the impacts of kernel size on local correlation. This approach revealed that sensitivity of local correlation decreased with increased kernel size. We selected the 9×9 kernel size as a proof of principle for this novel approach that balanced tradeoffs of noise and local sensitivity.

Pixel maps of the local correlation for JAM-pH to Pard3 and JAM-pH to drebrin were generated for the preprocessed, registered adhesion events. These pixel maps were analyzed based on user input using a macro in FIJI. First, the images were cropped to the adhesion site to help isolate single adhesion events. The time of first adhesion and thresholds were manually set for JAM-pH, drebrin-SNAP, and Halo-Pard3. Associated metrics of local correlation at the segmented JAM-pH contacts were determined in Rstudio using the script provided in the supplemental materials. Analysis of the sum correlation of positive or negative correlation was applied within the segmented mask on Pearson > 0 and Pearson < 0 respectively. Images were excluded from analysis if we identified inadequate normalized fluorescence or could not segment the JAM-pH adhesion using simple thresholds. Technical replicates across ≥3 biological replicates per condition were aggregated, plotted, and statistically compared. Trends as functions of time after first sign of cell adhesion or size of segmented adhesion were plotted without assumption of model for data uniformly subsampled at 30 second intervals using the LOESS method in geom_smooth.

### Analysis of the spatial distribution of local Pearson’s Correlation

We anecdotally observed peripheral positioning of drebrin fluorescence at JAM adhesions to glia, so we developed a novel approach to quantify this behavior by calculating the spatial distribution of the correlation maps within the segmented adhesions. We determined the contribution of the peripheral component of segmented adhesions by calculating the ratio of area and correlation measured at the periphery divided by the area and correlation from the total adhesion. A ratio of 1 signifies that all measured signal was present in the adhesion periphery, while a ratio of 0 signifies none of the signal measured in the total adhesion is contributed by the peripheral component. We generated distance maps of the segmented adhesions in Fiji. Peripheral adhesion was defined as regions within 2 pixels of the edge of adhesion (<0.5 µm). Incorporating these distance maps into our analysis we calculated the sum correlation and adhesion area for the regions that satisfy the 2-pixel criteria. We combined the local correlation datasets for the entire adhesion and the peripheral subset to determine the proportional contribution of JAM periphery to adhesion area, sum positive correlation, and sum negative correlationfor JAM:Pard3 or JAM:drebrin using the respective ratios:

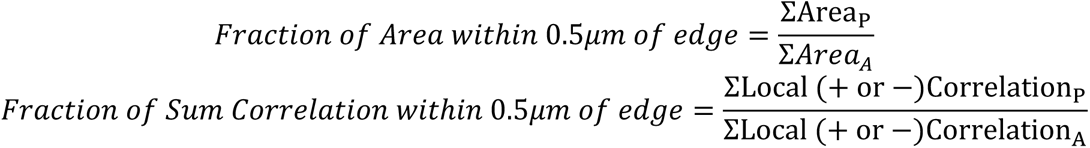

The subscripts P and A apply to the values calculated for just the edge or the entire adhesion respectively. The difference of the ratio of adhesion size and the ratio of correlation indicate that the distribution of correlation values in the periphery is different from that of the whole adhesion. Difference values greater than the adhesion size ratio indicate peripheral distribution, and values less than the adhesion size ratio indicate peripheral distribution. To generalize these measurements of spatial distribution to compare mutant receptors with potentially different adhesion characteristics, we calculated the difference between the peripheral ratio of local correlation and peripheral ratio of adhesion area as shown:

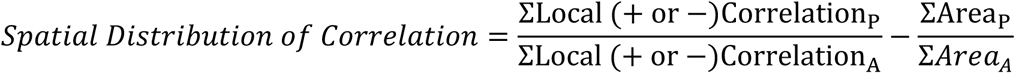

Resulting values greater than 0 indicate peripheral spatial distribution of sum correlation, whereas values less than 0 indicate an internal spatial distribution of sum correlation. The resulting trends in spatial distribution of positive or negative correlation as functions of time after first sign of cell adhesion or size of segmented adhesion were plotted without assumption of model for data uniformly subsampled at 30 second intervals using the LOESS method in geom_smooth.

### Western Blotting

All samples were prepared as formulations with 1X LDS sample buffer and 10 mM DTT that were incubated at 70 °C for 10-15 minutes and centrifuged at 4 °C 16,000xg for 5 minutes. Samples were added to Bolt™ 4-12% Bis-Tris Plus (Invitrogen) wedge-well polyacrylamide gels and were electrophoresed in a Mini Gel Tank (Invitrogen) using MES buffer at 200V for 25 minutes. Protein was transferred to an Immobilon PVDF membranes (Millipore Sigma) via wet transfer at 30V for 1 hour at 4 °C in 10% Methanol Tris/Glycine buffer using a Criterion™ Blotter (BioRad). Blots were all blocked for 1 hour at room temperature using Intercept (TBS) Blocking Buffer diluted 1:1 in TBS. Primary antibodies were applied to PVDF blots overnight at 4 °C as specified in the Key resources table then washed with TBS containing 0.1% Tween 20 (TBST). IRDye® (LI-COR) fluorescent secondary antibodies were applied to rinsed PVDF blots at room temperature and washed with TBST. All western blot images were acquired using an Odyssey CLx scanner (LI-COR) and band total signal analyzed with the associated Image Studio software.

### Statistics

Rstudio was used in this work to generate all plots and perform statistical analyses. The raw data used in statistical comparisons are plotted directly on the graphs as described in the figure legends. Due to limited sample sizes and visible trends that did not follow normal distribution in pairwise comparisons, we used Wilcoxon Rank-sum as a non-parametric method for the comparisons designated in figure legends. Significance is reported following the standard notation p<0.05 *, p<0.01**, p<0.001 ***, and p<0.0001 ****. Exact p-values for experiments reported in this work are provided in the supplemental materials.

### Cycloheximide Pulse-Chase

The relative stability of JAM-pH constructs was measured in lipofected Neuro-2a cells by tracking protein after blocking translation with 10 µg/mL cycloheximide similar to previously described methods [118]. First, 125,000 Neuro-2a cells were plated in 0.5mL on a cell-culture treated 24-well plate and incubated overnight at 37 °C and 5% CO_2_ mixed into ambient O_2_ to achieve 50-60% confluence. The following day, lipofection conditions were prepared as 225 µL of Opti-MEM® (Gibco) containing 1.5 µg pCig2 Cre T2A H2B-mCherry and 1.5 µg pCA LNL JAM construct was combined with 225 µL of Opti-MEM® with 9 µL of Lipofectamine™ 2000 (Invitrogen). 75 µL of this lipofection formulation was added to 6 wells of Neuro-2a cells. The cells were incubated overnight at 37 °C and 5% CO_2_ mixed into ambient O_2_ to achieve stable expression. The following day, 50 µL of 200 µg/mL in Opti-MEM® was added to wells as designated time points ranging from 0h up to 16h to a final concentration of 10 µg/mL cycloheximide. At the final time point, all cells were rinsed with DPBS and lysed in 100 µL of Pierce™ IP lysis buffer containing 1X Halt™ Protease and Phosphatase inhibitor (Thermo Scientific). The amount of JAM-pH in the sample was measured by Western Blot detection of GFP in the lysates. Protein half-life was calculated in GraphPad Prism based on the Least-squares fit of the data to one-phase decay. Biological replicates of half-life are plotted and statistically compared in RStudio.

### Surface Biotinylation

The relative amount of JAM-C-pH at the cell surface was measured in lipofected Neuro-2a cells by performing surface biotinylation followed by streptavidin pull-down based on manufacturer’s guidelines. 1,000,000 Neuro-2a cells were plated in 4mL of media on a 60mm cell-culture treated dish and incubated overnight at 37 °C and 5% CO_2_ mixed into ambient O_2_. The next day cells were lipofected with a 600 µL formulation containing 12 µL Lipofectamine™ 2000 (Invitrogen), 2µg pCig2 Cre T2A H2B-mCherry, and 2 µg pCA LNL JAM construct. The cells were again incubated overnight at 37 °C and 5% CO_2_ mixed into ambient O_2_ to achieve stable expression. The following day cells were cooled to 4 °C and treated with 0.5 mg/mL EZ-Link Sulfo-biotin-SS-NHS (Thermo Scientific) dissolved in 1XDPBS for 30 minutes. Reactions were quenched with 2 washes of 50mM Tris pH 7.5 in 1XDPBS and lysed in 300 µL Pierce™ IP lysis buffer containing 1X Halt™ Protease and Phosphatase inhibitor (Thermo Scientific). 50 µL of protein sample was set aside for subsequent analysis, and 250 µL of lysate was added to a 75 µL bead volume of Streptavidin Agarose Resin (Thermo Scientific). The lysate and beads were incubated at 4 °C for 1 hour by end-over-end mixing. Beads were rinsed twice with 250 µL lysis buffer and eluted in 50µL of reducing sample buffer at 70 °C. Protein composition was identified using our standard Western Blot procedure. JAM-pH surface biotinylation was measured as the ratio of biotinylated GFP detected to the amount of GFP detected in the input. The biological replicates are plotted and compared using ggplot in RStudio.

## Data Availability

The numerical data used for the analyses performed in this work are included in the manuscript and supporting files. Summaries of p-values with associated plots over time for each figure are provided in supporting files. The AlphaFold Multimer predictions the structure for 4 subunits of JAM-C extracellular domain or 2 subunits of JAM-B extracellular domain with 2 subunits of the JAM-C extracellular domain is provided in supporting files. The Fiji Macros, R scripts, and Jupyter Notebook used for analysis throughout this manuscript are available in the supporting files. Primary imaging data, processed images, and intermediary image analysis will be made available upon reasonable request.

---

**Figure 1 – figure supplement 1.**
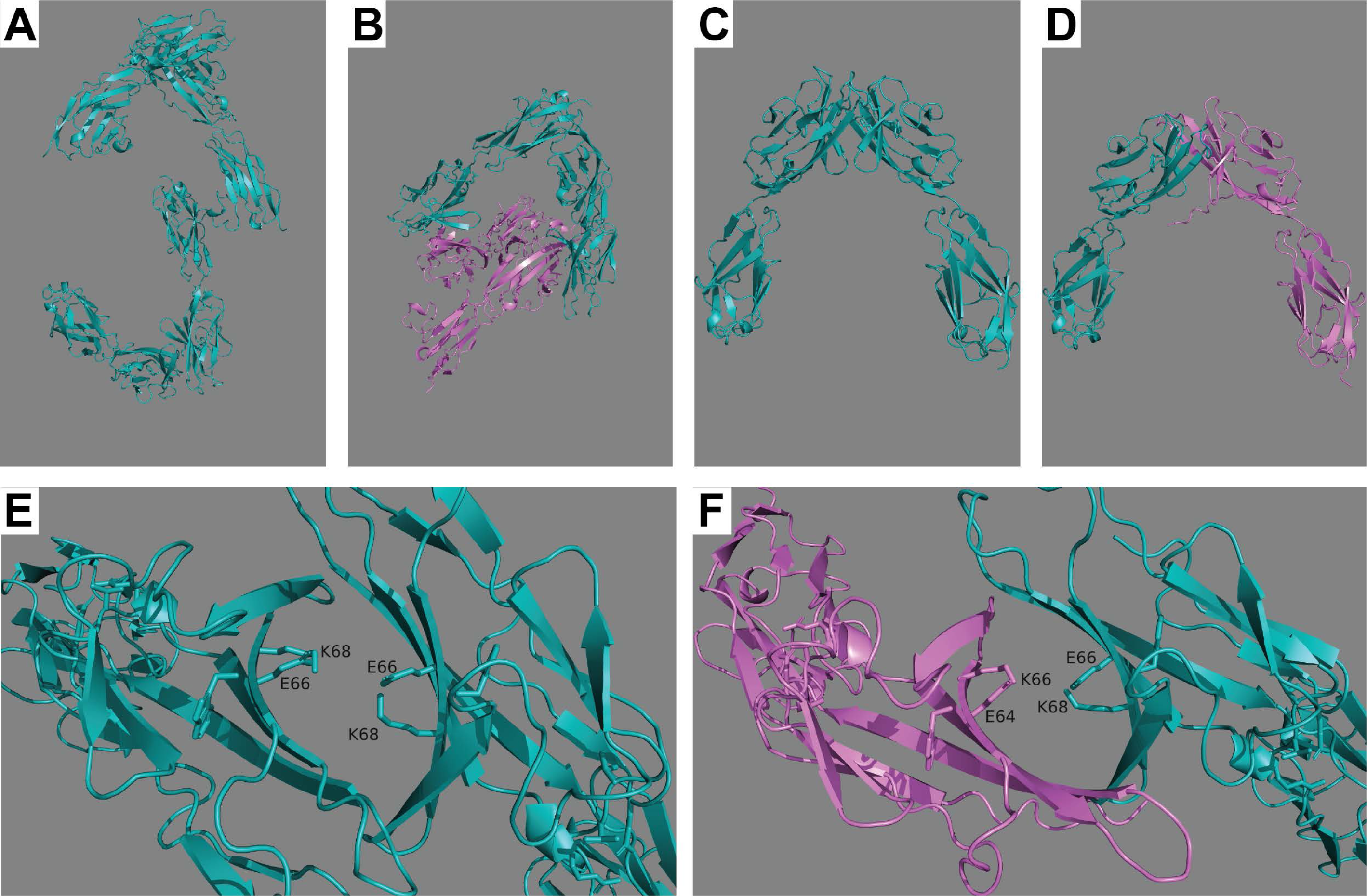
AlphaFold Multimer prediction of JAM dimers. Representative view of the predicted structure of 4 JAM subunits generated for either (**A**) 4 JAM-C extracellular domains or (**B**) 2 JAM-C with 2 JAM-B extracellular domains. The prediction yielded two independent cis interacting dimers dimers. Isolation of the predicted dimers from the 4 subunit predictions for (**C**) JAM-C to JAM-C and (**D**) JAM-C to JAM-B. The structural prediction shows that these dimers are predicted to form at the BLAST aligned charged residues for (**E**) JAM-C to JAM-C and (**F**) JAM-C to JAM-B cis adhesion sites.

**Figure 1 – figure supplement 2.**
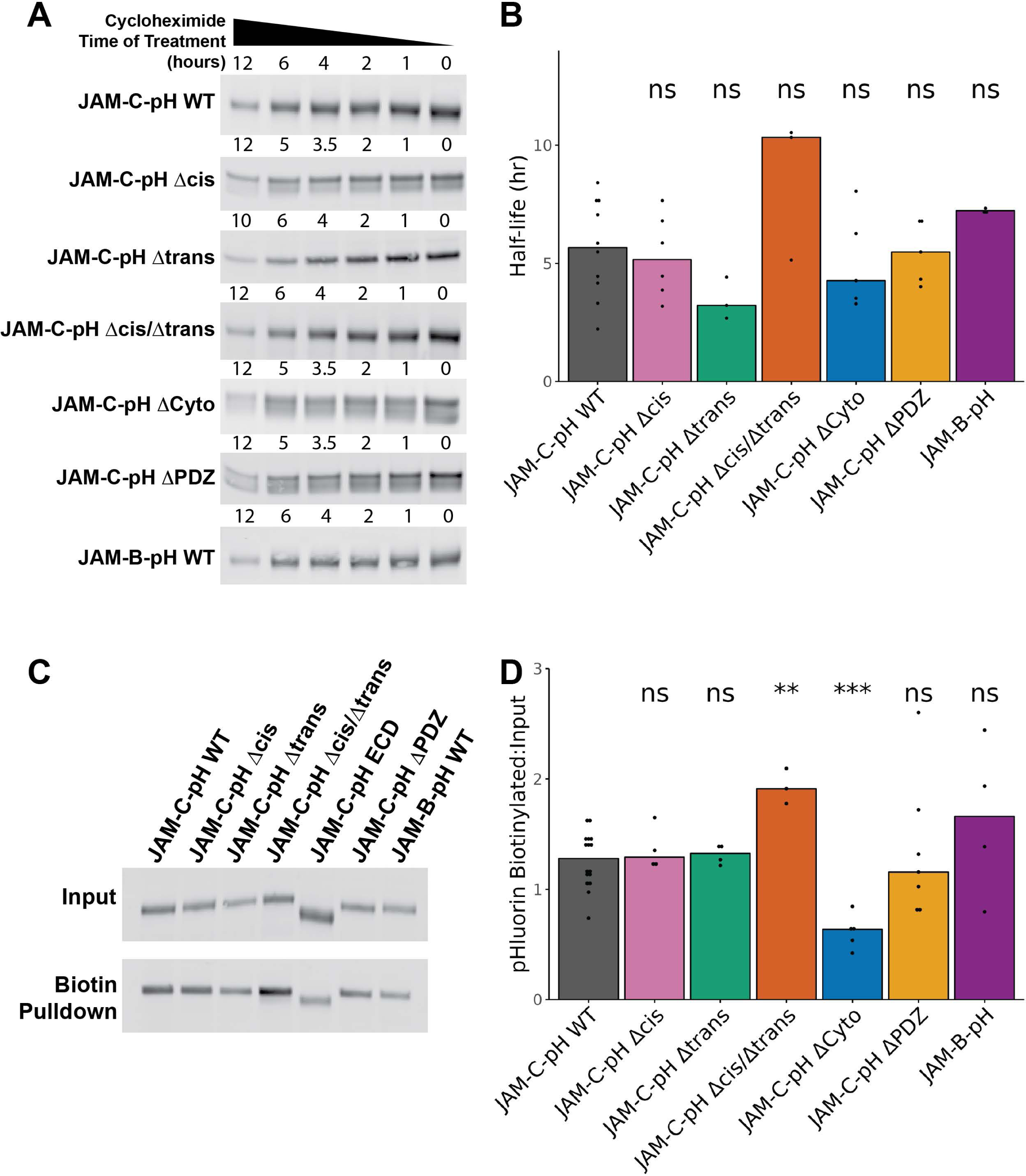
Validation of JAM-pH fluorescent protein construct stability and trafficking in Neuro-2a cell line. (**A,B**) Cycloheximide pulse was used to block protein translation and then protein stability was tracked over time via Western Blot. (**A**) Representative western blot images of cycloheximide time course for JAM-C-pH expressing Neuro-2a cells immunoblotted with Chicken αGFP (Aves). (**B**) Independent biological replicates of half-life calculated in GraphPad Prism by one-phase decay are plotted and statistically compared to JAM-C-pH WT via Wilcoxon Rank Sum test. (**C,D**) Trafficking of JAM-pH to the cell surface was measured with surface biotinylation. (**C**) Representative western blot images of Neuro-2a lysate and biotin pull down by streptavidin functionalized agarose beads (Thermo Scientific) immunoblotted with Chicken αGFP (Aves). (**D**) Surface biotinylation was quantified as the ratio of the JAM-pH detected in the pull down divided by the JAM-pH detected in the input. The ratios for biological replicates (N≥3) are plotted and statistically compared to JAM-C-pH WT using a Wilcoxon Rank Sum test. Statistical significance is represented by standard convention: ns p>0.05, * p<0.05, ** p<0.01, *** p<0.001, and **** p<0.0001.

**Figure 1 – figure supplement 3.**
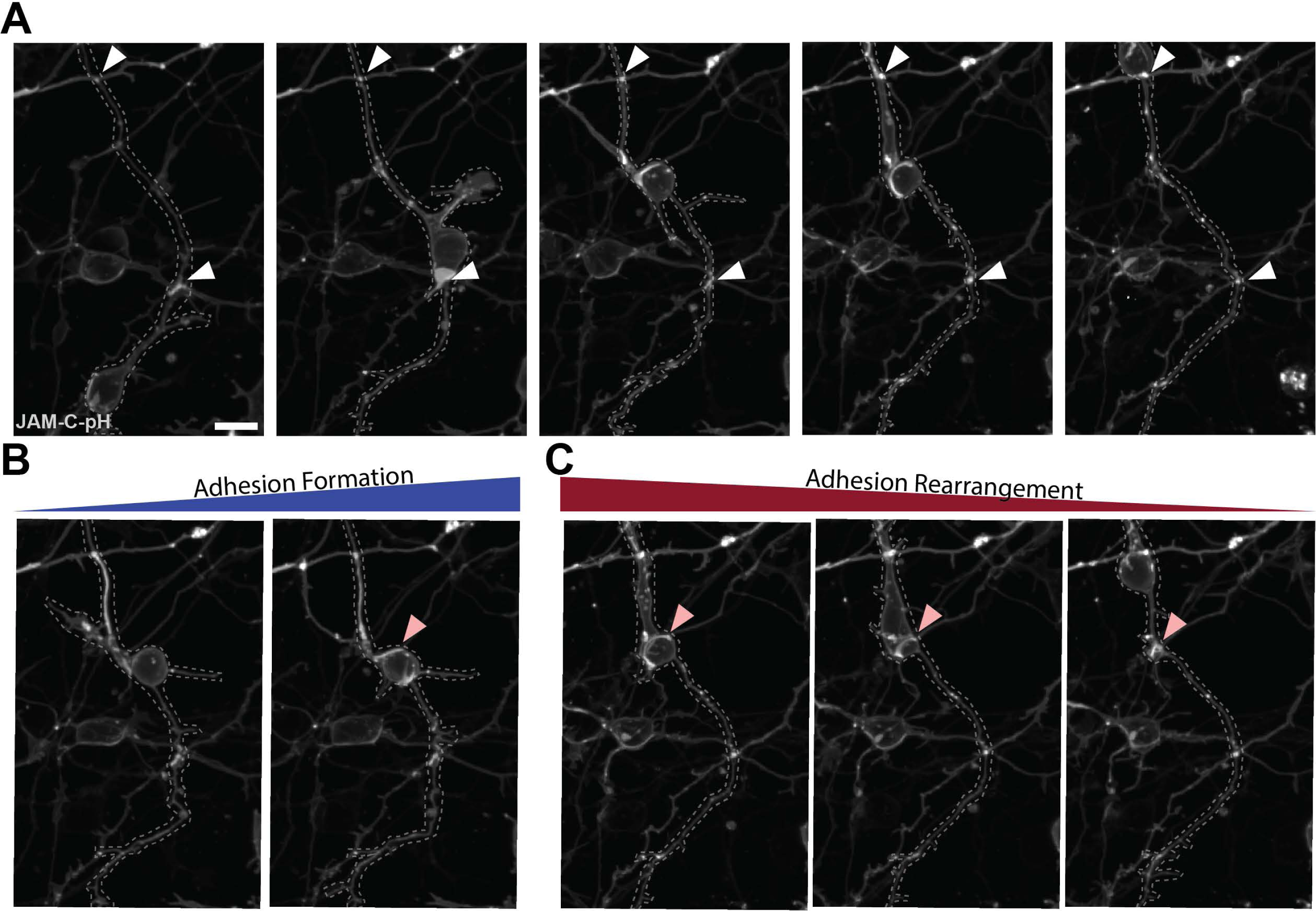
JAM-C-pH adhesions exhibit a wide range of dynamics in live JAM-C replaced CGNs. Representative Ex 488 nm fluorescence images of the migration of JAM-C-pH WT replaced CGNs. The images shown are max projections of 3D data acquired at 63x 1.46NA that have been gamma adjusted at 0.75 to enhance low signal areas. The migrating neuron is outlined in a dashed grey line. (**A**) Some adhesions are observed to last longer than 12 hours. A couple of instances of this behavior are noted with the presence of white arrowheads that remain relatively stationary for the entire time course that do not exhibit strict biases in subcellular location. (**B,C**) Within the time course, new adhesions are observed to form and undergo rearrangement. (**B**) A soma localized adhesion was observed to form within a 20-minute temporal interval marked with a pink arrowhead. (**C**) As an example of adhesions not maintaining a constant size, the cell migrating past the adhesion site marked with the pink arrowhead appears to stretch the fluorescent signal until the soma passes the adhesion location allowing the signal to contract laterally.

**Figure 1 – figure supplement 4.**
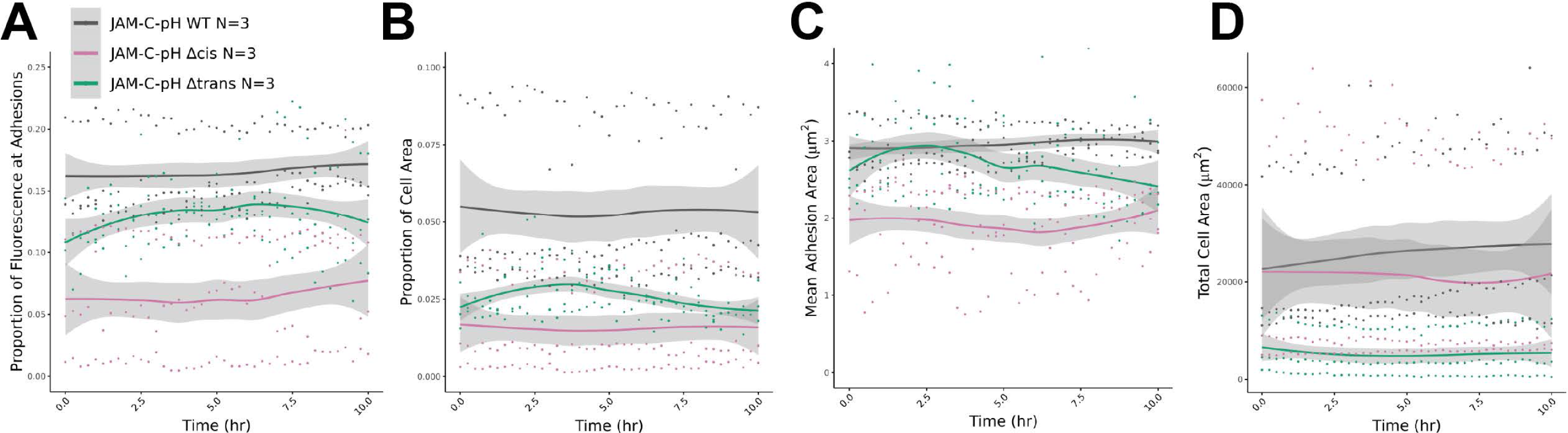
Total JAM-C-pH adhesion in JAM-C replaced CGNs is at a relative steady state 24 hours after nucleofection. (**A-D**) JAM-C-pH adhesions were live imaged overnight every 15 minutes to track adhesion over time. The data displayed is a Loess smoothed trend for the data points from N=3 biological replicates. In general, the (**A**) proportion of fluorescence at adhesions, (**B**) Proportion of cell area with adhesion, (**C**) median adhesion area measured in µm^2^, and (**D**) total cell area measured in µm^2^ are relatively constant over the 10-hour time course. Gray shading on the plots represents the 95% confidence intervals of the estimated means.

**Figure 1 – figure supplement 5.**
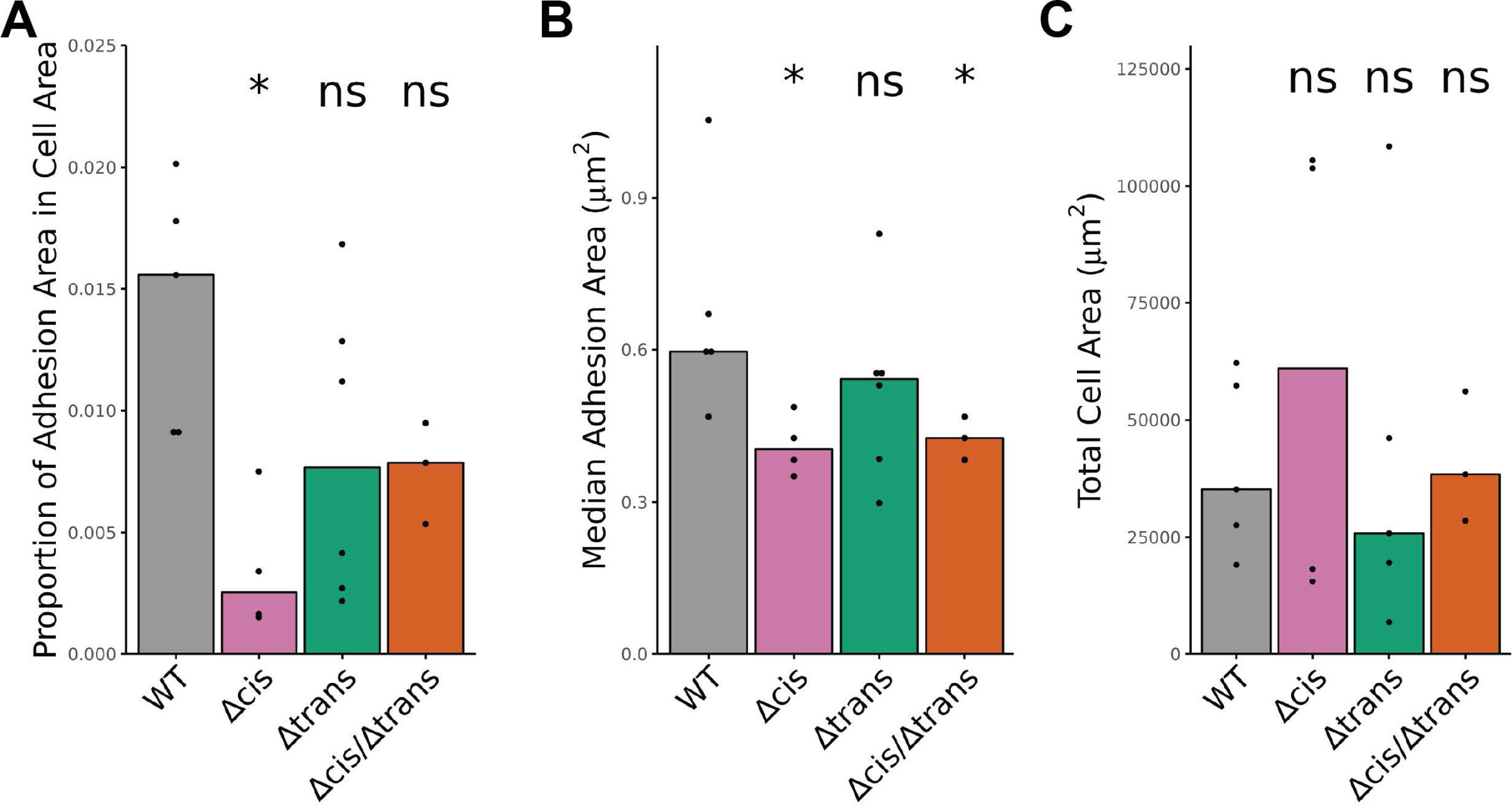
The quantification of JAM-C-pH adhesion is impacted by proportion of cell area and adhesion size but not total cell area. Comparisons of (**A**) proportion of cell area with segmented adhesions, (**B**) median adhesion area measured in µm^2^, (**C**) and the total analyzed cell area measured in µm^2^ at 24 hours of culture for independent biological replicates each compared to JAM-C-pH WT replaced CGNs using Wilcoxon Rank Sum tests. Statistical significance is represented by standard convention: ns p>0.05, * p<0.05, ** p<0.01, *** p<0.001, and **** p<0.0001.

**Figure 2 – figure supplement 1.**
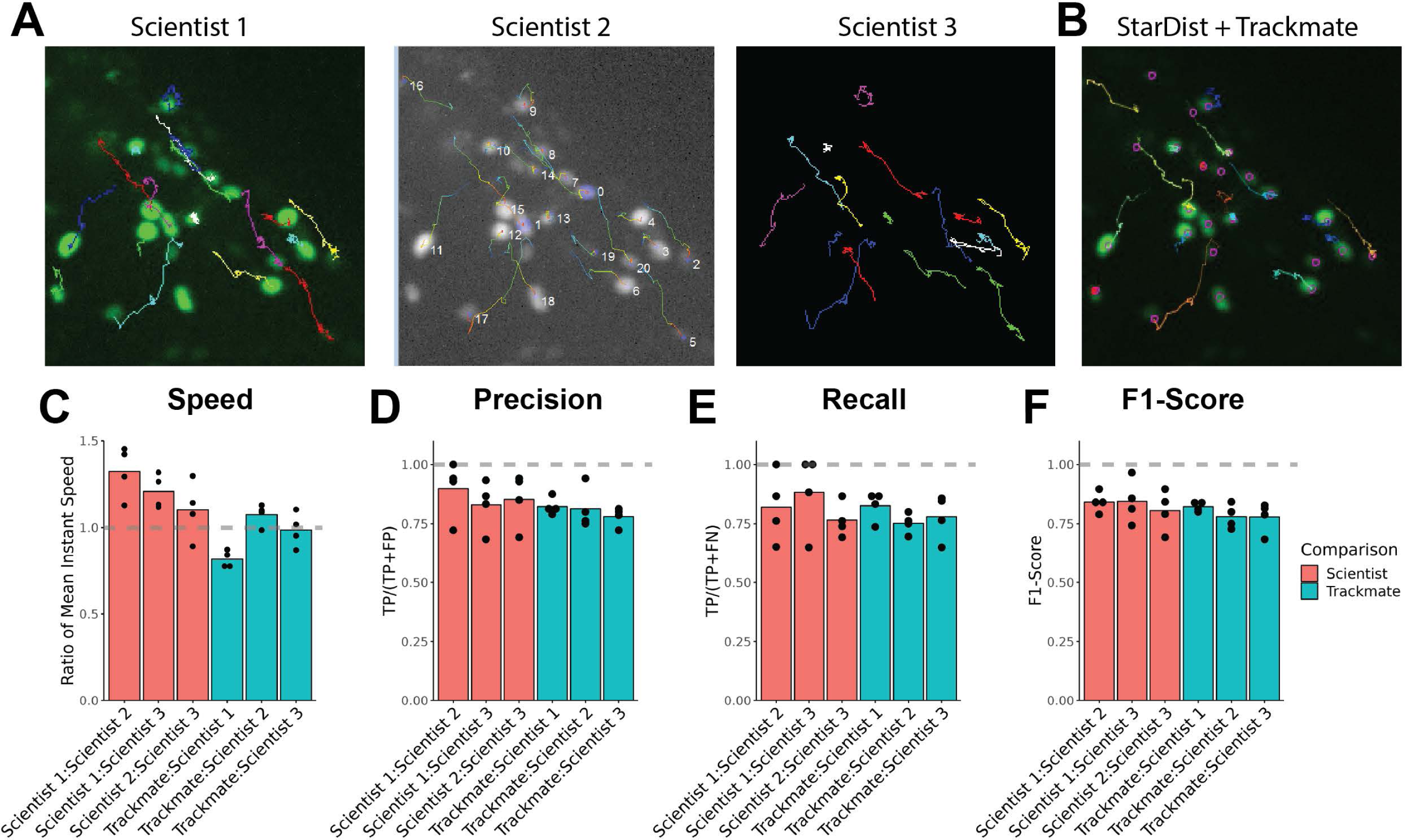
Trackmate and Stardist can track nuclear labled CGNs in cerebellar slice cultures. (**A**) Representative examples of manually tracked H2B-mCherry-labeled CGNs from 3 independent scientists (**B**) Representative example of CGN tracking using StarDist and Trackmate. (**C-F**) Quantification of the reproducibility of tracking performed on 4 separate data sets. Dashed lines are placed on bar charts at the ideal value for reproducibility for the associated metrics. Data points represent quantified values of independent biological replicates (**C**) Reproducibility of individual migration events was quantified as the ratio of migration speeds for the comparisons listed on the x-axis. Overall track quality was quantified by categorically annotating successful tracks and plotting the output (**D**) precision, (**E**) recall, and (**F**) F1-Score.

**Figure 2 – figure supplement 2.**
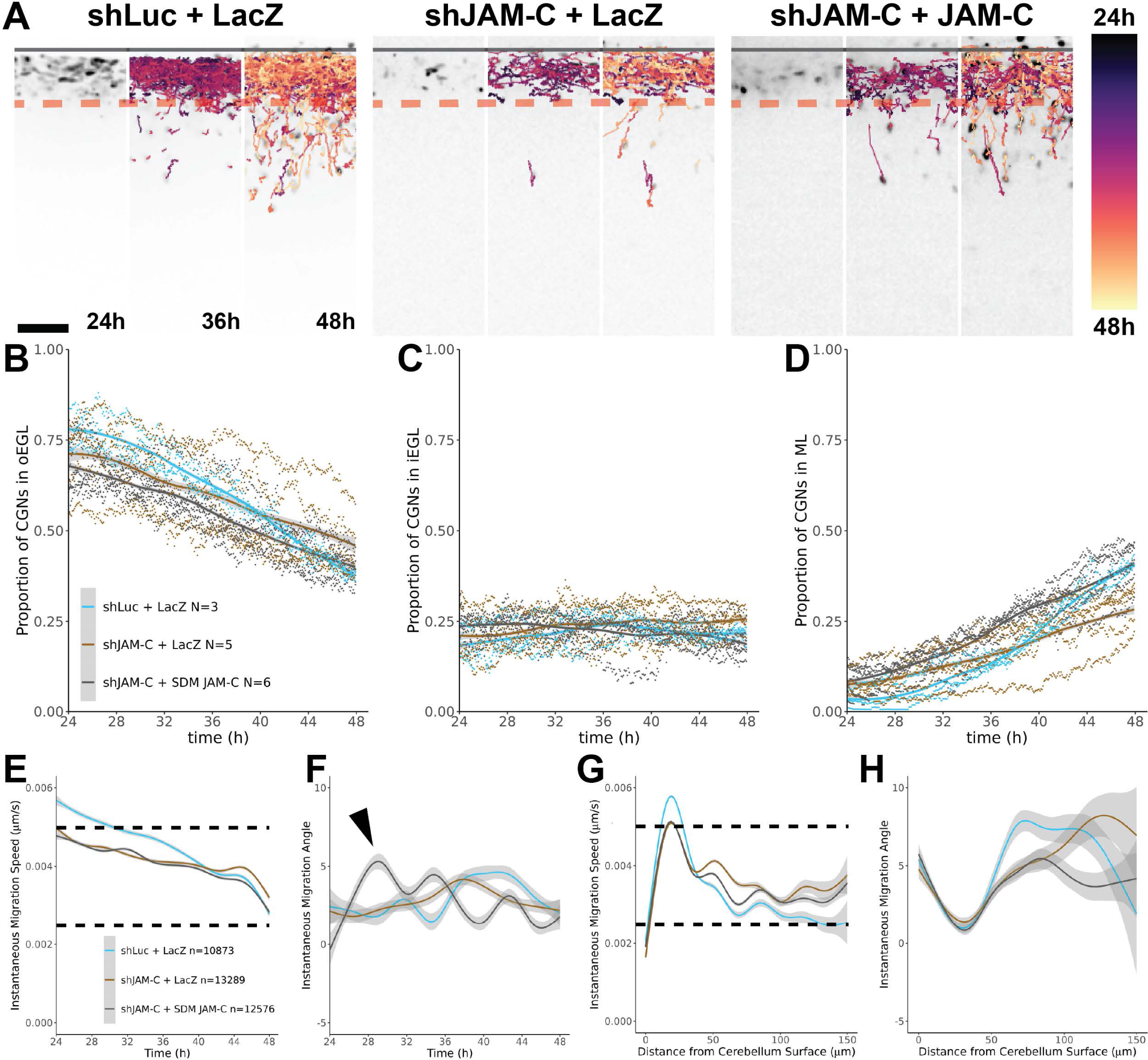
Tracking CGN migration for JAM-C rescue with StarDist and Trackmate reveals subtle changes in migration behavior at 24-48 hours. (**A**) Representative tracking data for validating JAM-C rescue at 24, 36, and 48 hours. Track position in time is color encoded based on the provided lookup table. Quantification and plotting of the proportion of CGNs in the (**B**) oEGL, (**C**) iEGL, and (**D**) ML over time. Aggregated proportion for each biological replicates is plotted directly on the graph at each time point, and the smoothed trend over time was plotted using gam. (**E-H**) Quantification of the migration tracks for each condition, aggregating data for tracked migration events across biological and technical replicates (n). Data is plotted as a gam smoothed trend as a descriptive representation of migration behavior. Gam plots of (**E**) migration speed and (**F**) migration angle as functions of time. Gam plots of (**G**) migration speed and (**H**) migration angle data as functions of distance from tissue edge. Gray shading on the plots represents the 95% confidence intervals of the estimated mean values.

**Figure 2 – figure supplement 3.**
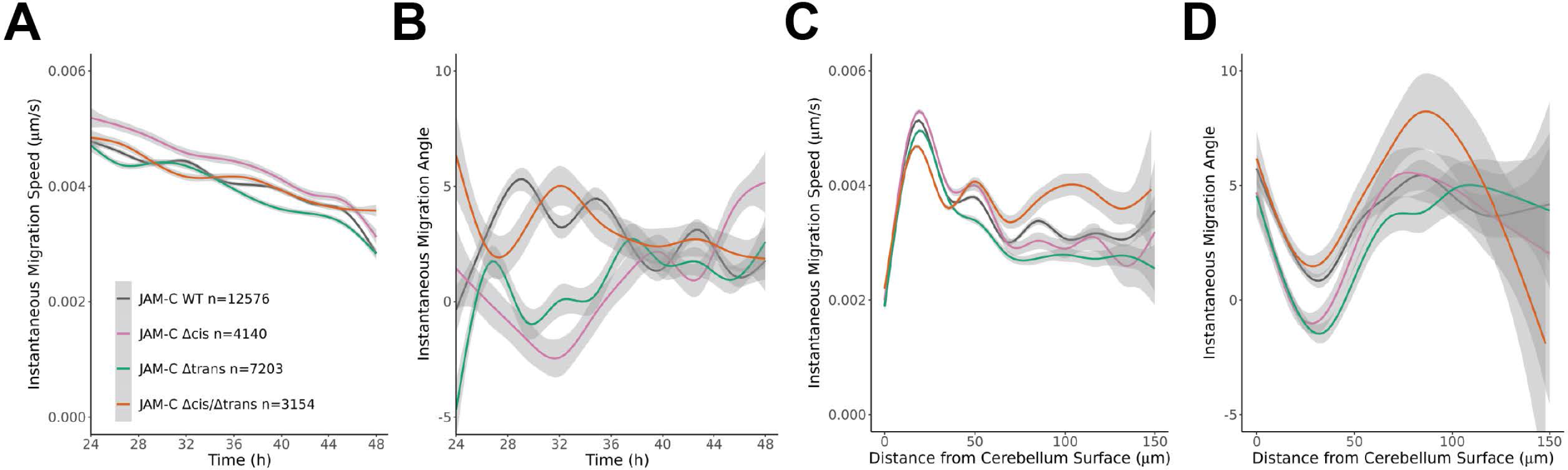
Migration track data for JAM replacement in slice cultures. (**A-D**) Quantification of migration tracks for each JAM replacement is shown, aggregating data for tracked migration events across biological and technical replicates (n). Data is plotted using gam smoothing as descriptive representations of migration behavior overall. Gam plots of (**A**) migration speed and (**B**) migration angle as functions of time. Gam plots of (**C**) migration speed and (**D**) migration angle data as functions of distance from tissue edge. Gray shading on the plots represents the 95% confidence intervals of the estimated mean values.

**Figure 4 – figure supplement 1.**
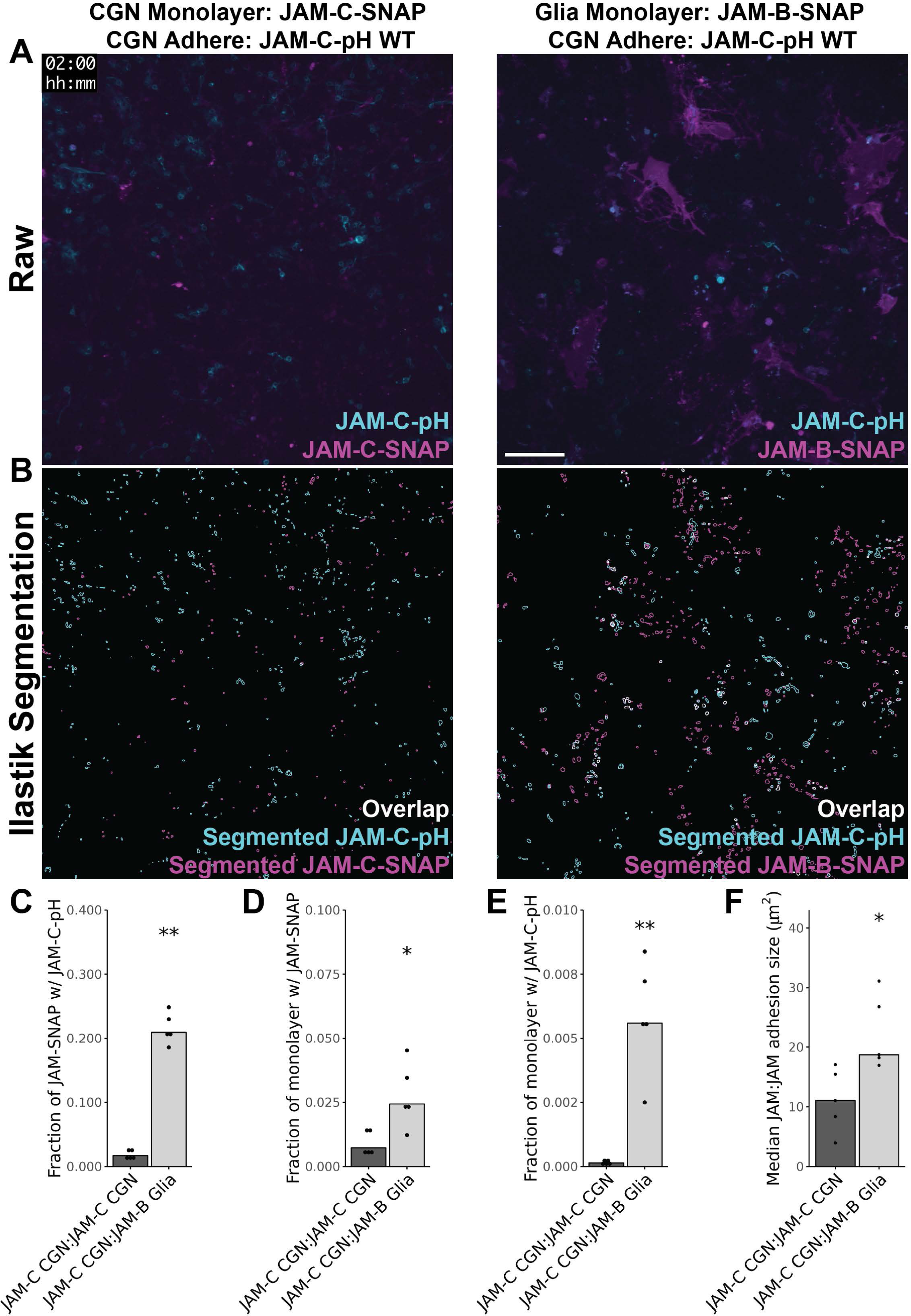
Stable de novo JAM contacts are more likely to form between CGN JAM-C and Glia JAM-B than between CGNs with JAM-C. (**A**) Representative 0.25 gamma adjusted, max projected, fluorescent images for JAM-C-pH WT replaced CGNs adhering to either JAM-C-SNAP expressing CGNs or JAM-B-SNAP expressing glia at 2 hours for entire fields of view acquired by a 20x 0.8 NA objective. Scale bar = 100 µm (**B**) Ilastik segmentation for JAM-C-pH and JAM-SNAP adhesions corresponding to the fluorescent imagers in (**A**). Segmentation is shown as the traces of the boundaries for segmented objects. (**C-F**) Quantification and statistical comparison of SNAP-pH adhesion parameters for JAM-SNAP expressing monolayers. (**C**) Fraction of JAM-SNAP with JAM-C-pH is defined as the overlapped area of Ilastik segmented JAM-C-pH and JAM-SNAP divided by total JAM-SNAP segmented area. (**D**) Fraction of monolayer with JAM-SNAP is defined as the proportion of SNAP segmented area within the cell body mask divided by the total area of the cell mask. (**E**) Fraction of monolayer with JAM-C-pH is defined as the proportion of pHluorin segmented area within the cell body mask divided by the total area of the cell mask. (**F**) Median JAM:JAM adhesion size is determined as the median area of JAM-SNAP segmented objects that overlap with JAM-C-pH segmented signal. Data points are reported as individual biological replicates statistically compared using Wilcoxon Rank Sum Tests. Statistical significance is represented by standard convention: ns p>0.05, * p<0.05, ** p<0.01, *** p<0.001, and **** p<0.0001.

**Figure 4 – figure supplement 2.**
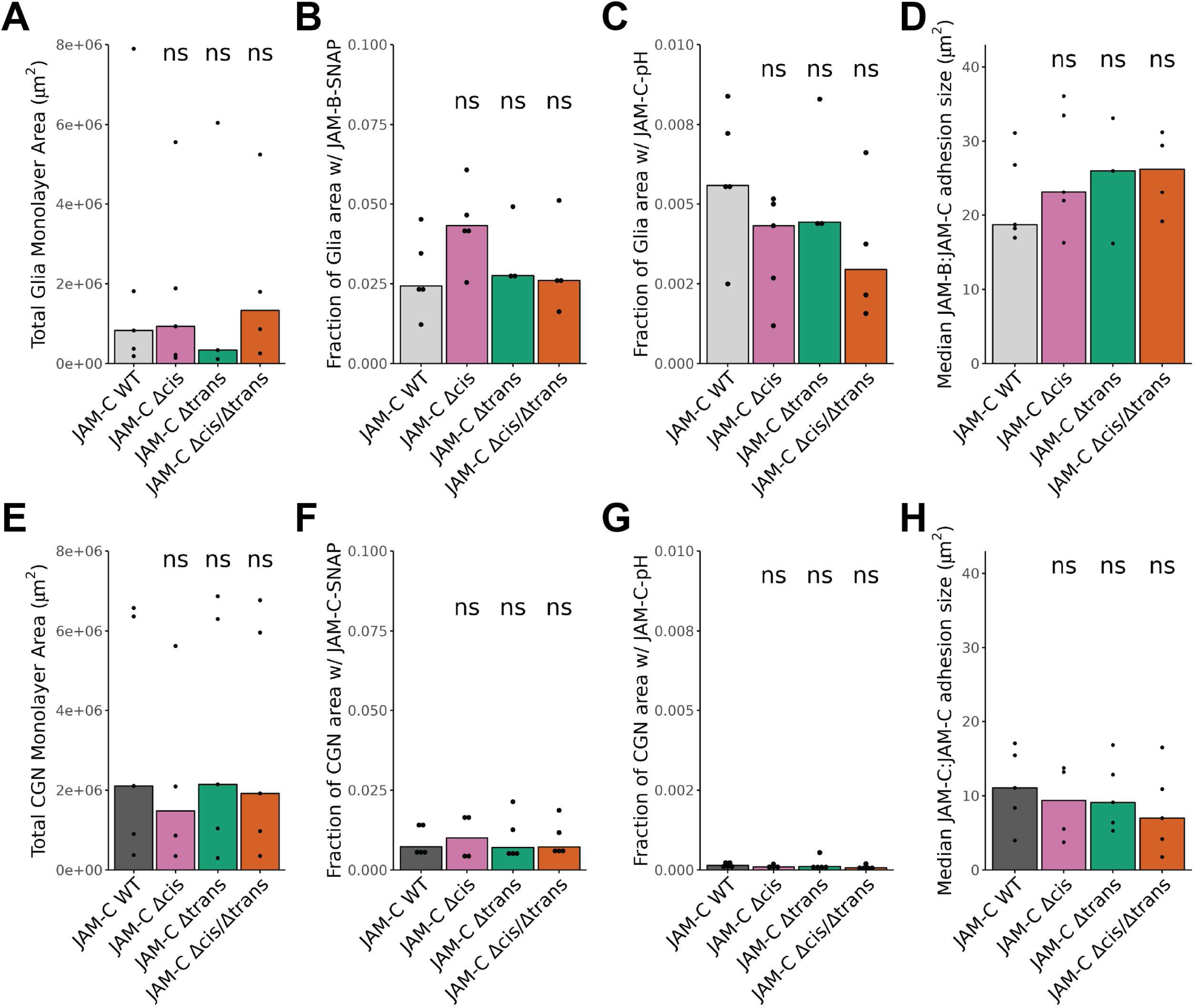
JAM-C binding in *cis* or *trans* did not significantly alter other parameters of SNAP-pH adhesion. (**A-H**) Quantification and statistical comparison of SNAP-pH adhesion parameters for (**A-D**) JAM-B-SNAP expressing glia and (**E-H**) JAM-C-SNAP expressing CGNs. (**A,E**) Total cell monolayer area is defined as the summed area of the cell body mask for biological replicates. (**B,F**) Fraction of monolayer with JAM-SNAP is defined as the proportion of SNAP segmented area within the cell body mask divided by the total area of the cell mask. (**C,G**) Fraction of monolayer with JAM-C-pH is defined as the proportion of pHluorin segmented area within the cell body mask divided by the total area of the cell mask. (**D,H**) Median JAM:JAM adhesion size is determined as the median area of JAM-SNAP segmented objects that overlap with JAM-C-pH segmented signal. Data points are reported as individual biological replicates statistically compared using Wilcoxon Rank Sum Tests. Statistical significance is represented by standard convention: ns p>0.05, * p<0.05, ** p<0.01, *** p<0.001, and **** p<0.0001.

**Figure 5 – figure supplement 1.**
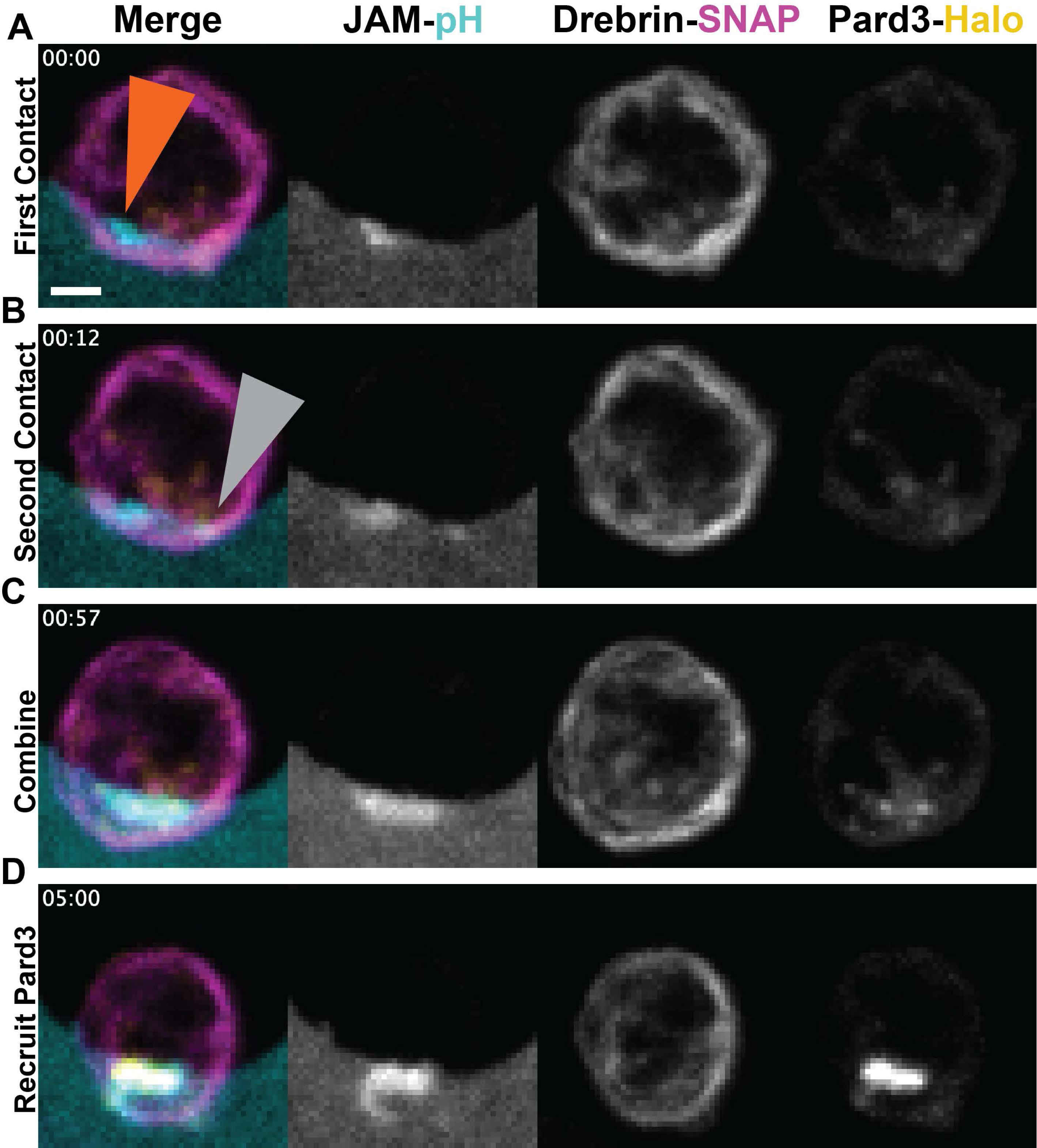
JAM-C recognition of glial JAM-B recruits Pard3 in seconds and drebrin in minutes. (**A-D**) Representative max projection images of JAM-pH, drebrin-SNAP, and Halo-Pard3 during JAM-C WT contact formation to JAM-B in glia acquired at 3 second intervals. Data is presented with individual channels represented in grayscale and merged data presented in color. (**A**) First sign of contact between the CGN and glia as a proportionally bright area of JAM-pH marked with orange arrowhead. (**B**) A second JAM contact forms after 12 seconds that directly overlaps with Halo-Pard3 marked by a gray arrowhead. (**C**) The two initial contacts fuse at 57 seconds with Halo-Pard3 coincidence. (**D**) At 5 minutes, more Halo-Pard3 has been recruited to the contact site and drebrin-SNAP is excluded to the contact periphery.

**Figure 5 – figure supplement 2.**
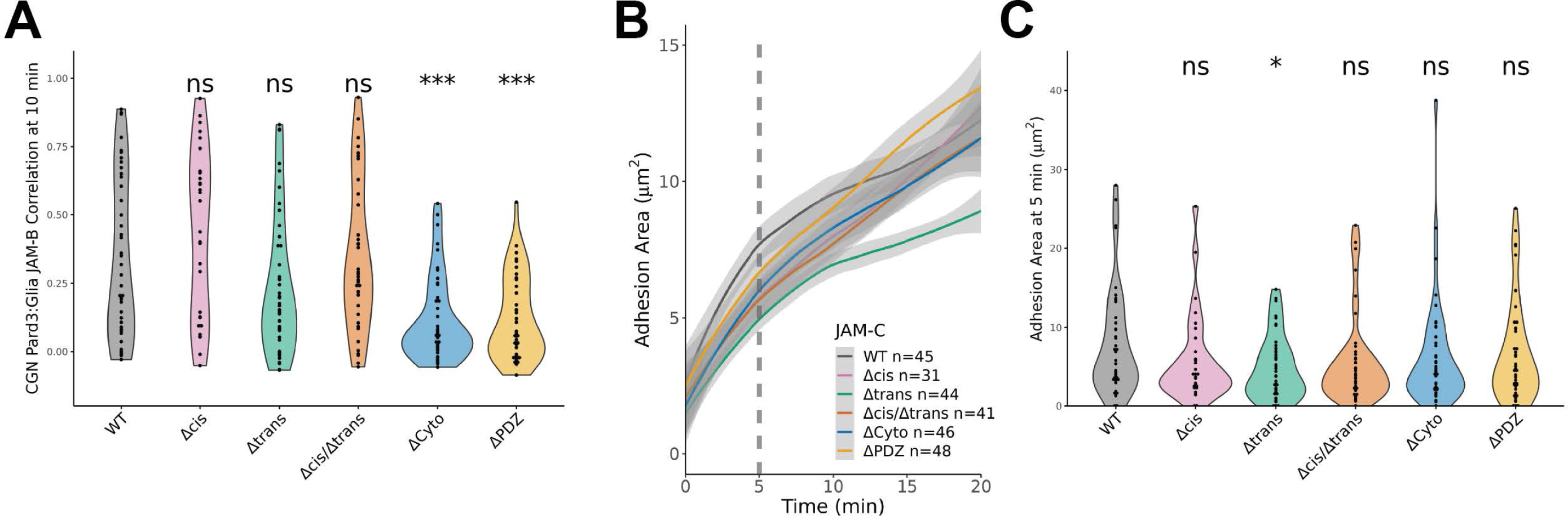
Δtrans JAM-C delays Pard3 recruitment and adhesion formation. (**A**) Statistical comparison of Pearson correlation between JAM-pH and Halo-Pard3 at 10 minutes after first sign of adhesion. The Pearson correlation of each technical replicate is plotted within the violin plot. Pairwise statistical significance is calculated in relation to JAM-C WT replaced CGNs using Wilcoxon Rank Sum Tests. (**B**) Loess smoothed plot of adhesion size as a function of time after first sign of adhesion is shown. The dashed line at 5 minutes indicates the time point for subsequent statistical comparison. (**C**) Statistical comparison of adhesion size at 5 minutes after first sign of adhesion. The adhesion size of each technical replicate is plotted within the violin plot. The plots and statistical comparisons were generated for the combined technical replicates (n) from at least 3 independent biological replicates (N≥3) uniformly subsampled at 30 second intervals. Pairwise statistical significance is calculated in relation to JAM-C WT replaced CGNs using Wilcoxon Rank Sum Tests. Gray shading on the plots represents the 95% confidence intervals of the estimated mean values. Statistical significance is represented by standard convention: ns p>0.05, * p<0.05, ** p<0.01, *** p<0.001, and **** p<0.0001.

**Figure 5 – figure supplement 3.**
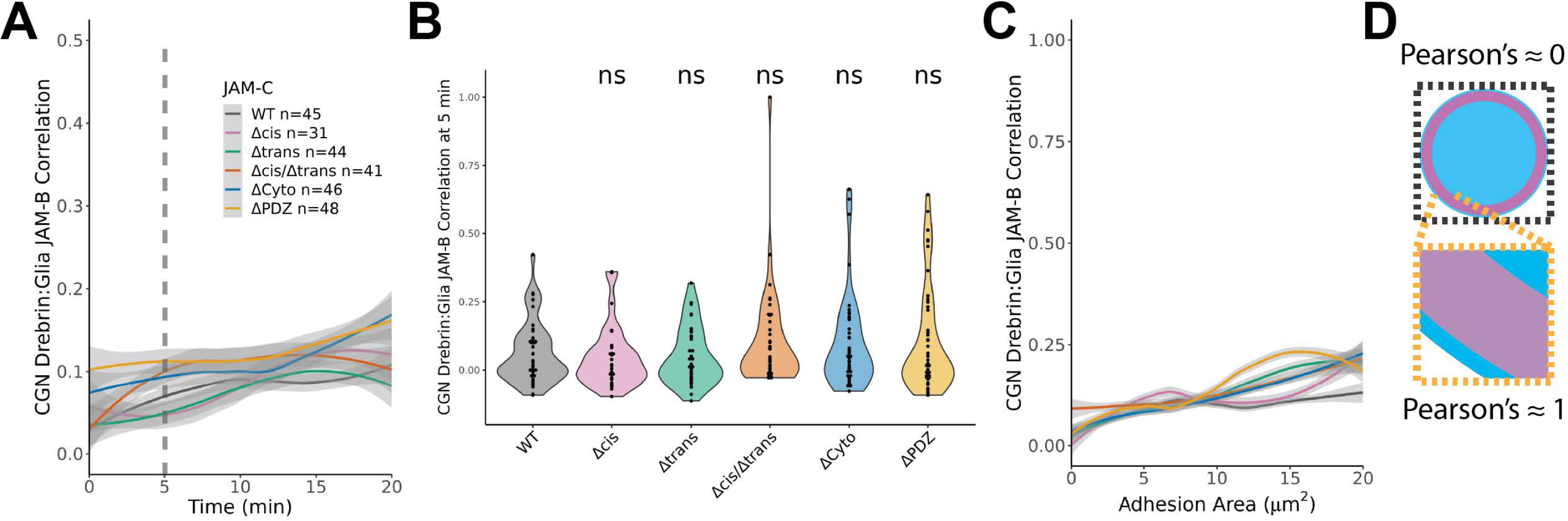
JAM contact formation on glia does not directly recruit drebrin within the contact site. (**A**) Loess smoothed plot of drebrin-SNAP recruitment to glial JAM contacts measured by Pearson correlation for individual cells as a function of the time after first adhesion. The dashed line indicates the point in time used for statistical comparison. (**B**) Statistical comparison of Pearson correlation between JAM-pH and drebrin-SNAP at 5 minutes after first sign of adhesion. The Pearson correlation of each technical replicate is plotted within the violin plot. Pairwise statistical significance is calculated in relation to JAM-C WT replaced CGNs using Wilcoxon Rank Sum Tests. (**C**) Loess smoothed plot of drebrin-SNAP recruitment to glial JAM contacts measured by Pearson correlation for individual cells as a function of the segmented adhesion size. (**D**) Diagrammatic explanation of low Pearson correlation of the JAM-pH contact and drebrin. Since drebrin-SNAP borders the contact, regions of correlation and anticorrelation yield a correlation of 0. When zoomed in at the contact edge, the Pearson Correlation will trend towards 1. Plots and statistical comparisons were generated for the combined technical replicates (n) from at least 3 independent biological replicates (N≥3) uniformly subsampled at 30 second intervals. Gray shading on the plots represents the 95% confidence intervals of the estimated mean values. Statistical significance is represented by standard convention: ns p>0.05, * p<0.05, ** p<0.01, *** p<0.001, and **** p<0.0001.

**Figure 6 – figure supplement 1.**
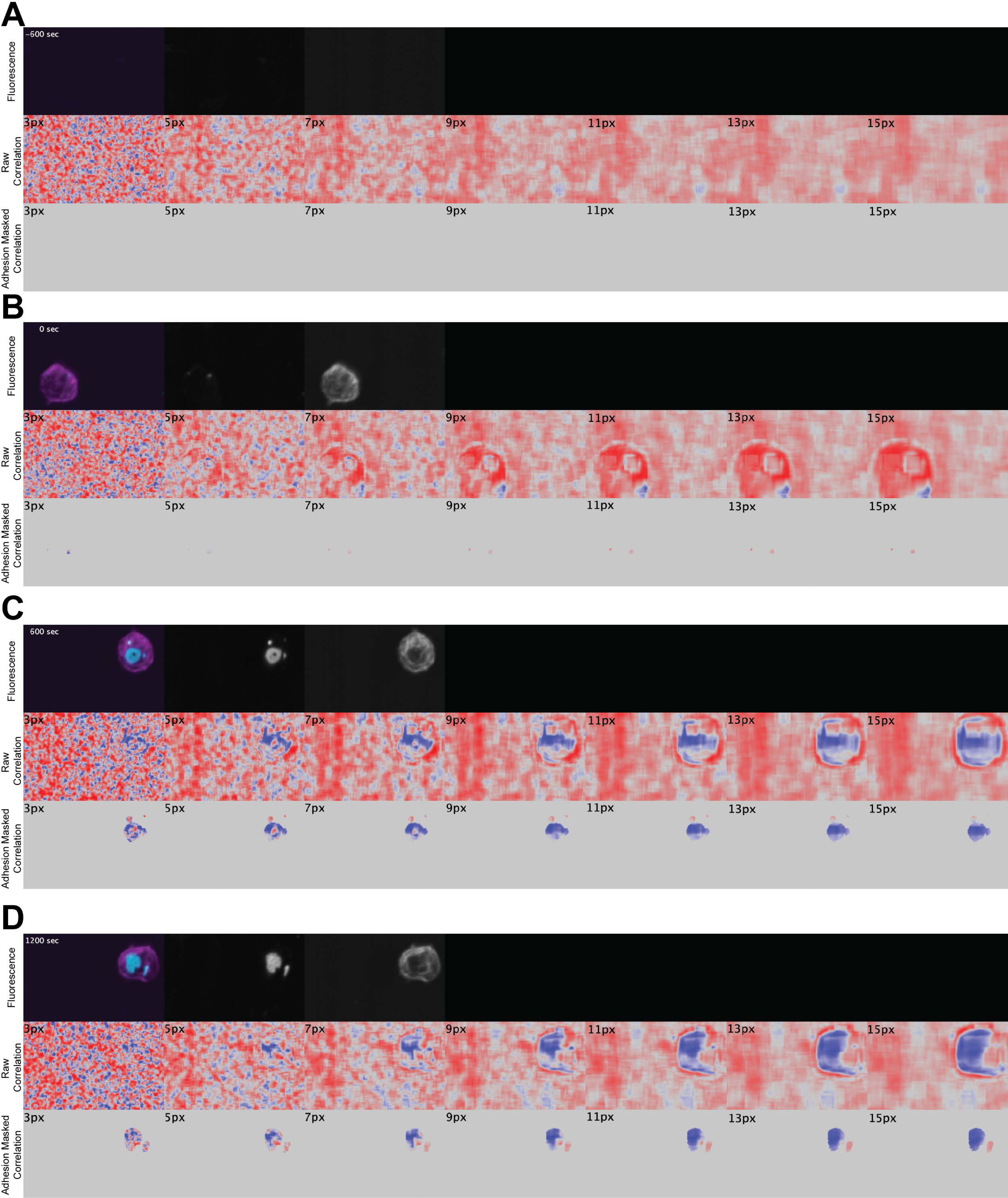
Sensitivity analysis of correlation kernel indicates 9 pixels strikes a balance between sensitivity and noise. (**A-D**) Representative time points in the timelapse data of the local correlation of JAM:drebrin used for empirical selection of kernel size. The first row in in each image shows the fluorescence of individual channels in grayscale and merge in color. The second row shows the raw data output of local correlation analysis at varied pixel window sizes indicated in the top left corner ranging from 3-15 pixels. The third row applies a manually determined segmentation mask to the raw correlation to restrict our analyses to the JAM contact as our feature of interest. The time frames included represent (**A**) prior to adhesion, (**B**) first sign of adhesion, (**C**) adhesion formation, and (**D**) stable adhesion.

**Figure 6 – figure supplement 2.**
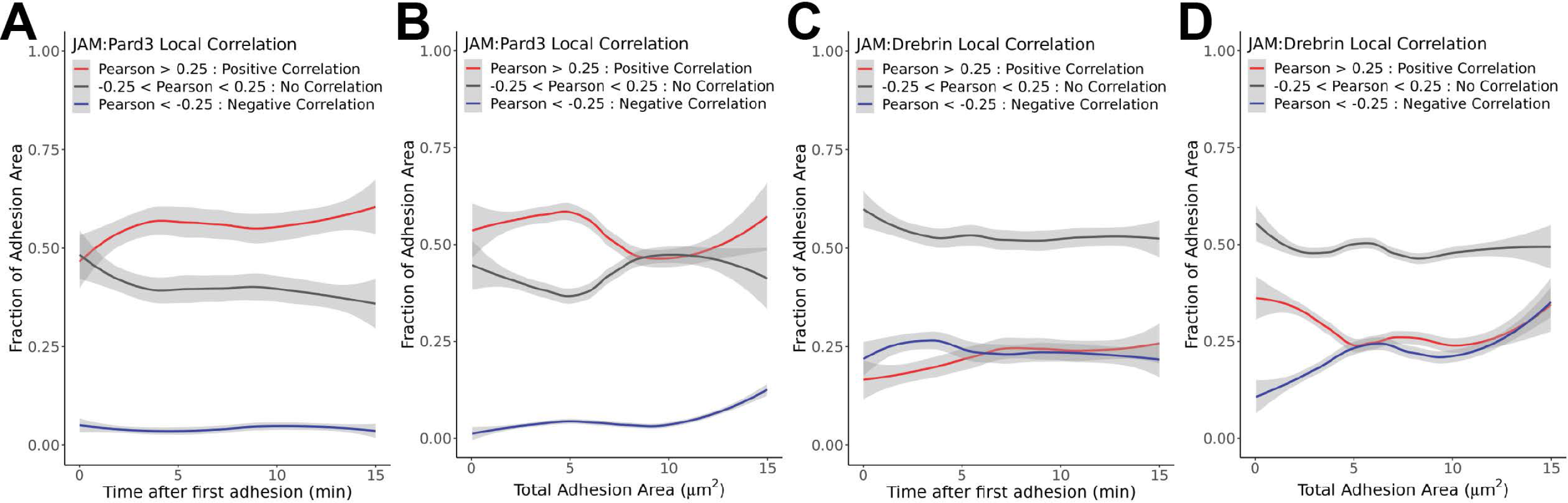
Local correlation of JAM contacts reveals JAM:Pard3 correlation is mostly positive whereas JAM:drebrin has regions of positive and negative correaltion. (**A-D**) Data of the proportional makeup of local correlation at WT JAM-C adhesions to glial JAM-B for all technical replicates (n=44) uniformly subsampled at 30 second intervals. Loess smoothed plots of the proportions of positive correlation (Pearson > 0.25), no correlation (-0.25<Pearson<0.25), or negative correlation of JAM:Pard3 as a function of (**A**) time after first adhesion and (**B**) adhesion area. Loess smoothed plots of the proportions of positive correlation, no correlation, or negative correlation of JAM:drebrin as a function of (**C**) time after first adhesion and (**D**) adhesion area. Gray shading on the plots represents the 95% confidence intervals of the estimated mean values.

**Figure 6 – figure supplement 3.**
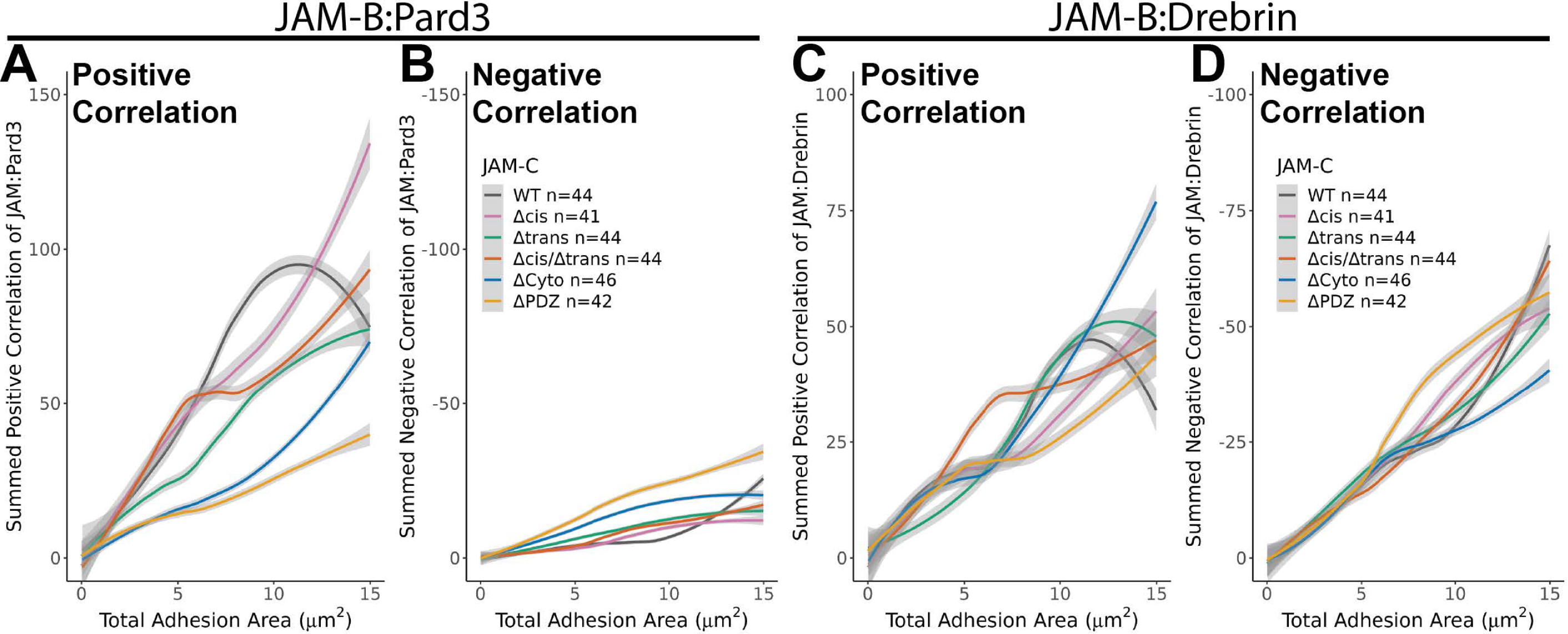
JAM:Pard3 and JAM:drebrin sum correlation is associated with increasing JAM adhesion size. (**A-D**) Data of the local recruitment of drebrin and Pard3 to JAM contacts based on JAM adhesion size. Loess smoothed plots of the sum of the (**A**) positive (Pearson > 0) and (**B**) negative (Pearson<0) JAM:Pard3 local correlation as a function of JAM adhesion area. Loess smoothed plots of the sum of the (**C**) positive (Pearson > 0) and (**D**) negative (Pearson<0) JAM:drebrin local correlation as a function of JAM adhesion area. The plots of sum correlation were generated from the combination of technical replicates (n) from at least 3 independent biological replicates (N≥3) uniformly subsampled at 30 second intervals. Gray shading on the plots represents the 95% confidence intervals of the estimated mean values.

**Figure 6 – figure supplement 4.**
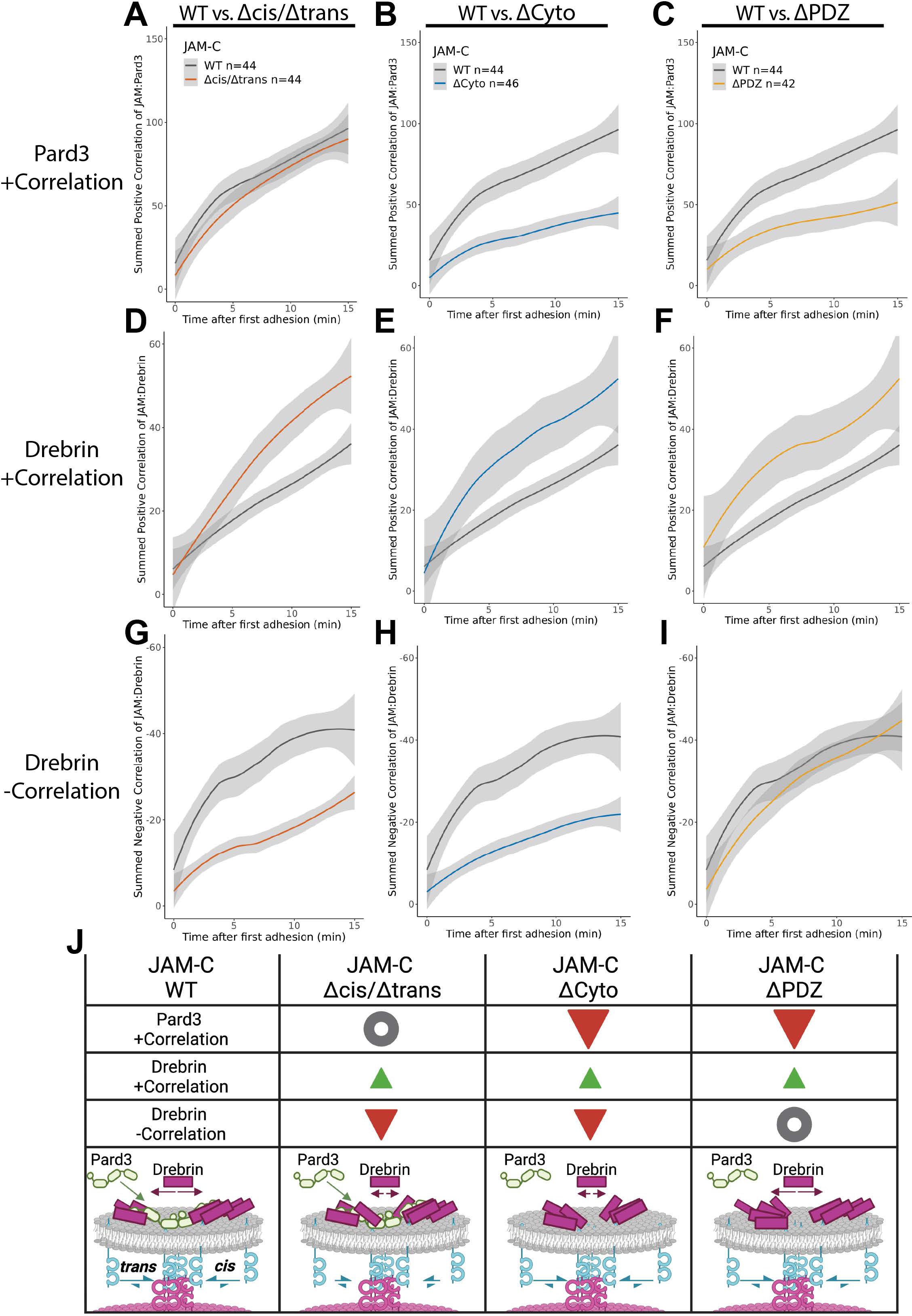
Paired comparisons of sum local correlation highlight the role of intracellular interactions and JAM adhesion. (**A-C**) Loess smoothed curves of the sum of positive JAM:Pard3 correlation as a function of time compared to JAM-C WT for (**A**) JAM-C Δcis/Δtrans, (**B**) ΔCyto, and (**C**) ΔPDZ. (**D-F**) Loess smoothed curves of the sum of positive JAM:drebrin correlation as a function of time compared to JAM-C WT for (**D**) JAM-C Δcis/Δtrans, (**E**) ΔCyto, and (**F**) ΔPDZ. (**G-I**) Loess smoothed curves of the sum of negative JAM:drebrin correlation as a function of time compared to JAM-C WT for (**G**) JAM-C Δcis/Δtrans, (**H**) ΔCyto, and (**I**) ΔPDZ. Loess smoothed plots were generated from the data of all technical replicates (n) as shown in plot legends. (**J**) Diagrammatic summary of correlation data. Gray circles indicate no change from WT JAM-C, red inverted triangles indicate decrease relative to WT JAM-C, and green triangles indicate increased value relative to WT JAM-C. Diagrams in the bottom row propose the molecular arrangement associated with each JAM construct. Gray shading on the plots represents the 95% confidence intervals of the estimated mean values.

**Figure 6 – figure supplement 5.**
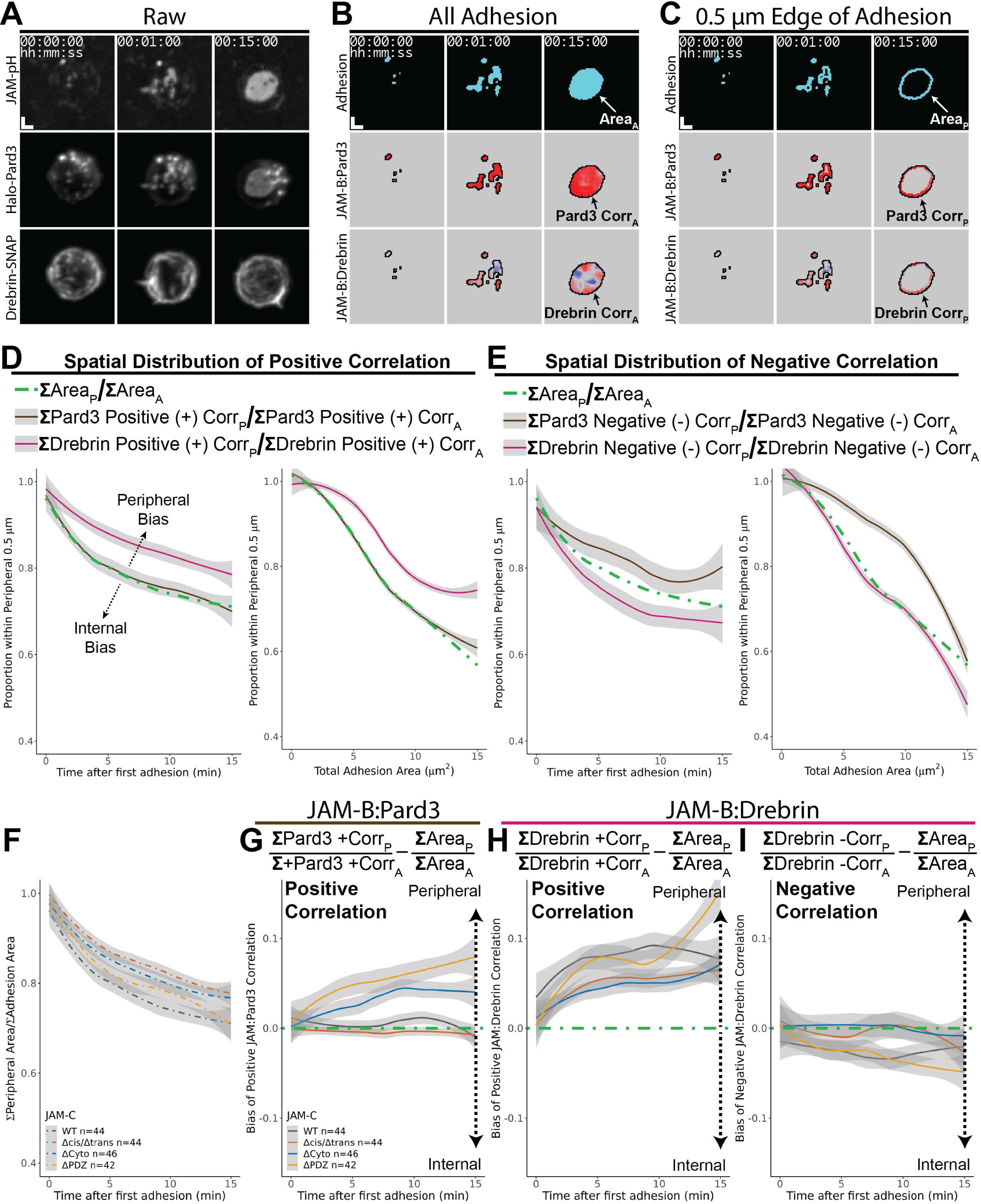
Local correlation of JAM:drebrin but not JAM:Pard3 is spatially distribution to the outer 0.5 µm of JAM contacts. (**A-C**) Example images used to calculate the spatial distribution of local correlation are shown. (**A**) Representative grayscale fluorescence images gamma adjusted by 0.7 at the first sign of adhesion, 1 minute after first adhesion, and 15 minutes after first adhesion. (**B**) Data associated with the fluorescent images in (**A**) showing the segmentation mask of JAM adhesion with the associated correlation maps for JAM:Pard3 and JAM:drebrin denoted by the subscript ‘A’. (**C**) Data associated with only the 0.5 µm periphery of the JAM adhesion in (**B**) denoted by the subscript ‘P’. (**D-G**) The deviation of proportional correlation from adhesion area reveals spatial distribution of Pard3 and drebrin for WT JAM-C adhering to glial JAM-B (n=44). Loess smoothed plots of the proportion of positive correlation or adhesion area within the 0.5 µm periphery of the JAM contact divided by the entire adhesion are shown as a function of (**D**) time after first adhesion or (**E**) adhesion size. Loess smoothed plots of the proportion of negative correlation or adhesion area within the 0.5 µm periphery for the JAM contact divided by the entire adhesion are shown as a function of (**D**) time after first adhesion or (**E**) adhesion size. (**F-I**) JAM-C mutants exhibit distribution of JAM:Pard3 and JAM:drebrin local correlation from the combined data for all technical replicates (n) indicated in associated plot legends. (**F**) Loess smoothed plot of the ratio of the sum adhesion area within 0.5 µm of adhesion edge divided by total adhesion area for JAM-C constructs to glial JAM-B is shown. (**G-I**) Plots show the associated deviation from peripheral adhesion area by subtracting the ratio from the proportional sum correlation. (**G**) Loess smoothed plot of the spatial distribution of JAM:Pard3 sum positive correlation as a function of time after first adhesion is shown. Loess smoothed plots of the spatial distribution are shown for JAM:drebrin (**H**) sum positive correlation and (**I**) sum negative correlation as a function of time after first adhesion. Gray shading on the plots represents the 95% confidence intervals of the estimated mean values.

**Figure 7 – figure supplement 1.**
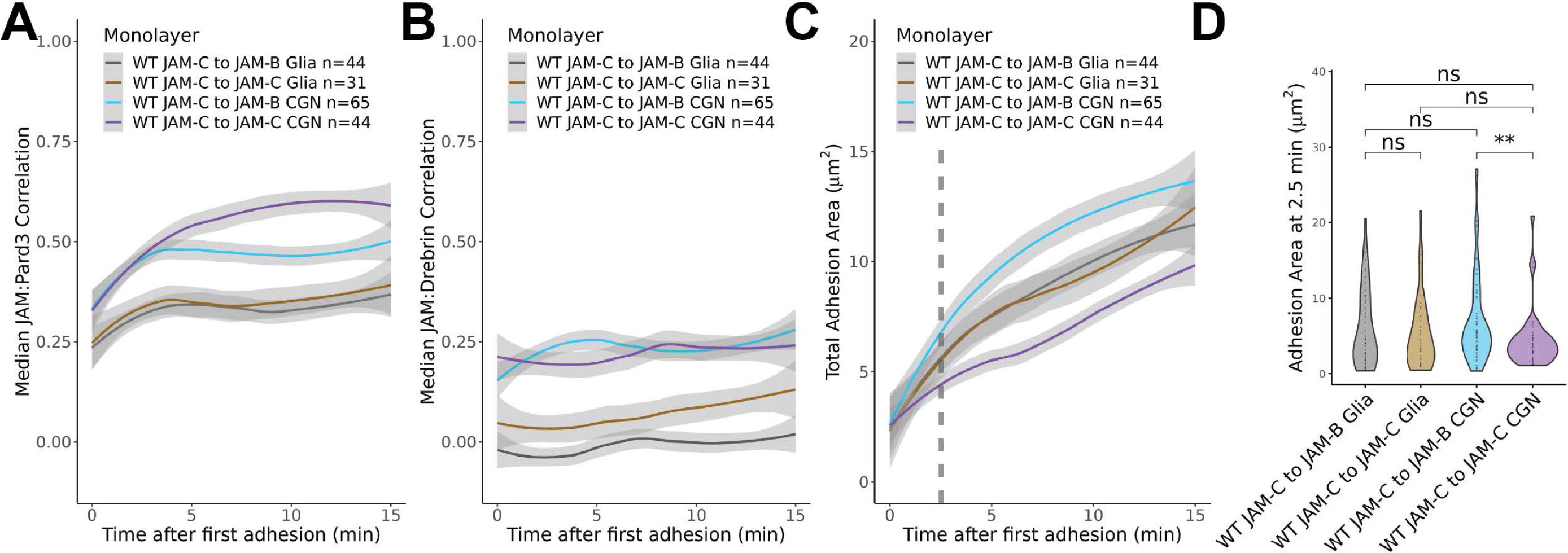
CGN monolayers are preferential to glia for drebrin and Pard3 recruitment to JAM contacts even though JAM-C affects CGN-CGN adhesion size. (**A-B**) Loess smoothed plots of the median correlation within segmented adhesions are shown for (**A**) JAM:Pard3 and (**B**) JAM:Drebrin as a function of time after first adhesion for WT JAM-C expressing CGNs adhering to monolayers of glia or CGNs expressing either JAM-C or JAM-B. (**C**) Loess smoothed plot of adhesion area are shown for WT JAM-C expressing CGNs adhering to monolayers of glia or CGNs expressing either JAM-C or JAM-B. (**D**) Statistical comparison of JAM adhesion size at 2.5 minutes after first adhesion. Pairwise statistical comparisons were performed as Wilcoxon rank sum comparisons in relation to either WT JAM-C to JAM-C CGN adhesion or WT JAM-C to JAM-B glia adhesion as indicated by the brackets. Loess plots and statistical comparisons were generated from the combination of all technical replicates (n) across at least 3 biological replicates replicates (N≥3). Gray shading on the plots represents the 95% confidence intervals of the estimated mean values. Statistical significance is represented by standard convention: ns p>0.05, * p<0.05, ** p<0.01, *** p<0.001, and **** p<0.0001.

**Figure 7 – figure supplement 2.**
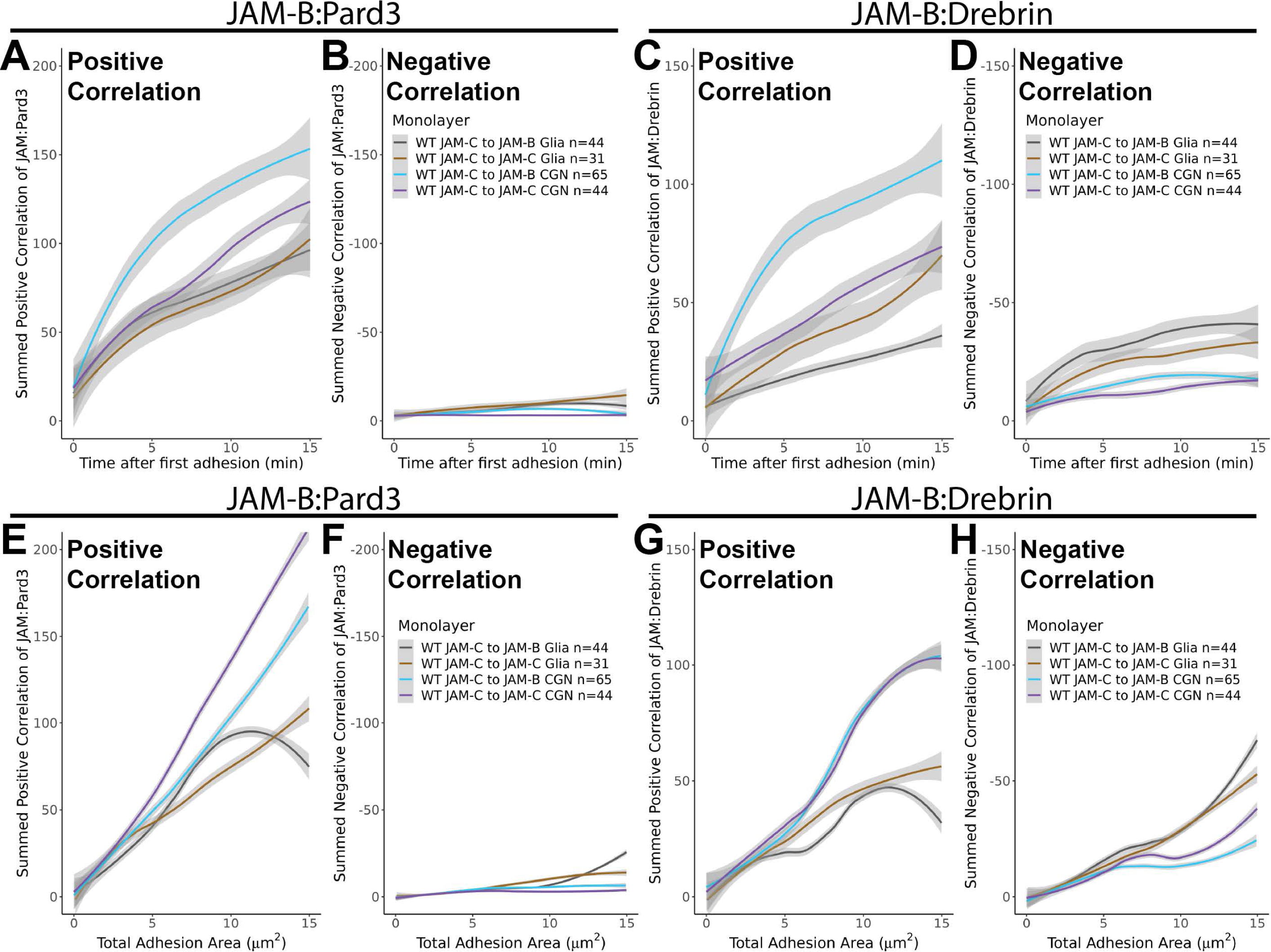
Recruitment of Pard3 and drebrin is mostly driven by cell substrate rather than JAM receptor. (**A-D**) Comparison of the sum correlation of JAM:Pard3 and JAM:drebrin for JAM-C expressing CGNs adhering to JAM-B or JAM-C on glia or CGNs as a function of time after first adhesion. Loess smoothed plots of JAM:Pard3 (**A**) sum positive correlation and (**B**) sum negative correlation. Loess smoothed plots of JAM:drebrin (**C**) sum positive correlation and (**D**) sum negative correlation. (**E-H**) Comparison of the sum correlation of JAM:Pard3 and JAM:drebrin for JAM-C expressing CGNs adhering to JAM-B or JAM-C on glia or CGNs as a function of adhesion area. Loess smoothed plots of JAM:Pard3 are shown for (**E**) sum positive correlation and (**F**) sum negative correlation. Loess smoothed plots of JAM:drebrin are shown for (**G**) sum positive correlation and (**H**) sum negative correlation. Loess plots were generated from the combination of all technical replicates (n) indicated in the plot legends. Gray shading on the plots represents 95% confidence intervals of the estimated mean values.

**Figure 7 – figure supplement 3.**
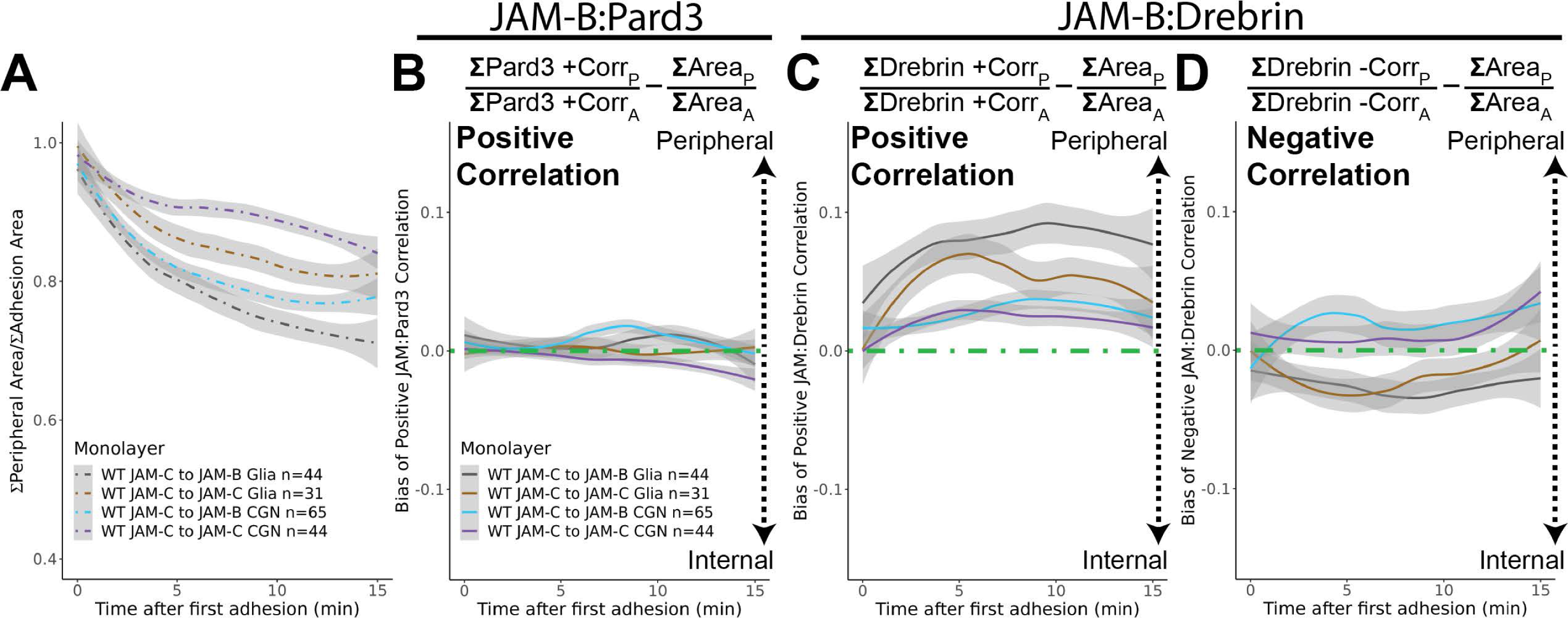
JAM contacts on glia elicit a stronger peripheral spatial distribution for drebrin than CGNs but not for Pard3. (**A**) Loess smoothed plot of the ratio of the sum adhesion area within 0.5 µm of adhesion edge divided by total adhesion area for WT JAM-C expressing CGNs adhering to JAM-C or JAM-B on glia or CGN monolayers. (**B-D**) Plots show the associated deviation from peripheral adhesion area by subtracting the ratio from the proportional sum correlation. (**B**) Loess smoothed plot of the spatial distribution of JAM:Pard3 sum positive correlation as a function of time after first adhesion is shown for WT JAM-C expressing CGNs added to types of monolayers specified in plot legends. Loess smoothed plots of the spatial distribution are shown for JAM:drebrin (**H**) sum positive correlation and (**I**) sum negative correlation as a function of time after first adhesion is shown for WT JAM-C expressing CGNs added to the types of monolayers specified in plot legends. Gray shading on the plots represents the 95% confidence intervals of the estimated mean values.

**Figure 7 – figure supplement 4.**
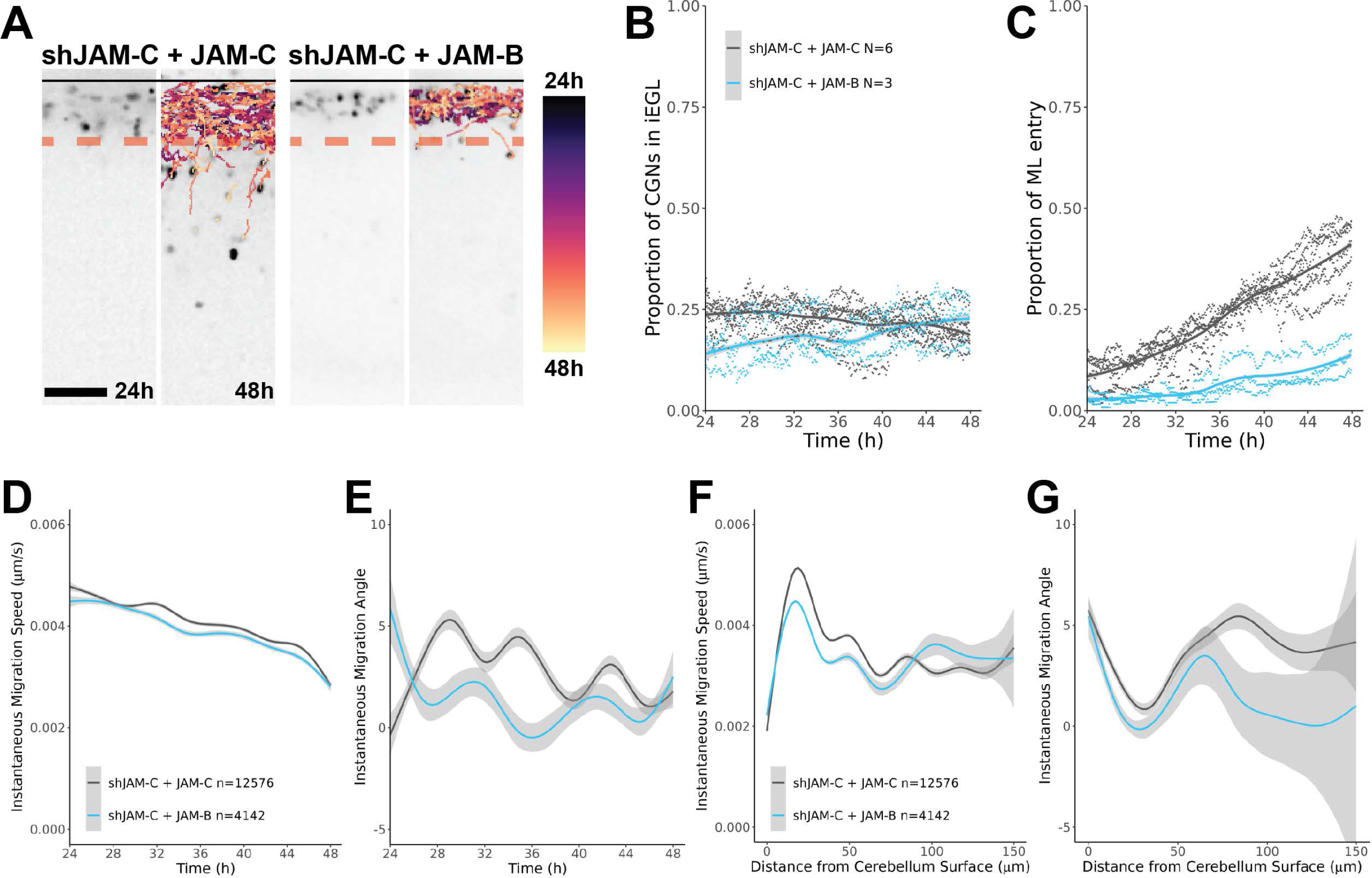
JAM-B replacement in *ex vivo* slice cultures delays CGN iEGL entry and halts ML entry. (**A**) Representative images at 24 and 48 hours of CGNs tracked in *ex vivo* slice cultures by live imaging slices electroporated to express either shJAM-C and WT JAM-C or shJAM-C and WT JAM-B. Time position of the tracked CGNs is color encoded as shown by the provided lookup table. (**B-C**) Plots of the layer occupancy of CGNs are shown from the proportions calculated for individual biological replicates (N) are shown. (**B**) Loess smoothed plot of the proportion of CGNs in the iEGL as a function of time is shown. (**C**) Loess smoothed plot of the proportion of CGNs that have entered the ML as a function of time is shown. (**D-G**) Plots of migration data for all cell migration events (n) aggregated across all the biological replicates (N) are shown. Loess smoothed plots of (**D**) migration speed and (**E**) instantaneous migration angle relative to the cerebellar surface as a function of time are shown. Loess smoothed plots of (**F**) migration speed and (**G**) instantaneous migration angle relative to the cerebellar surface distance from the cerebellar surface are shown. Gray shading on the plots represents the 95% confidence intervals of the estimated mean values. Scale bar = 50 µm.

